# Morphological Reprogramming of Primary Cilia Length Mitigates the Fibrotic Phenotype in Fibroblasts Across Diverse Fibrotic Conditions

**DOI:** 10.1101/2024.01.06.574461

**Authors:** Priyanka Verma, Bharat Yalavarthi, Swati Bhattacharyya, Dinesh Khanna, Johann E. Gudjonsson, Lam C. Tsoi, Rebecca Wells, Rebecca L Ross, Natalia Riobo-Del Galdo, Francesco Del Galdo, Sean M. Fortier, Maria E. Teves, John Varga, Dibyendu Bhattacharyya

## Abstract

Fibrosis is a hallmark of systemic sclerosis (SSc) and many diverse and incurable diseases. Myofibroblast activation, a common cellular phenomenon shared across fibrotic diseases, is marked by actin polymerization known to affect primary cilia (PC) length. We discovered that fibroblasts from diverse fibrotic conditions display significantly reduced PC length *ex vivo.* Treatment of healthy fibroblasts with profibrotic TGF-β1 induced PC shortening, while silencing *ACTA2* in SSc skin fibroblasts caused PC elongation. Importantly, we found that PC length is negatively correlated with cellular expression of α-SMA in TGF-β1-stimulated healthy fibroblasts, or pharmacologically de-differentiated myofibroblasts. PC elongation by microtubule polymerization induction in SSc skin fibroblasts using LiCl or the HDAC6 inhibitor tubacin, reversed and mitigated fibrotic responses. Our results implicate a key role for microtubule polymerization in restraining fibrotic responses and suggest that modulation of PC dynamics may represent a potential therapeutic strategy for SSc and other treatment-resistant diseases associated with fibrosis.

**Teaser.** PC length shortening is a hallmark of fibrosis.

## INTRODUCTION

Fibrotic disorders represent a leading cause of mortality worldwide (*1*). Systemic sclerosis (SSc), a heterogeneous chronic systemic disease, is dominated by fibrosis affecting multiple tissues synchronously (*2, 3*). A subset of SSc patients develop fibrotic interstitial lung disease and other life-threatening organ involvement (*4*). Despite advances during the past decade, safe and effective disease-modifying therapies for SSc are still lacking, and a cure remains elusive (*5, 6*). Systemic sclerosis is one of a burgeoning group of both common and rare conditions complicated by fibrosis (*3*). These include idiopathic pulmonary fibrosis (IPF), characterized by scarring and persistent fibroblast activation in the lung (*7*). Fibrosis of the orbits is the hallmark of thyroid-associated ophthalmopathy (TAO), which currently has limited treatment options (*8*).

The diversity across fibrotic diseases presents a challenge for designing broad-spectrum anti-fibrotic drugs. Even within a specific fibrotic disorder, such as SSc, there is substantial phenotypic diversity among fibroblasts from affected tissues (*9*). Nevertheless, myofibroblast differentiation is recognized as a common phenomenon that is shared among distinct fibrotic diseases and affected organs (*10*). While the fundamental role played by morphogens such as Transforming Growth Factor-β (TGF-β), Wnt, and Sonic Hedgehog (SHH) in pathological myofibroblast differentiation in SSc and other fibrotic diseases is established, the precise mechanisms underlying pathological myofibroblast transition and sustained activation in different conditions remain poorly understood (*11*).

Myofibroblast activation is marked by higher-order of actin polymerization, including formation of F-actin stress fibers and inclusion of alpha-smooth muscle actin (αSMA) into a stress fiber network, which in turn contributes to the contractile properties of myofibroblast (*12*). Primary cilia (PC) are solitary membrane organelles present in virtually all non-dividing vertebrate cells. Previous studies indicated that polymerization of the actin cytoskeleton, F-actin branching, and stress fiber formation each inhibited the formation of PC; while on the other hand, actin depolymerization or depletion promoted PC elongation (*13, 14*). We hypothesized that PC morphology might be impacted in all fibrotic disorders, and such a phenotype might be exploited to design broad-spectrum anti-fibrotic drugs.

Primary or non-motile cilia play fundamental roles in cell signaling (*15, 16*). Acting as sensory cellular antennae for chemical and biomechanical signals, PC exert a profound influence on tissue repair and regeneration (*17*). Remarkably, PC host key components of multiple morphogen signaling pathways implicated in myofibroblast transition (*18*). Downregulation of PC genes has been associated with SSc pathogenesis(*19, 20*). Moreover, dysfunction of PC has been implicated in several forms of fibrosis (*18*). While increased PC length and expression of cilia-related genes has been demonstrated in fibroblasts in cardiac and pulmonary fibrosis (*21–23*), a group of conditions called ciliopathies characterized by abnormal PC formation and function exhibit pronounced myofibroblast accumulation and fibrosis in affected tissues (*24–27*). Furthermore, ciliary loss in fibroblasts induced by disruption of the PC gene Ift88 is accompanied by myofibroblast transition (*28*). Collectively, these observations suggest a functional link between PC and myofibroblasts(*18, 29*), but also highlight cell-type-specific and potentially biphasic mechanisms in myofibroblast transition and fibrosis (*30*). Despite these observations from disease models, the expression and function of PC in fibrosis, their potential role in pathogenesis, and their therapeutic targeting remain largely unknown.

In the present study, we demonstrate that PC length is reduced in fibroblasts explanted from three distinct fibrotic conditions compared to their matched healthy controls and is negatively associated with myofibroblast activation. Furthermore, the profibrotic signal TGF-β elicits PC shortening in fibroblasts, while dedifferentiation of myofibroblasts to quiescence, or genetic deletion of αSMA, is accompanied by elongation of PC in parallel with attenuation of the fibrotic phenotype. Using a variety of pharmacological agents to positively or negatively modulate *ex vivo* ciliogenesis in fibroblasts, we observed a consistent reciprocal inverse association between PC length and fibrotic phenotypes. Altogether, these results uncover novel mechanistic links between the length of PC in fibroblasts and their fibrotic phenotype across distinct fibrotic conditions. We propose that pharmacological modulation of PC dynamics represents a potential therapeutic approach for SSc, and other currently incurable fibrotic diseases.

## RESULTS

### Fibroblasts from diverse fibrotic conditions and across organs display significantly reduced PC length *ex vivo*

Myofibroblast activation, a common cellular phenomenon shared across all fibrotic diseases, is marked by higher-order of actin polymerization, F-actin stress fiber formation, and incorporation of αSMA, providing increased contractility (*12, 14*). Previous studies suggested that higher contractile force and polymerization of the actin cytoskeleton negatively regulate ciliogenesis (*13, 14, 31–33*). Based on these observations, we hypothesized that PC length will be reduced in fibrotic fibroblasts irrespective of their disease or tissue origin, primarily due to their higher-order actin polymerization underlying myofibroblast transition.

To examine this hypothesis, we compared PC length in fibroblasts explanted from affected organs (skin, orbit, lung) from three fibrotic disorders and their matched healthy controls(Table-S1). Fibroblasts were cultured under conditions that promote PC growth for 24 h, and then immunolabelled using antibodies to Arl13b, a marker for PC membrane (*34*). We observed that SSc skin fibroblasts *in vitro* display significantly shorter PC, compared to matched healthy skin fibroblasts (Fig. 1A, D, E). Furthermore, analyzing the PC length in fibroblasts from healthy controls and Very Early Diagnosis of Systemic Sclerosis (VEDOSS) indicated similar PC length shortening as observed in SSc fibroblasts (Suppl. Fig. S2). Patients with VEDOSS have no clinical skin involvement, but are at heightened risk for developing SSc within 5 years but do show an increased profibrotic phenotype compared to healthy controls (*35*). Fibroblasts explanted from IPF lungs (Fig. 1C, D) or TAO orbits, similarly display significantly shortened PC compared to their healthy matched controls (Fig. 1B, D).

**Figure 1.**
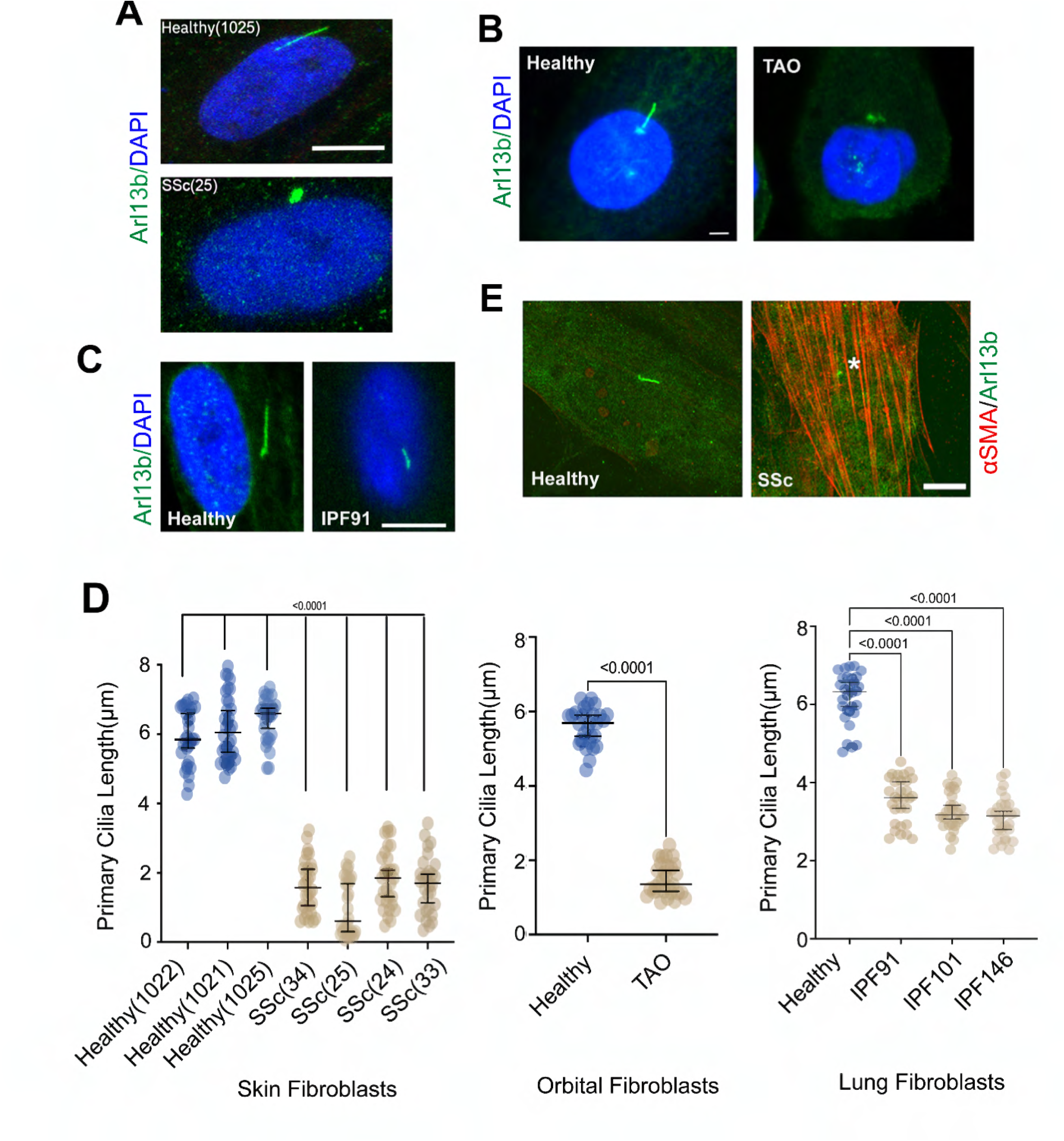
Fibrotic fibroblasts display reduced primary cilia length. Healthy and patient-derived fibroblasts (**A, D**: Healthy (n=3) and SSc skin fibroblasts (n=4); **B,D**: Healthy and thyroid-associated ophthalmopathy (TAO) orbital fibroblasts (n=2 in each group); **C,D**: Healthy (n=3 independent experiment) and IPF (n=3) lung fibroblasts at low passage were incubated in media with 1% FBS. After 16 h, fibroblasts were fixed and immunolabelled using antibodies against Arl13b (green) **(A)**, Arl13b (green) **(B, C)** or Arl13b (green) and αSMA (red) **(E)**. Blue represents DAPI. Scale bar 5 μm **(A, B, C)** or 10 μm **(E)**. **D.** Cells were imaged with a Leica SP8 confocal microscope with z-volume, and PC lengths were measured from the 3D reconstruction of the Z-volumes using ImageJ (mean ± SEM from 30 determinations). Each dot represents the PC length from a different cell.

### Treatment with TGF-β1 induces PC shortening in healthy fibroblasts

TGF-β1 drives myofibroblast transition in fibroblasts (*18*). To determine the impact of TGF-β1 on PC, we incubated serum-starved skin or lung fibroblasts for 24 h, followed by immunolabelling using antibodies to Arl13b and αSMA. Treatment of healthy skin and lung fibroblasts with TGF-β1 was associated with a time-and dose-dependent reduction in PC length, with concurrent increase in αSMA levels and incorporation into stress fibers (Fig. 2A-D, Suppl. Fig. S1).

**Figure 2.**
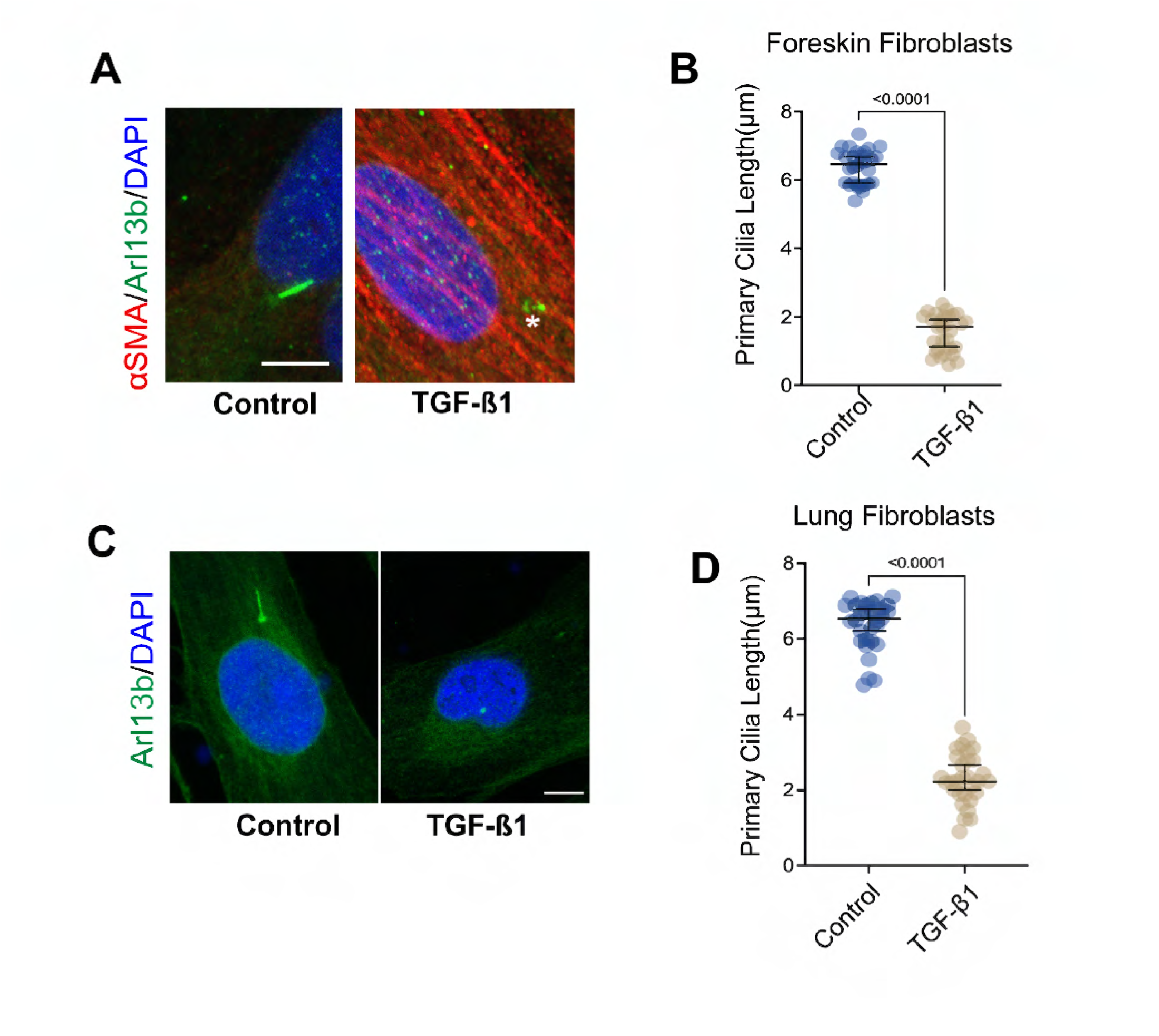
TGFβ1-induced primary cilia shortening in foreskin and lung fibroblasts is associated with myofibroblast differentiation. **(A)** Foreskin (A) or CCL210 lung (**C**) fibroblasts were incubated without or with TGF-β1 (10 ng/ml) for 24 h in media with 0.1%BSA, then fixed and immunolabelled using antibodies against Arl13b (green) and αSMA (red). Representative images. Scale bar 10 μm. **(B, D)** Quantification of PC length (mean ± SEM from 20 determinations). PC lengths were measured from the 3D reconstruction of the Z-volumes using ImageJ.

### Primary cilia length is restored in dedifferentiated fibrotic lung fibroblasts

Pharmacological agents can drive dedifferentiation of TGF-β1-stimulated lung myofibroblasts, with down-regulation of αSMA (*36*). To investigate how myofibroblast dedifferentiation impacts PC, TGF-β1-treated CCL210 lung fibroblasts were incubated with PGE_2_ or FGF2, as described previously (*36*). PGE_2_ and FGF2-induced dedifferentiation of activated lung fibroblasts was accompanied by elongation of PC (Fig. 3A, B). Notably, PC length in these fibroblasts was negatively associated with their αSMA levels (Fig. 3C). In contrast to lung myofibroblasts, treatment of activated skin fibroblasts with PGE_2_ and FGF2 failed to induce their dedifferentiation (data not shown).

**Figure 3:**
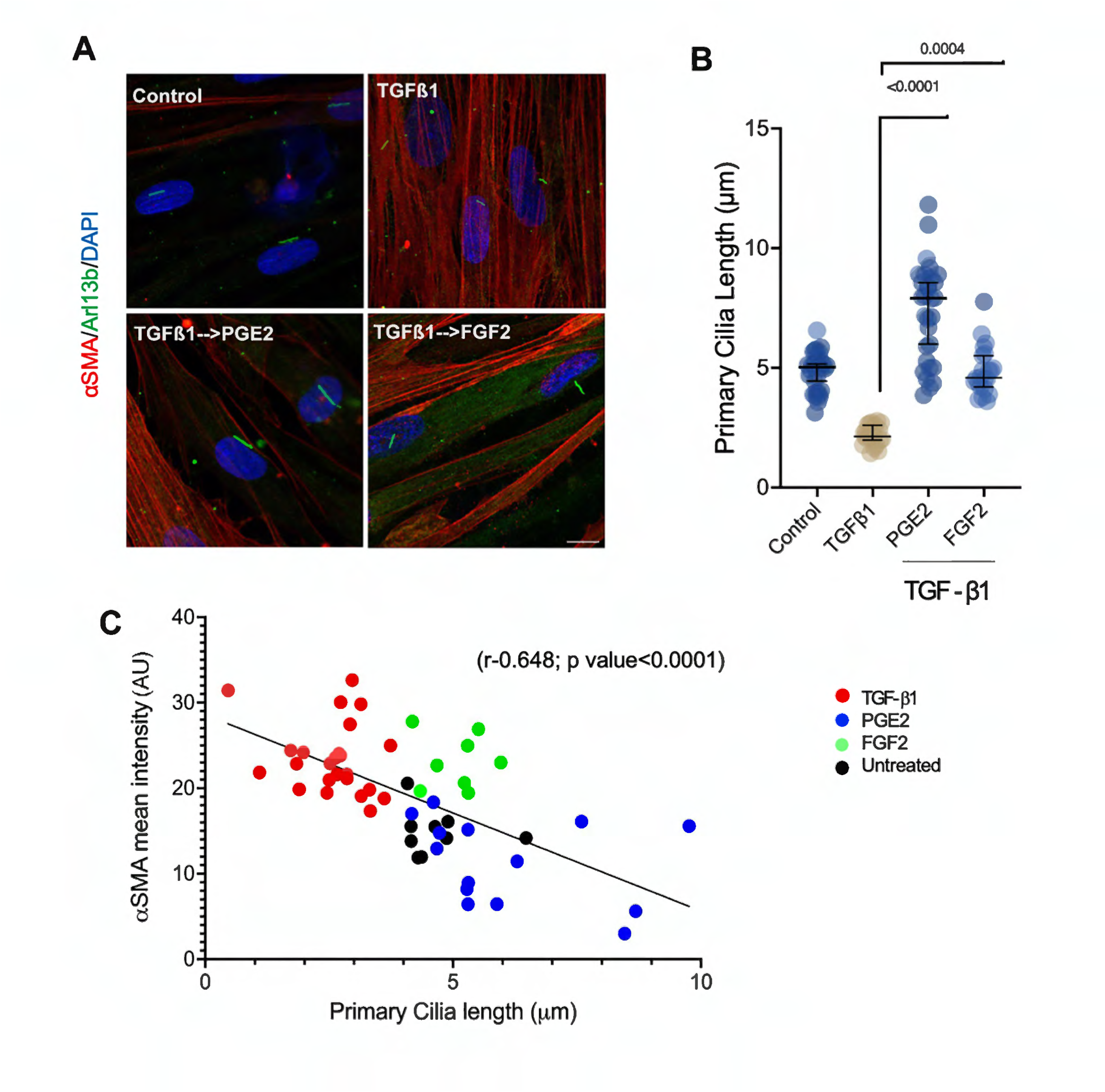
Status of PC length in de-differentiated Lung fibroblasts. **(A)**. Adult lung fibroblast (CCL210) cultures were incubated in media with 0.1% BSA followed by TGF-β1 (10 ng/ml) for 48 h. TGF-β–elicited myofibroblasts were then dedifferentiated in media with PGE2 (1 μM) or FGF2 (50 ng/mL) for 96 h. Fixed fibroblasts were immunolabelled with antibodies to Arl13b and αSMA. Scale bar 10 µm. **(B)**. Quantitative measurements of PC length by 3D reconstruction using ImageJ. Each dot represents an independent cell. (**C**). Correlation of PC length with αSMA levels. Each dot represents PC length and average αSMA intensity from a single cell; Pearson correlation (r-0.648; p value<0.0001). At least 10 determinations per condition were measured.

### Knockdown of *ACTA2* in SSc skin fibroblasts causes PC elongation

The observed negative correlation of PC length with αSMA levels prompted us to investigate how αSMA impacted PC length. For this purpose, SSc fibroblasts transfected with *ACTA2*-specific siRNA were immunolabelled using antibodies to Arl13b or αSMA. Levels of *ACTA2* were downregulated by ∼80% by *ACTA2*-specific siRNA in SSc fibroblasts, whereas no change in *COL1A1* levels was noted (Fig. 4A-C). Notably, loss of the activated myofibroblast phenotype in SSc fibroblasts with *ACTA2* knockdown was associated with elongation of PC (Fig. 4D, E).

**Figure 4:**
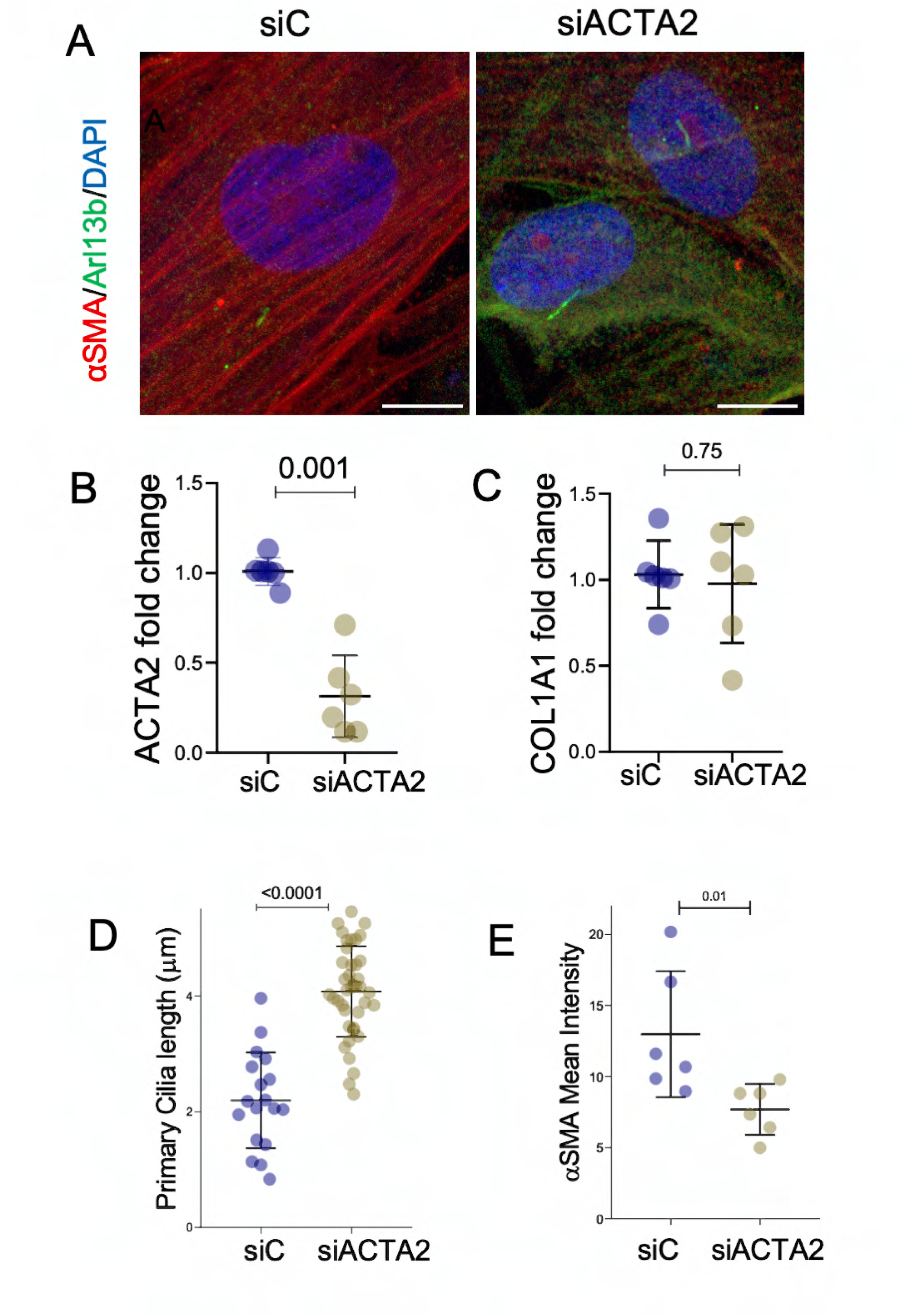
Knockdown of *ACTA2* increases PC length. (**A)**. SSc skin fibroblasts (n=3) were transfected with *ACTA2* or control siRNA. After 24 h incubation, cells were fixed and immunolabelled using antibodies against Arl13b (green) and αSMA (red). Representative images; scale bar 10 μm. (**B,C)**. qPCR (mean ± SD) normalized with GAPDH. Unpaired t test. **(D)**. Quantitative measurements of PC length by 3D reconstruction using ImageJ. Each dot represents an independent cell. **(E)**. αSMA intensity (per individual cell) was measured by ImageJ. siC= siRNA control, siACTA2= ACTA2 siRNA

### Pharmacological PC elongation in SSc skin fibroblasts mitigates their fibrotic phenotype

The concept of therapeutic reprogramming of organelle morphology was first proposed over a decade ago (*37*), but has remained mostly in the theoretical realm (*38, 39*). We hypothesized that restoring PC length in SSc fibroblasts could mitigate their fibrotic phenotype. For this purpose, SSc skin fibroblasts (n=9) were incubated with LiCl, previously shown to increase PC length (*39*). LiCl treatment resulted in a significant time-dependent increase in PC length, even exceeding that seen in healthy fibroblast levels, and was associated with a substantial reduction of αSMA and type I collagen levels (Fig. 5A-G and Suppl. Fig. S3). To assess whether drug-induced PC elongation in these cells was reversible, confluent serum-starved SSc fibroblasts incubated with LiCl for 12 h were incubated in media in the absence of LiCl for an additional 12 h. Washout of LiCl resulted in reducing PC length in SSc fibroblasts to 4-6 µm, comparable to healthy fibroblasts, but not reaching pre-treatment values (Suppl. Fig. S3B). Remarkably, LiCl-treated SSc fibroblasts retained diminished αSMA levels even upon removal of LiCl, comparable to SSc fibroblasts with continued LiCl exposure (Suppl. Fig. S3A, S3C). These results suggest that LiCl-induced PC elongation and reduced αSMA levels in SSc fibroblasts are persistent. Importantly, LiCl treatment of TGF-β1-treated fibroblasts both prevented and reversed upregulation of *ACTA2* and *COL1A1* gene expression (Suppl. Fig. S4). Treatment of SSc skin fibroblasts (n=2) with Nintedanib, a triple kinase inhibitor approved for the treatment of SSc-associated interstitial lung disease (ILD) (*40*) resulted in significant increase in PC length accompanied by reduction of αSMA levels (Suppl. Fig. S8).

**Figure 5.**
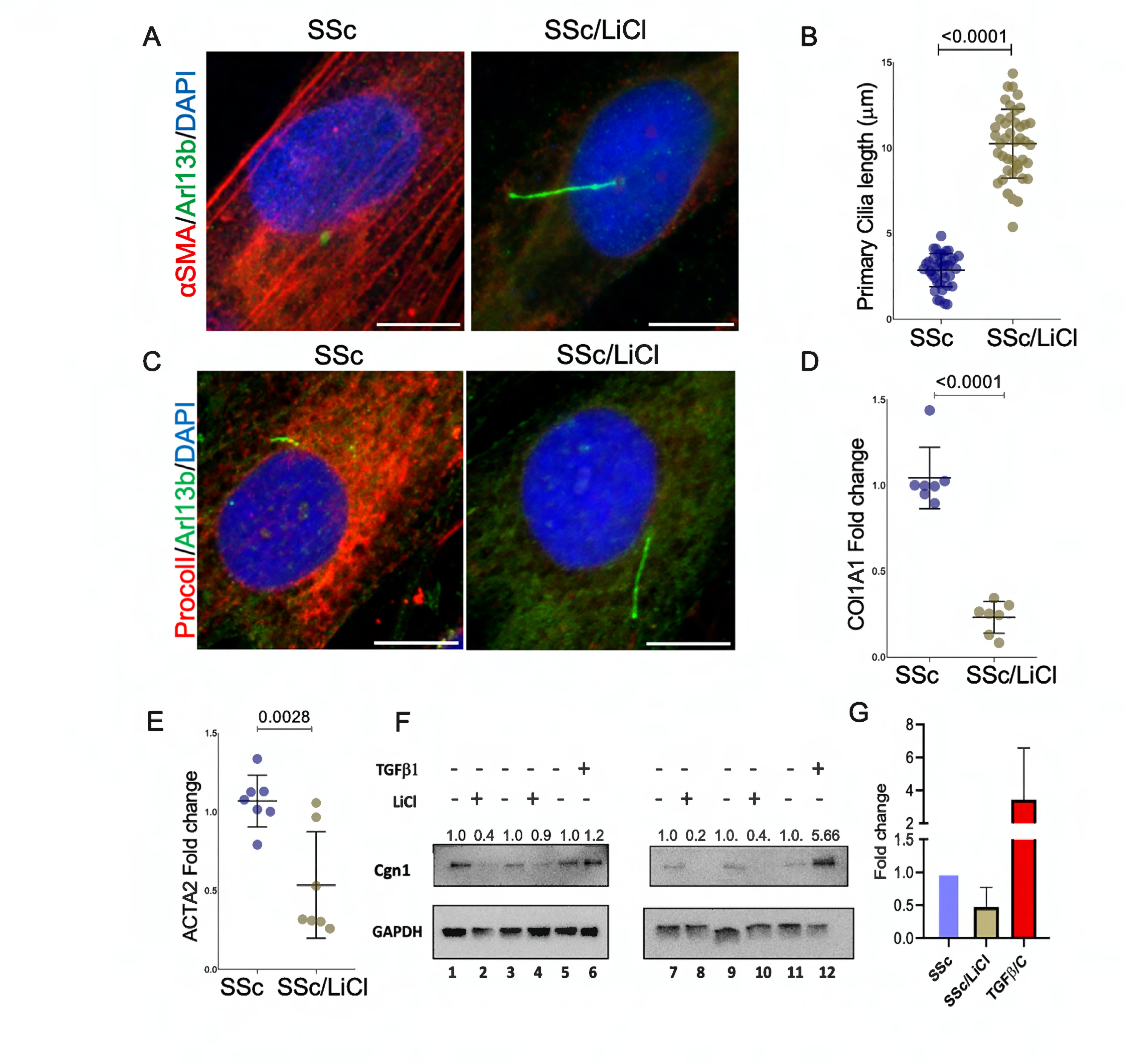
LiCl-mediated PC elongation in SSc fibroblasts attenuates their fibrotic phenotype. **(A, C)** SSc skin fibroblasts (n=4) were incubated in media with 0.1% BSA and without or with LiCl (50 mM) for 24h, followed by immunolabelling using antibodies against Arl13b (green), procollagen type I (red) and αSMA (red). Scale bar 10 μm. **(B)**. Measurements of cilia length by 3D reconstruction using ImageJ. Each dot represents an independent cell. **(D,E)**. Real-time quantitative PCR (mean ± SD) normalized with GAPDH. Unpaired t test **(F).** Cell lysates were subjected to immunoblot analysis. Lanes 1,2: SSc29; 3,4: SSc Dd8;7,8: SSc23; 9,10: SSc39; 5,6,11,12: foreskin fibroblasts. Right panel, **(G)** Fold change (corrected for GAPDH).

### TGF-β1 and LiCl elicit opposing responses in SSc skin fibroblasts

Actin polymerization is implicated in PC shortening via both ciliary resorption and ciliary fission Since myofibroblasts are characterized by actin polymerization, we surmised that this phenomenon underlies PC shortening seen in these cells. We noted that while TGF-β1 reduces PC length and augments αSMA in healthy skin fibroblasts, LiCl treatment has the opposite effect in SSc skin fibroblasts. In order to identify genes and pathways differentially regulated by LiCl, we undertook a transcriptome-wide comparison of LiCl-treated and untreated SSc skin fibroblasts, and untreated and TGF-β1-treated healthy skin fibroblasts (GSE232435). Treatment of SSc fibroblasts with LiCl was associated with differential expression of 4,933 genes (2102 up, 2831 down), and TGF-β1 differentially regulated 4,211 genes (1,792 up, 2,419 down) compared to the corresponding time-matched untreated controls (Suppl. Fig. S5A). Meta-analysis of both RNA-seq datasets revealed 1,718 shared genes (Table S3; Suppl. Fig. S5B) and 13 shared pathways (Table S4; Suppl. Fig. S5C) common to both conditions. We next selected the genes that are regulated in the opposite manner for further analysis. LiCl treatment reduced the expression of profibrotic genes and increased the expression of anti-fibrotic genes; while TGF-β1 treatment had the opposite effects (Fig. 6A). Interestingly, several differentially expressed genes are involved in actin and microtubule polymerizations. These include upregulation of transcripts for actin disassembly (*MICALL1*) and microtubule polymerization (*TPP*, *TPPP3*), and downregulation of transcripts for αSMA, type I collagen, TGF-β1 receptors, and genes implicated in microtubule destabilization (*HDAC6* and *TEAD2*) in LiCl-treated SSc fibroblasts. We validated these results using real time qPCR for selected genes (*TPPP3*, *HDAC6*, *IL6*, *BMP2*, *TEAD2*, and *MICALL1*) (Fig. 6B). Thus, in contrast to TGF-β1, which elicits shortening of PC associated with a fibrotic response, LiCl elongates PC and mitigates fibrotic response in fibrotic fibroblasts.

**Figure 6.**
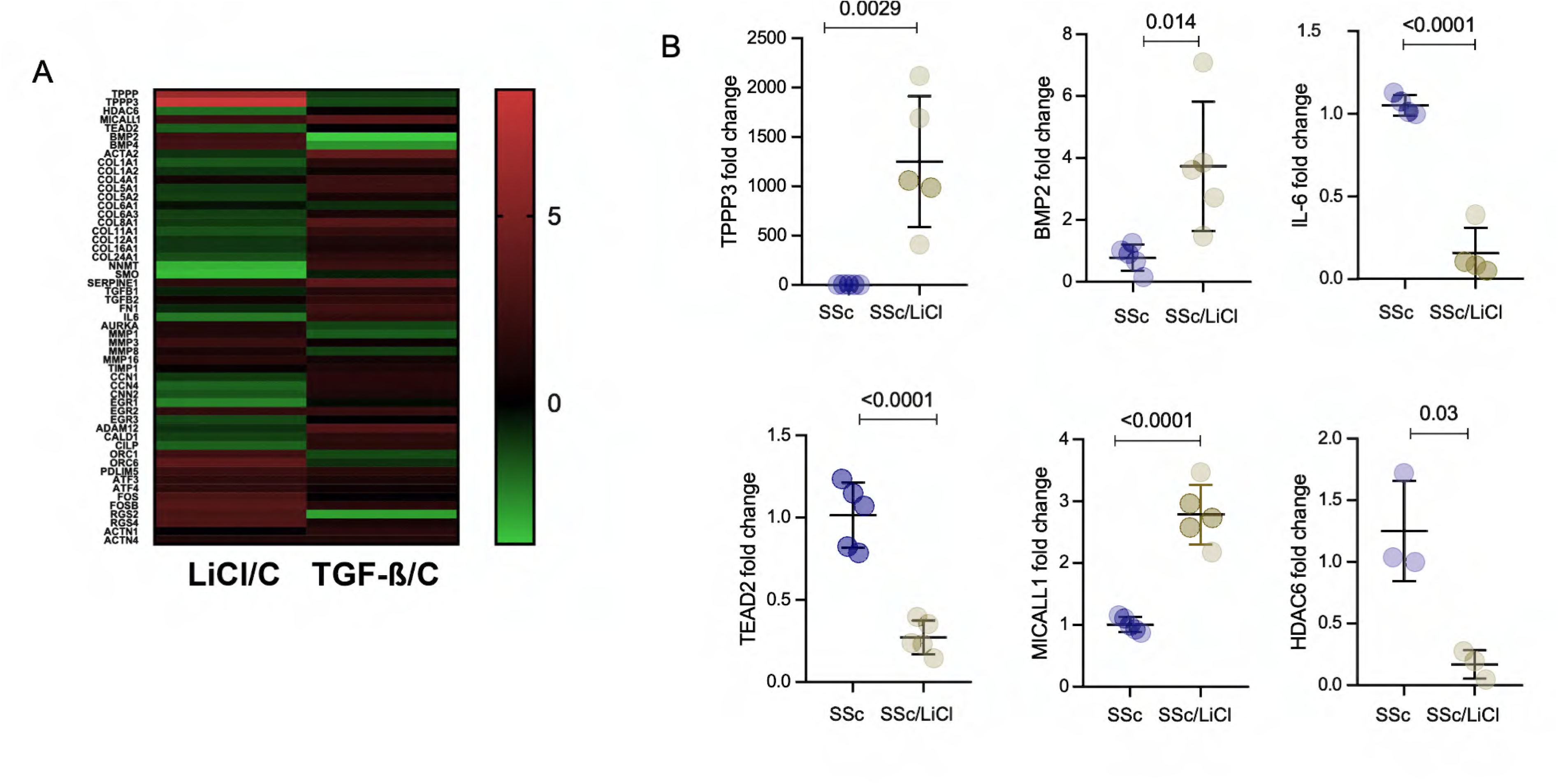
TGF-β1 and LiCl treatment elicit opposite responses in SSc skin fibroblasts. Confluent SSc skin fibroblasts (n=5) were incubated in media with 0.1% BSA without or with LiCl (50 mM) for 24 h, and foreskin fibroblasts (n=3) were incubated with TGF-β1 (10 ng/mL) for 24 h. Total RNA was subjected to bulk RNAseq, using Illumina NovaSeq. **(A)**. Heat map display of fibrosis associated differentially regulated genes by LiCl treated SSc fibroblast (LiCl/C lane) and TGF-β1 treated foreskin fibroblast (TGF-β/C lane) (GSE232435). Color scale depicts range of log_2_fold changes in gene expression. Red, increased; green, decreased. **(B).** Real-time qPCR. Unpaired t test.

### Inhibiting microtubule depolymerization elongates PC in SSc fibroblasts and mitigates their fibrotic phenotype

Our RNA-seq analysis of LiCl-treated SSc skin fibroblasts revealed a significant increase in the expression of microtubule polymerization genes (TPP), and concurrent downregulation of *HDAC6*, which destabilizes ciliary axoneme microtubules, implicating microtubule polymerization in PC elongation as well as fibrosis remission. Single cell RNA sequencing of SSc and healthy skin biopsies revealed that HDAC6 expression was upregulated in SSc fibroblasts and keratinocytes (Suppl. Fig. S7). Analysis of available public database shows an enriched expression of HDAC6 in myofibroblast and smooth muscle cells in IPF lungs (Suppl. Fig. S6A) (42). Furthermore, we also found that HDAC6 expression was upregulated in smooth muscle cells (Suppl. Fig. S6B) in the lungs from bleomycin-treated mice (*43*).

Tubacin, a selective HDAC6 inhibitor, has been shown to cause elongation of PC (*44*). To determine if HDAC6-mediated PC elongation modulates fibrotic responses in SSc skin fibroblasts, we incubated cultures with tubacin for 24 h. We found that tubacin treatment resulted in downregulated *HDAC6* transcript levels and HDAC6 activity in these cells that was accompanied by a significant decrease in αSMA (*ACTA2*) and *COL1A1* expression (Fig. 7A). As expected, we found a significant increase in acetylated α tubulin levels in SSc fibroblast treated with tubacin (Fig. 7C, Suppl. Fig. S10). Acetylated α tubulin is involved in microtubule stabilization, and loss of acetylated α tubulin is reported in aberrant TGF-β1-induced epithelial-to-mesenchymal and wound healing process (*45*). These results indicate that by blocking microtubule destabilization, the HDAC6 inhibitor caused elongation of PC and attenuated the fibrotic cellular phenotype comparable to LiCl treatment (Fig 7 B, D). Of note, the effect of tubacin was dose-dependent, and lower concentrations (1 μM) did not promote PC elongation, suggesting full inhibition of HDAC6 is necessary for this effect.

**Figure 7:**
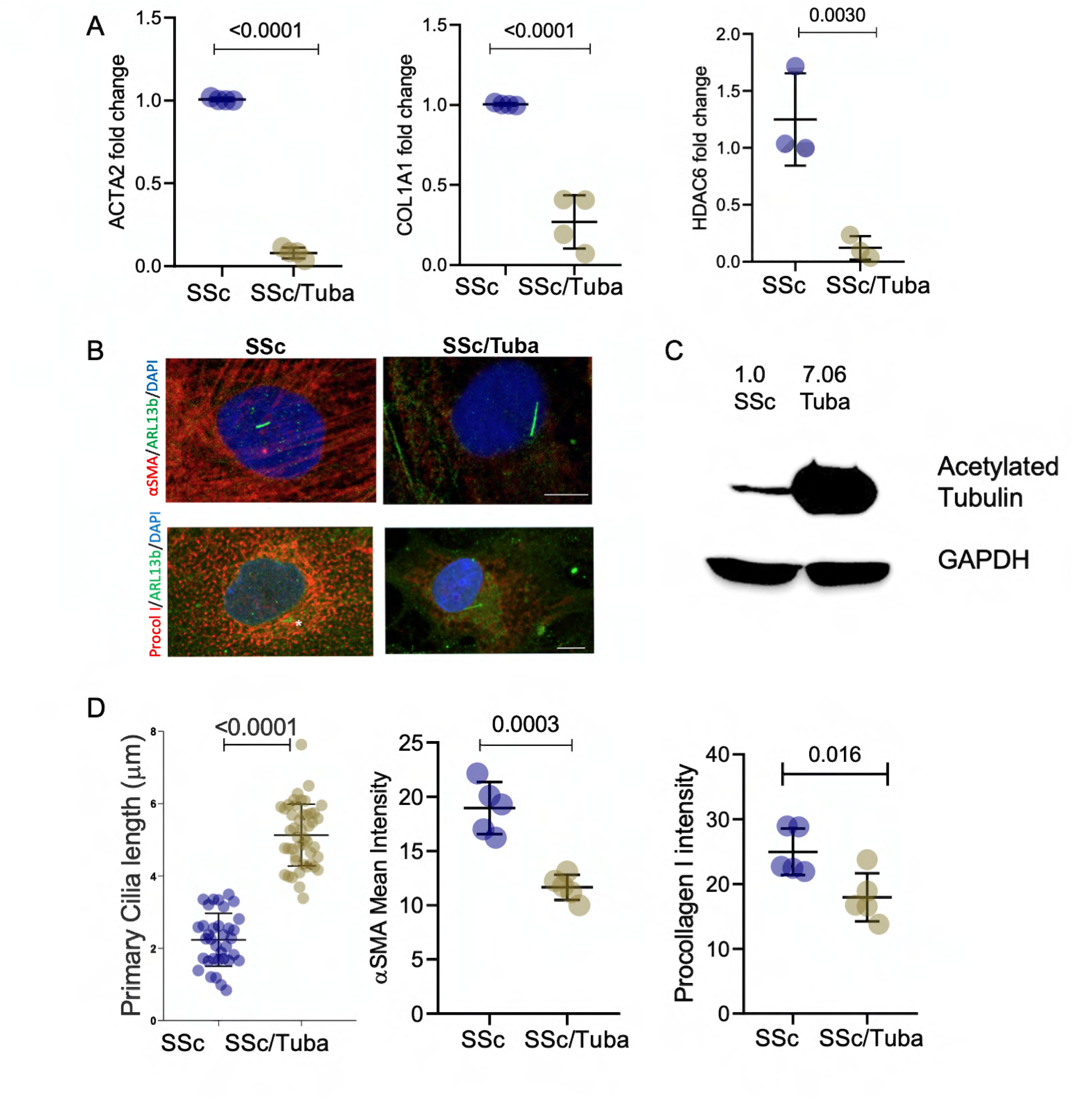
Inhibition of microtubule depolymerization in SSc fibroblasts elongates PC and attenuates fibrotic phenotype. Confluent SSc skin fibroblasts (n=5) were incubated in media with 0.1% BSA without or with Tubacin(Tuba) (20µM) for 24 h. **(A)**. qPCR of normalized with GAPDH. Unpaired t test **(B)**. Immunolabelling with antibodies to Arl13b, αSMA and procollagen I. Representative images; bar=10 μm. **(C)**. Immunoblotting of acetylated α tubulin **(D)**. Quantification of PC length (each dot represents a single cell), from αSMA and procollagen I (each point represents average intensity from 10 cells per cell line) intensity were measured by ImageJ. Unpaired t test.

**Figure 8:**
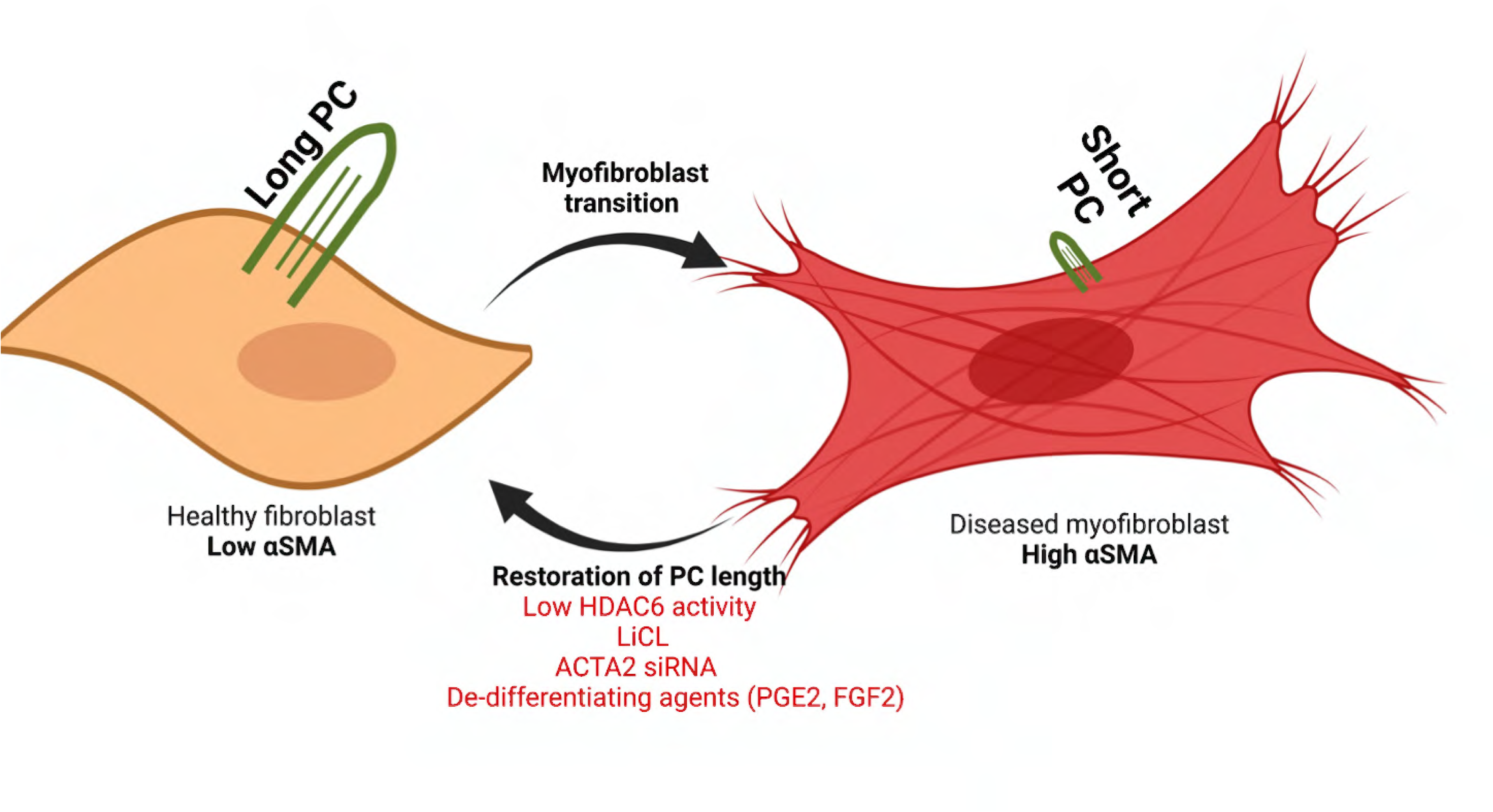
HDAC6 levels correlate with fibrosis and PC length. Schematic model suggesting healthy skin fibroblasts display long PC and almost no αSMA, while SSc fibroblasts display shorter PC and high αSMA. Pharmacological restoration of PC length in SSc fibroblasts can mitigate their fibrotic phenotype. Restoration of PC length by augmented microtubule polymerization (LiCl) or by preventing microtubule depolymerization (HDAC6 inhibition) reduces fibrotic markers and mitigates the fibrotic phenotype.

## DISCUSSION

The identification of cellular mechanisms that promote fibrogenesis, leading to novel therapeutic strategies for SSc, has been our longstanding goal. We recently uncovered compelling evidence connecting the PC-associated gene *SPAG17* with SSc pathophysiology(*19, 20*). Considering the role of PC in tissue remodeling, PC’s importance in states of fibrosis was anticipated (*18, 29*). The extensive actin polymerization, documented to influence PC morphology, is omnipresent in all fibrotic myofibroblasts — this acute observation — rationalized our endeavor to investigate whether morphological alteration of PC is a hallmark of fibrosis. The results described here show that PC shortening in fibroblasts is indeed a common feature shared across multiple fibrotic diseases. The wide range of diversity among various fibrotic diseases presented a challenge for designing broad-spectrum drugs, which can be circumvented by exploiting our discovery. Our work is conceptually innovative in presenting a novel paradigm linking the shortening of PC length in the pathogenesis of fibrosis.

We also observed PC shortening in VEDOSS (Suppl. Fig. S2), an early form of SSc, suggesting that the phenotype is manifested during the early onset of the disease, before fibrosis is clinically evident, thus increasing the chances of that cilia modulation will be effective at limiting fibrosis progression. It is also to be noted that we are measuring PC length in explanted fibroblasts. For cells that also harbor motile cilia, as in lung airway epithelial cells, regulation is complex but similar; motile cilia number per cell increases in IPF (*23*), while PC get shortened and disappears as usual before motile cilia even appeared (*46*). Motile cilia, although structurally similar, are functionally very different than primary cilia and possibly behave differently in fibrosis. The reported increased in ciliary frequency may be due to the increased motile cilia number in the high number of epithelial cells in the fibrotic lesions of IPF lung (*22*).

We used LiCl, nintedanib and the HDAC6 inhibitor tubacin to induce PC elongation and αSMA and type I collagen inhibition. Each of these agents has been shown previously to repress myofibroblast differentiation in cultured fibroblast (*47–49*). Interestingly, in addition to the reported antifibrotic effects of nintedanib in SSc and CCL210 fibroblast, we also observed PC elongation (Suppl. Fig. S8-S9). These results further support our hypothesis — that PC shortening in fibrotic fibroblasts is a hallmark of fibrosis, and restoring PC length to the normal level is associated with recovering to a healthy condition by preventing or mitigating fibrotic phenotypes, irrespective of the mode of recovery. Such a readily measurable visual cellular alteration can be easily exploited for future antifibrotic drug discovery. The potential benefit of selective therapeutic targeting to restore cilia length to treat fibrosis would be enormous.

The negative correlation of αSMA with PC length would incite a myriad of interests in defining the role αSMA in myofibroblast differentiation and morphological regulation of PC. Stress fiber assembly is known to affect PC morphology. Our results document that the inclusion step of αSMA into stress fiber assembly affects PC morphology even further. On the other hand, a lower level of αSMA is associated with a relatively elongated PC. Moreover, an increase in PC length in the SSc cells with knockdown of *ACTA2* transcript suggests that even a reduction of *ACTA2* transcript level can also cause elongation of PC.

Our results suggest that PC elongation via microtubule polymerization lowers the activated myofibroblast phenotype. Microtubule polymerization has been documented before to mitigate fibrotic phenotype(*49*), while microtubule depolymerization is known to induce stress fiber formation(*50*). The presence of polymerized microtubules has been documented to restrict actin assembly(*51*). The precise molecular events linking microtubule and actin cytoskeletal dynamics remain to be further explored in the context of fibrosis. These observations are novel and carry therapeutic implications, suggesting that pharmacological modulation of PC elongation might represent a novel therapeutic strategy for SSc and other currently incurable fibrotic conditions.

## MATERIALS AND METHODS

### Cell culture and reagents

Primary cultures of skin fibroblasts were obtained by explantation from neonatal foreskin samples, or from skin biopsies from SSc patients (n=8) and healthy adults (n=5). Fibroblasts were also explanted from skin biopsies from patients with VEDOSS (n=3) (no detectable skin thickening, modified Rodnan skin score = 0) and classified based on previous publications(*35, 52, 53*). Primary lung fibroblasts from patients with IPF (n=3) were obtained and cultured from explanted lungs from our institutional biorepository as described previously(*54*). Adult CCL210 fibroblasts (ATCC) were used as the control for IPF patients. All CCL210 cell line related experiments were performed in three independent experiments. Orbital fibroblast derived from deep orbital connective tissue waste obtained during surgical decompressions for severe thyroid-associated ophthalmopathy (n=2) (age range 37-58) (TAO) in patients with Grave’s disease(*55*). Healthy orbital fibroblasts (n=2) were obtained from waste tissue obtained during decompression surgery from healthy individuals undergoing cosmetic surgeries.

Biopsies were performed with written informed consent, as per protocols approved by the Institutional Review Board for Human Studies at Northwestern University, the University of Michigan, and the University of Leeds, UK (IRB 00186936 to JV, NRES-011NE to FDG, IRAS 15/NE/0211). Clinical information for subjects in this study is shown in Table S1.

All types of fibroblasts were maintained in Dulbecco’s Modified Eagle’s medium (Gibco, Grand Island, NY) supplemented with 10% fetal bovine serum (Gibco BRL, Grand Island, NY), 1% vitamin solutions, and 2-mM-glutamine (Lonza, Basel, Switzerland). All types of fibroblasts were grown in adherent monolayers in 6 well plastic plates for RNA/cell lysate preparation and 8 well chamber slides for immunofluorescence (IF) studies and studied at low passage.

Fibroblasts were serum-starved for 12 h (1% FBS, Gibco, Grand Island, NY), followed by TGF-β1 (10ng/ml; Peprotech, NJ) and LiCl (50mM; Sigma Aldrich), tubacin (10 and 20µM; Selleckchem, Houston, TX), Nintedanib (24h 2 µM; Selleckchem, Houston, TX) and PGE2 (96hr, 1 μM, Cayman Chemicals, Ann Arbor, MI) or FGF2 (96 h, 50 ng/mL, R&D, Minneapolis, MN) treatments as necessary. LiCl was added to the cultures 24 h after TGF-β1 stimulation for regression studies and one hour pretreatment of LiCl followed by treatment with TGF-β1(10ng/ml) for 24 h for preventive studies.

#### RNA isolation and mRNA quantification

Fibroblasts were subjected to total RNA isolation using the quick-RNA^TM^ MiniPrep Kit from Zymo research. RNA was reverse transcribed to cDNA, followed by qPCR as previously described using Supermix (cDNA Synthesis Supermix; Quanta Biosciences, Gaithersburg, MD) (*56, 57*). Primers used are shown in Table S2.

#### Bulk RNA sequencing and data analysis

Total RNA isolated from LiCl-treated and untreated fibroblast was isolated and subjected to RNA-seq as described(*56, 57*). Data from RNAseq of TGF-β1-treated foreskin fibroblasts was from GSE232435.

#### Immunoblotting analysis

Whole cell lysate was prepared from harvested fibroblasts, and immunoblotting was performed as previously described(*56, 57*). Briefly, Whole cell lysates were fractioned on SDS-PAGE electrophoresis, followed by transfer of proteins. Membranes were blocked and incubated with primary antibodies specific for type I collagen (Southern Biotechnology, Birmingham, AL, 1:1000) and GAPDH (Santacruz, Dallas, Texas, 1:2000), followed by the addition of appropriate secondary antibodies. Membranes were then detected for bands using the enhanced chemiluminescence detection reagent as described(*56, 57*). Band intensities were quantitated using Image J software and normalized with GAPDH loading control.

#### siRNA knockdown

siRNA transfection was performed as described previously(*58*). Briefly, human ACTA2 (sc-43590, Santacruz, Dallas, Texas) and control siRNA (sc-37007, Santacruz, Dallas, Texas) were transfected into cells with Lipofectamine^™^ 3000 (Invitrogen, CA, USA) as per manufacturer protocol. Transfection efficiency was measured via qPCR and immunofluorescence imaging of αSMA staining (Sigma Aldrich, #A52281, Rockville, MD1:400).

#### Immunofluorescence confocal microscopy

Treated and untreated fibroblasts were seeded in 8 well chamber slides, and Immunofluorescence staining was performed as previously described(*56, 57*). Briefly, fixed fibroblasts were incubated with primary antibodies specific to procollagen I (Invitrogen, Waltham, Massachusetts, 1:200), αSMA (sigma, Rockville, MD1:400), Acetyl-α-Tubulin (Cell Signaling #5335, Danvers, MA 1:500) and ARL13b (Proteintech, 17711-1-AP, Rosemont, IL) followed by addition of appropriate secondary antibodies and DAPI (Sigma-Aldrich, Rockville, MD0.2 μg/mL). Images were captured using the Leica SP8 confocal system (The Microscopy Core facility, U of M) or the Zeiss LSM880 confocal system (Bioimaging Facility, University of Leeds). Cilia length was measured using 3D reconstruction using ImageJ 3D viewer. Collagen and αSMA immunofluorescence intensities were measured using ImageJ.

#### Single cell RNAseq data analysis

Biopsies from patients with early diffuse SSc were subjected to scRNAseq as described (*59*). GSE141259 and GSE135893 datasets were used in this study for HDAC6 transcript level.

#### Statistical Analysis

Student’s t-test (Unpaired) was used to compare two groups, with a p-value <0.05 considered statistically significant. Data are presented as means ± SEM, and Graph Pad Prism (Graph Pad Software version 8, Graph Pad Software Inc., CA) was used for data analysis.

## Supporting information

Supplementary informations

## Acknowledgments

We thank members of the University of Michigan Leica Imaging Core facilities. We also thank Carolina Jorro for the quantification of confocal images and Prof. Terry Smith and Roshini Fernando for providing TAO and healthy orbital fibroblast from the University of Michigan. We acknowledge support from the Bioinformatics Core of the University of Michigan Medical School’s Biomedical Research Core Facilities (RRID: SCR_019168).

## Author contributions

DB, MET, and JV conceptualized the study. PV, BY, RW, and DB performed all the experiments and data analysis, while DB, JV, and FDG oversaw all the experiments. SB shared experimental expertise and helped in drafting the manuscript. SF performed lung fibroblast experiments. DK and SF provided skin and lung fibroblasts. JG provided the single-cell RNA sequence analysis for HDAC6. RR and RW performed experiments with VEDOSS dermal fibroblasts. DB prepared the original draft of the manuscript. DB, PV, SB, MET, FDG, JG, and JV edited the manuscript and provided input. All the authors reviewed the final manuscript.

## Data and materials availability

Raw bulk RNASeq data are stored in the University of Michigan Data Den archival facility. All the other raw and processed data is stored in the University of Michigan Dropbox and shared folder S:\Intmed_Rheum\Research\Varga_Lab. All data are available upon request.

## REFERENCES

1. T. A. Wynn, Common and unique mechanisms regulate fibrosis in various fibroproliferative diseases. J Clin Invest 117, 524–529 (2007).

2. C. P. Denton, D. Khanna, Systemic sclerosis. Lancet 390, 1685–1699 (2017).

3. J. Varga, D. Abraham, Systemic sclerosis: a prototypic multisystem fibrotic disorder. J Clin Invest 117, 557–567 (2007).

4. A. M. Hoffmann-Vold et al., Progressive interstitial lung disease in patients with systemic sclerosis-associated interstitial lung disease in the EUSTAR database. Ann Rheum Dis 80, 219–227 (2021).

5. Y. Allanore et al., Systemic sclerosis. Nat Rev Dis Primers 1, 15002 (2015).

6. E. R. Volkmann, J. Varga, Emerging targets of disease-modifying therapy for systemic sclerosis. Nat Rev Rheumatol 15, 208–224 (2019).

7. D. J. Lederer, F. J. Martinez, Idiopathic Pulmonary Fibrosis. N Engl J Med 378, 1811–1823 (2018).

8. P. N. Taylor et al., New insights into the pathogenesis and nonsurgical management of Graves orbitopathy. Nat Rev Endocrinol 16, 104–116 (2020).

9. S. F. Madsen et al., Fibroblasts are not just fibroblasts: clear differences between dermal and pulmonary fibroblasts’ response to fibrotic growth factors. Sci Rep 13, 9411 (2023).

10. B. Hinz et al., Recent developments in myofibroblast biology: paradigms for connective tissue remodeling. The American journal of pathology 180, 1340–1355 (2012).

11. J. H. W. Distler et al., Shared and distinct mechanisms of fibrosis. Nat Rev Rheumatol 15, 705–730 (2019).

12. B. Hinz, G. Gabbiani, C. Chaponnier, The NH2-terminal peptide of alpha-smooth muscle actin inhibits force generation by the myofibroblast in vitro and in vivo. J Cell Biol 157, 657–663 (2002).

13. P. Avasthi, W. F. Marshall, Stages of ciliogenesis and regulation of ciliary length. Differentiation 83, S30–42 (2012).

14. J. J. Malicki, C. A. Johnson, The Cilium: Cellular Antenna and Central Processing Unit. Trends in cell biology 27, 126–140 (2017).

15. R. Pala, N. Alomari, S. M. Nauli, Primary Cilium-Dependent Signaling Mechanisms. Int J Mol Sci 18, (2017).

16. Z. Anvarian, K. Mykytyn, S. Mukhopadhyay, L. B. Pedersen, S. T. Christensen, Cellular signalling by primary cilia in development, organ function and disease. Nat Rev Nephrol 15, 199–219 (2019).

17. M. Hosio et al., Primary Ciliary Signaling in the Skin-Contribution to Wound Healing and Scarring. Front Cell Dev Biol 8, 578384 (2020).

18. D. Bhattacharyya, M. E. Teves, J. Varga, The dynamic organelle primary cilia: emerging roles in organ fibrosis. Curr Opin Rheumatol 33, 495–504 (2021).

19. E. D. O. Roberson et al., Alterations of the Primary Cilia Gene SPAG17 and SOX9 Locus Noncoding RNAs Identified by RNA-Sequencing Analysis in Patients With Systemic Sclerosis. Arthritis Rheumatol 75, 108–119 (2023).

20. P. Sapao et al., Reduced SPAG17 Expression in Systemic Sclerosis Triggers Myofibroblast Transition and Drives Fibrosis. J Invest Dermatol 143, 284–293 (2023).

21. E. Villalobos et al., Fibroblast Primary Cilia Are Required for Cardiac Fibrosis. Circulation 139, 2342–2357 (2019).

22. J. Lee et al., Increased Primary Cilia in Idiopathic Pulmonary Fibrosis. Mol Cells 41, 224–233 (2018).

23. E. Kim et al., Aberrant Multiciliogenesis in Idiopathic Pulmonary Fibrosis. Am J Respir Cell Mol Biol 67, 188–200 (2022).

24. A. Andreu-Cervera, M. Catala, S. Schneider-Maunoury, Ciliopathies and hedgehog-related forebrain developmental disorders. Neurobiol Dis, 105236 (2020).

25. Y. Li, K. Ling, J. Hu, The emerging role of Arf/Arl small GTPases in cilia and ciliopathies. Journal of cellular biochemistry 113, 2201–2207 (2012).

26. D. J. McConnachie, J. L. Stow, A. J. Mallett, Ciliopathies and the Kidney: A Review. Am J Kidney Dis 77, 410–419 (2021).

27. J. F. Reiter, M. R. Leroux, Genes and molecular pathways underpinning ciliopathies. Nat Rev Mol Cell Biol 18, 533–547 (2017).

28. A. D. Egorova et al., Lack of primary cilia primes shear-induced endothelial-to-mesenchymal transition. Circ Res 108, 1093–1101 (2011).

29. M. E. Teves, J. F. Strauss, 3rd, P. Sapao, B. Shi, J. Varga, The Primary Cilium: Emerging Role as a Key Player in Fibrosis. Curr Rheumatol Rep 21, 29 (2019).

30. M. Rozycki et al., The fate of the primary cilium during myofibroblast transition. Mol Biol Cell 25, 643–657 (2014).

31. M. Bershteyn, S. X. Atwood, W. M. Woo, M. Li, A. E. Oro, MIM and cortactin antagonism regulates ciliogenesis and hedgehog signaling. Dev Cell 19, 270–283 (2010).

32. J. Kim et al., Functional genomic screen for modulators of ciliogenesis and cilium length. Nature 464, 1048–1051 (2010).

33. N. Sharma, Z. A. Kosan, J. E. Stallworth, N. F. Berbari, B. K. Yoder, Soluble levels of cytosolic tubulin regulate ciliary length control. Mol Biol Cell 22, 806–816 (2011).

34. E. D. Gigante, M. R. Taylor, A. A. Ivanova, R. A. Kahn, T. Caspary, ARL13B regulates Sonic hedgehog signaling from outside primary cilia. Elife 9, (2020).

35. L. S. Lonzetti et al., Updating the American College of Rheumatology preliminary classification criteria for systemic sclerosis: addition of severe nailfold capillaroscopy abnormalities markedly increases the sensitivity for limited scleroderma. Arthritis Rheum 44, 735–736 (2001).

36. S. M. Fortier, et al., Myofibroblast dedifferentiation proceeds via distinct transcriptomic and phenotypic transitions. JCI Insight 6, (2021).

37. W. F. Marshall, Organelle size control systems: from cell geometry to organelle-directed medicine. Bioessays 34, 721–724 (2012).

38. S. A. Gradilone, M. J. L. Pisarello, N. F. LaRusso, Primary Cilia in Tumor Biology: The Primary Cilium as a Therapeutic Target in Cholangiocarcinoma. Curr Drug Targets 18, 958–963 (2017).

39. C. L. Thompson, A. Wiles, C. A. Poole, M. M. Knight, Lithium chloride modulates chondrocyte primary cilia and inhibits Hedgehog signaling. FASEB J 30, 716–726 (2016).

40. D. Khanna et al., Systemic Sclerosis-Associated Interstitial Lung Disease: How to Incorporate Two Food and Drug Administration-Approved Therapies in Clinical Practice. Arthritis Rheumatol 74, 13–27 (2022).

41. S. Stilling, T. Kalliakoudas, H. Benninghoven-Frey, T. Inoue, B. H. Falkenburger, PIP2 determines length and stability of primary cilia by balancing membrane turnovers. Commun Biol 5, 93 (2022).

42. A. C. Habermann et al., Single-cell RNA sequencing reveals profibrotic roles of distinct epithelial and mesenchymal lineages in pulmonary fibrosis. Sci Adv 6, eaba1972 (2020).

43. M. Strunz et al., Alveolar regeneration through a Krt8+ transitional stem cell state that persists in human lung fibrosis. Nat Commun 11, 3559 (2020).

44. S. Fu et al., Mechanical loading inhibits cartilage inflammatory signalling via an HDAC6 and IFT-dependent mechanism regulating primary cilia elongation. Osteoarthritis Cartilage 27, 1064–1074 (2019).

45. S. Gu et al., Loss of alpha-Tubulin Acetylation Is Associated with TGF-beta-induced Epithelial-Mesenchymal Transition. J Biol Chem 291, 5396–5405 (2016).

46. R. Jain et al., Temporal relationship between primary and motile ciliogenesis in airway epithelial cells. Am J Respir Cell Mol Biol 43, 731–739 (2010).

47. E. J. Chung, Y. H. Sohn, S. H. Kwon, S. A. Jung, J. H. Lee, Lithium chloride inhibits TGF-beta1-induced myofibroblast transdifferentiation via PI3K/Akt pathway in cultured fibroblasts from Tenon’s capsule of the human eye. Biotechnol Lett 36, 1217–1224 (2014).

48. P. Pinol-Jurado et al., Nintedanib decreases muscle fibrosis and improves muscle function in a murine model of dystrophinopathy. Cell Death Dis 9, 776 (2018).

49. S. Saito et al., Tubastatin ameliorates pulmonary fibrosis by targeting the TGFbeta-PI3K-Akt pathway. PLoS One 12, e0186615 (2017).

50. B. P. Liu, M. Chrzanowska-Wodnicka, K. Burridge, Microtubule depolymerization induces stress fibers, focal adhesions, and DNA synthesis via the GTP-binding protein Rho. Cell Adhes Commun 5, 249–255 (1998).

51. J. Pineau et al., Microtubules restrict F-actin polymerization to the immune synapse via GEF-H1 to maintain polarity in lymphocytes. Elife 11, (2022).

52. J. Avouac et al., Preliminary criteria for the very early diagnosis of systemic sclerosis: results of a Delphi Consensus Study from EULAR Scleroderma Trials and Research Group. Ann Rheum Dis 70, 476–481 (2011).

53. M. Matucci-Cerinic et al., The challenge of early systemic sclerosis for the EULAR Scleroderma Trial and Research group (EUSTAR) community. It is time to cut the Gordian knot and develop a prevention or rescue strategy. Ann Rheum Dis 68, 1377–1380 (2009).

54. A. M. Scruggs, G. Grabauskas, S. K. Huang, The Role of KCNMB1 and BK Channels in Myofibroblast Differentiation and Pulmonary Fibrosis. Am J Respir Cell Mol Biol 62, 191–203 (2020).

55. T. J. Smith et al., Fibroblasts expressing the thyrotropin receptor overarch thyroid and orbit in Graves’ disease. J Clin Endocrinol Metab 96, 3827–3837 (2011).

56. S. Bale, et al., Pharmacological inhibition of TAK1 prevents and induces regression of experimental organ fibrosis. JCI Insight 8, (2023).

57. W. Wang et al., Fibroblast A20 governs fibrosis susceptibility and its repression by DREAM promotes fibrosis in multiple organs. Nat Commun 13, 6358 (2022).

58. S. Bhattacharyya, W. Wang, L. V. Graham, J. Varga, A20 suppresses canonical Smad-dependent fibroblast activation: novel function for an endogenous inflammatory modulator. Arthritis Res Ther 18, 216 (2016).

59. B. Shi et al., Senescent Cells Accumulate in Systemic Sclerosis Skin. J Invest Dermatol 143, 661–664 e665 (2023).

