## Supplementary informations for "Morphological Reprogramming of Primary Cilia Length Mitigates the Fibrotic Phenotype in Fibroblasts Across Diverse Fibrotic Conditions"

Priyanka Verma et al

,

This PDF file includes:

Supplementary Text

Tables S1 to S4

Figs. S1 to S10

Table S1- Clinical characteristics of the study subjects

| Study code | Age | Sex | Type | Ethnicity | Race | MRSS score |
|---|---|---|---|---|---|---|
| SSc_23 | 37 | M | Early Diffuse SSc | Non-Hispanic | Caucasian | 47 |
| SSc_24 | 69 | M | Late Diffuse SSc | Non-Hispanic | Caucasian | 14 |
| SSc_25 | 45 | F | Early Limitted SSc | Non-Hispanic | Caucasian | 12 |
| SSc_29 | 50 | F | Early Diffuse SSc | Non-Hispanic | Caucasian | 33 |
| SSc_39 | 54 | F | VEDOSS (Early SSc) | Non-Hispanic | Caucasian | 0 |
| DoDET-003 | 51 | F | Early diffuse SSc | N/A | Caucasian | N/A |
| DoDET-005 | 43 | F | Early diffuse SSc | N/A | Caucasian | N/A |
| DoDET-007 | 54 | F | Early diffuse SSc | N/A | Non-Caucasian | N/A |
| DoDET-008 | 61 | F | Early diffuse SSc | N/A | Caucasian | N/A |
| DoDET-009 | 67 | M | Early diffuse SSc | N/A | Caucasian | N/A |
| IPF-96 | 64 | M | IPF | N/A | N/A | N/A |
| IPF-106 | 66 | M | IPF | N/A | N/A | N/A |
| IPF-146 | 28 | M | Unclassified IPF | N/A | N/A | N/A |
| SSc1 | 43 | M | Diffuse cutaneous systemic sclerosis | N/A | White - British | 43 |
| SSc2 | 67 | F | Diffuse cutaneous systemic sclerosis | N/A | White - British | 35 |
| SSc3 | 35 | F | Diffuse cutaneous systemic sclerosis | N/A | Asian - Bangladeshi | 1 |
| VEDOSS 1 | 34 | F | VEDOSS | N/A | White - British | 0 |

| | | | | | | |
|---|---|---|---|---|---|---|
| VEDOSS 2 | 39 | M | VEDOSS | N/A | Asian - Indian | 0 |
| VEDOSS 3 | 66 | F | VEDOSS | N/A | White - British | 0 |
| WADA | N/A | N/A | TAO | N/A | N/A | |
| SLMUG | N/A | N/A | TAO | N/A | N/A | |

F, Female; M, Male; N/A, not applicable. Early, disease duration <3 years from first non-Raynaud disease manifestation; late >3 years from first non-Raynaud manifestation. MRSS, modified Rodnan skin score (1 to 51). SSc, systemic sclerosis; IPF, Idiopathic Pulmonary Fibrosis; VEDOSS, Very Early Diagnosis of systemic Sclerosis, TAO, Thyroid-associated ophthalmopathy. Controls were healthy subjects (90% female; median age, 56 years; range, 47 to 68 years).

TableS2 -List of primers used for gene expression.

| | |
|---|---|
| hTPPP3-F | Forward: 5'- AAGTCTGCTCGGGTCATCAAC -3' |
| hTPPP3-R | Reverse: 5'- GAGCCCGTGTATCTGCTGG -3' |
| hBMP2-F | Forward: 5'- ACCCGCTGTCTTCTAGCGT-3' |
| hBMP2-F | Reverse: 5'- TTTCAGGCCGAACATGCTGAG -3' |
| hIL-6-F | Forward: 5'- AAATTCGGTACATCCTCGACGG -3' |
| hIL-6-R | Reverse: 5'- GGAAGGTTCAGGTTGTTTTCTGC -3' |
| hACTA-F | Forward: 5'- CAGGGCTGTTTTCCCATCCAT -3' |
| hACTA-R | Reverse: 5'- GCCATGTTCTATCGGGTACTTC -3' |
| hCOL1A1-F | Forward: 5'- CTGAGTCAGCAGATTGAGAACA -3' |
| hCOL1A1-R | Reverse: 5'- AGGTTGCAGCCTTGGTTAG -3' |
| hTEAD2-F | Forward: 5'- CTTCGTGGAACCGCCAGAT -3' |
| hTEAD2-R | Reverse: 5'- GGAGGCCACCCTTTTTCTCA -3' |
| hGAPDH-F | Forward: 5'- CATGAGAAGTATGACAACAGCCT -3' |
| hGAPDH-R | Reverse: 5'- AGTCCTTCCACGATACCAAAGT -3' |
| hMICALL1-F | Forward: 5'- CTCATCCACGAGAAGCACCTAC -3' |
| hMICALL1-R | Reverse: 5'- GCTCATACTCGACATCAGCCTG -3' |
| hHDAC6-F | Forward: 5'- GAGGGAGAACTCCGTGTCCTA -3' |
| hHDAC6-R | Reverse: 5'- AATAGCCATCCATAAGACTGTGC -3' |

Table S3 Common genes found in the meta-analysis of both datasets.

Common genes with >1-fold increase or decrease and adjpv < 0.01 were defined as being differentially expressed. LiCl lane: Differentially regulated genes by LiCl treated SSc fibroblast; TGF-β1 lane: TGF-β1 treated foreskin fibroblast (GSE232435); adjpv= adjusted p value

| entity | name | $Log_2$fc (LiCL) | adjpv (LiCl) | $Log_2$fc (TGF-β1) | adjpv (TGF-β1) |
|---|---|---|---|---|---|
| 22848 | AAK1 | 1.170906 | 0.000001 | 0.748867 | 0.000001 |
| 16 | AARS1 | 1.584249 | 0.000001 | 0.748588 | 0.000001 |
| 10349 | ABCA10 | -1.86513 | 2.21E-05 | -1.2164 | 0.033382 |
| 23461 | ABCA5 | -2.08775 | 0.000001 | -1.15708 | 0.000001 |
| 23460 | ABCA6 | -2.62958 | 0.000001 | -1.99291 | 0.000001 |
| 10347 | ABCA7 | -1.60976 | 0.000001 | -1.37202 | 4.45E-05 |
| 10351 | ABCA8 | -2.05702 | 5.64E-05 | -0.91585 | 0.0066 |

| | | | | | |
|---|---|---|---|---|---|
| 10350 | ABCA9 | -1.78077 | 0.000001 | -1.50566 | 0.000001 |
| 5244 | ABCB4 | -1.14021 | 0.005894 | -1.25502 | 3.75E-06 |
| 23457 | ABCB9 | -1.10856 | 0.000883 | -0.65852 | 0.03999 |
| 8714 | ABCC3 | -1.23982 | 0.000001 | 1.933112 | 0.000001 |
| 1.01E+08 | ABHD14A-ACY1 | -1.73136 | 2.18E-05 | -0.81381 | 0.032926 |
| 116236 | ABHD15 | -1.21153 | 0.000001 | -0.84602 | 0.000001 |
| 51099 | ABHD5 | 1.478639 | 0.000001 | 0.953823 | 0.000001 |
| 79575 | ABHD8 | -1.19451 | 1.52E-05 | -0.62963 | 0.000001 |
| 3983 | ABLIM1 | 1.667841 | 0.000001 | 1.896931 | 0.000001 |
| 84142 | ABRAXAS1 | -1.87606 | 0.000001 | -0.69759 | 4.79E-06 |
| 84129 | ACAD11 | -1.4137 | 0.000001 | -0.81371 | 1.56E-06 |
| 35 | ACADS | -1.45334 | 0.000001 | -0.87566 | 0.000001 |
| 36 | ACADSB | -1.67167 | 0.000001 | -0.82813 | 0.000001 |
| 79777 | ACBD4 | -2.65035 | 0.000001 | -0.9891 | 1.82E-06 |
| 57007 | ACKR3 | -1.57459 | 0.000001 | 1.632617 | 0.000001 |
| 51554 | ACKR4 | -1.64078 | 0.000001 | -3.88959 | 0.000001 |
| 285737 | ACKR4P1 | -1.96625 | 0.005044 | -1.41043 | 0.02032 |
| 641371 | ACOT1 | -2.21189 | 0.001579 | 0.604286 | 2.35E-05 |
| 8309 | ACOX2 | -3.06353 | 0.000001 | -1.41072 | 0.000001 |
| 55289 | ACOXL | -3.25479 | 0.000001 | -0.70819 | 0.003461 |
| 80221 | ACSF2 | -2.47179 | 0.000001 | -0.83945 | 0.000001 |
| 23305 | ACSL6 | -2.04082 | 0.000124 | -1.91789 | 0.000994 |
| 84532 | ACSS1 | -2.50223 | 0.000001 | -2.71298 | 0.000001 |
| 55902 | ACSS2 | -2.97012 | 0.000001 | -1.3871 | 0.000001 |
| 81 | ACTN4 | 1.24483 | 0.000001 | 1.224808 | 0.000001 |
| 94 | ACVRL1 | -1.27252 | 0.000001 | -1.93177 | 0.000001 |
| 51816 | ADA2 | -1.3502 | 2.03E-06 | -1.00244 | 0.00412 |
| 53616 | ADAM22 | -1.02588 | 0.002062 | -2.45803 | 0.000001 |
| 10863 | ADAM28 | -2.37122 | 0.004864 | -1.42741 | 0.014999 |

| | | | | | |
|---|---|---|---|---|---|
| 80332 | ADAM33 | -2.33169 | 0.000001 | -1.45774 | 0.000001 |
| 81794 | ADAMTS10 | -2.73718 | 0.000001 | 1.271882 | 0.000001 |
| 170689 | ADAMTS15 | -2.25476 | 9.91E-05 | -1.85378 | 0.000001 |
| 642935 | ADAMTS7P4 | -2.72082 | 0.000001 | -4.29715 | 0.000001 |
| 11095 | ADAMTS8 | -2.18554 | 3.49E-05 | -1.5384 | 2.26E-06 |
| 92949 | ADAMTSL1 | -1.89998 | 0.000001 | -0.9731 | 1.2E-05 |
| 90956 | ADCK2 | -1.37315 | 0.000001 | -0.70716 | 3.46E-06 |
| 107 | ADCY1 | 2.872086 | 0.000001 | -2.04849 | 0.000001 |
| 114 | ADCY8 | 3.653907 | 0.000001 | -1.77738 | 0.000001 |
| 283383 | ADGRD1 | -2.61278 | 0.000001 | -1.60578 | 1.05E-05 |
| 976 | ADGRE5 | 2.641813 | 0.000001 | -1.01758 | 0.000001 |
| 57211 | ADGRG6 | 1.170319 | 0.000491 | 0.983619 | 6.52E-05 |
| 125 | ADH1B | -4.14724 | 0.000001 | -4.95257 | 0.000001 |
| 137872 | ADHFE1 | -1.88855 | 2.84E-05 | -1.53011 | 0.000964 |
| 133 | ADM | -3.31834 | 0.000001 | -1.50959 | 0.000001 |
| 147 | ADRA1B | 1.49586 | 7.5E-06 | -2.00845 | 1.33E-06 |
| 150 | ADRA2A | -2.94798 | 0.000001 | 0.628874 | 0.023555 |
| 84830 | ADTRP | 4.018464 | 0.000001 | 1.665138 | 0.000001 |
| 60312 | AFAP1 | -1.1526 | 0.000001 | 0.61826 | 1.58E-06 |
| 414189 | AGAP6 | 1.339873 | 0.000001 | 0.909709 | 5.94E-06 |
| 60509 | AGBL5 | -1.91086 | 0.000001 | -0.61718 | 3.36E-06 |
| 56895 | AGPAT4 | 1.208082 | 0.000001 | 0.941086 | 0.000001 |
| 185 | AGTR1 | -1.10417 | 0.001436 | 1.327693 | 0.000001 |
| 57085 | AGTRAP | -1.74816 | 0.000001 | 1.418675 | 0.000001 |
| 113146 | AHNAK2 | -1.1328 | 0.000001 | -1.38052 | 0.000001 |
| 84962 | AJUBA | -1.806 | 0.000001 | 0.703911 | 0.000001 |
| 122481 | AK7 | -2.52646 | 0.000001 | -1.42043 | 1.9E-05 |
| 1.19E+08 | AKAP1-DT | -1.65722 | 0.000989 | -1.12321 | 0.015844 |
| 254268 | AKNAD1 | 1.449797 | 0.000739 | -2.87197 | 0.000001 |

| | | | | | |
|---|---|---|---|---|---|
| 1645 | AKR1C1 | 1.671409 | 0.002695 | -0.99542 | 0.025879 |
| 210 | ALAD | -1.2871 | 0.000001 | -0.76022 | 0.000001 |
| 160428 | ALDH1L2 | -2.49106 | 0.000001 | 1.854233 | 0.000001 |
| 221 | ALDH3B1 | -1.82119 | 0.000001 | -1.85038 | 0.000001 |
| 7915 | ALDH5A1 | -2.6981 | 1.09E-05 | -1.00793 | 0.000953 |
| 4329 | ALDH6A1 | -2.64081 | 0.000001 | -1.05247 | 0.000001 |
| 501 | ALDH7A1 | -2.69888 | 0.000001 | -0.6905 | 0.000001 |
| 230 | ALDOC | -1.71493 | 0.000001 | -0.91882 | 0.000001 |
| 115701 | ALPK2 | -2.10397 | 0.000001 | 1.840606 | 0.000001 |
| 60529 | ALX4 | -1.74233 | 0.000296 | -1.03257 | 0.000352 |
| 57463 | AMIGO1 | -2.53151 | 0.000001 | -1.14793 | 0.000001 |
| 154796 | AMOT | -1.01065 | 0.000001 | -2.91702 | 0.000001 |
| 272 | AMPD3 | -1.37878 | 0.000001 | -0.65539 | 0.001456 |
| 201283 | AMZ2P1 | -1.11783 | 4.04E-06 | -0.6766 | 0.000461 |
| 162282 | ANKFN1 | -2.60172 | 0.000001 | -3.01301 | 0.000001 |
| 23141 | ANKLE2 | 1.375155 | 0.000001 | 0.724846 | 0.000001 |
| 27063 | ANKRD1 | 2.254689 | 0.000001 | 4.47044 | 0.000001 |
| 55608 | ANKRD10 | 1.442095 | 0.000001 | 0.715206 | 0.000105 |
| 1.01E+08 | ANKRD10-IT1 | 1.149406 | 0.000754 | 1.326741 | 0.008264 |
| 341405 | ANKRD33 | 1.31501 | 0.003639 | -2.87736 | 0.000001 |
| 651746 | ANKRD33B | -2.03372 | 0.000001 | -3.03394 | 0.000001 |
| 284615 | ANKRD34A | -2.37222 | 0.000001 | -0.80704 | 0.003274 |
| 148741 | ANKRD35 | -2.55574 | 0.000001 | -2.14336 | 0.000001 |
| 91526 | ANKRD44 | -1.54878 | 2.24E-06 | 1.809655 | 0.000001 |
| 441869 | ANKRD65 | -1.99346 | 0.000752 | -1.9636 | 0.000001 |
| 54443 | ANLN | 1.513751 | 0.000001 | -1.64322 | 0.000001 |
| 121601 | ANO4 | 3.778998 | 0.000001 | -1.06554 | 0.000277 |
| 57719 | ANO8 | -1.32602 | 0.000001 | -0.67755 | 2.36E-05 |
| 290 | ANPEP | -1.29447 | 0.000001 | -0.6747 | 7.31E-05 |

| | | | | | |
|---|---|---|---|---|---|
| 306 | ANXA3 | 2.864748 | 0.000001 | -1.56229 | 0.000001 |
| 316 | AOX1 | -2.16478 | 0.000001 | -0.93247 | 0.000001 |
| 1173 | AP2M1 | -1.00074 | 0.000001 | 0.750677 | 0.000001 |
| 10307 | APBB3 | -1.07205 | 4.73E-05 | 0.631686 | 0.002981 |
| 147495 | APCDD1 | -3.29448 | 0.000001 | -4.46242 | 0.000001 |
| 149773 | APCDD1L-DT | 1.883864 | 0.000001 | 2.089306 | 0.000001 |
| 27350 | APOBEC3C | -1.79158 | 0.000001 | -1.10122 | 0.000001 |
| 140564 | APOBEC3D | -1.3294 | 7.68E-05 | -1.22386 | 0.003174 |
| 200316 | APOBEC3F | -2.86727 | 0.000001 | -0.92292 | 0.000365 |
| 60489 | APOBEC3G | -3.78186 | 0.000001 | -1.24983 | 0.000001 |
| 8542 | APOL1 | -1.91346 | 0.000001 | -1.57566 | 8.3E-06 |
| 80833 | APOL3 | -2.24849 | 0.000001 | -2.89567 | 0.000001 |
| 80830 | APOL6 | -2.90794 | 0.000001 | -0.77442 | 0.000001 |
| 358 | AQP1 | 4.536154 | 0.000001 | 2.256335 | 0.000001 |
| 364 | AQP7 | -2.33984 | 0.002672 | -2.83827 | 0.000001 |
| 116984 | ARAP2 | 3.042222 | 0.000001 | 2.451146 | 0.000001 |
| 392 | ARHGAP1 | -1.13143 | 0.000001 | 0.663718 | 0.000001 |
| 9824 | ARHGAP11A | 1.054456 | 0.000001 | -0.83784 | 0.000001 |
| 57569 | ARHGAP20 | 1.241734 | 1.59E-05 | -0.63281 | 0.01975 |
| 83478 | ARHGAP24 | -1.14085 | 0.000284 | -1.66804 | 0.000001 |
| 79822 | ARHGAP28 | -2.98864 | 0.000001 | -1.98145 | 0.000001 |
| 395 | ARHGAP6 | 1.071922 | 0.000158 | -3.86037 | 0.000001 |
| 397 | ARHGDIB | 2.375274 | 1.2E-05 | 1.862823 | 0.000001 |
| 50650 | ARHGEF3 | -1.0512 | 0.000001 | -1.56661 | 0.000001 |
| 50649 | ARHGEF4 | -2.22676 | 0.000001 | -1.21072 | 0.000001 |
| 9459 | ARHGEF6 | -1.64759 | 0.000001 | -1.08238 | 0.000001 |
| 406 | ARNTL | 1.719731 | 0.000001 | 0.963357 | 0.000001 |
| 348110 | ARPIN | -1.93811 | 0.000001 | -0.74799 | 0.000001 |
| 409 | ARRB2 | -1.34123 | 0.000001 | -0.83305 | 0.000001 |

| | | | | | |
|---|---|---|---|---|---|
| 27106 | ARRDC2 | -1.0522 | 0.000001 | -0.75159 | 0.000001 |
| 57412 | AS3MT | -2.3699 | 0.000001 | -1.1886 | 0.000001 |
| 653308 | ASAH2B | 1.305646 | 0.000001 | 0.700201 | 4.33E-05 |
| 55616 | ASAP3 | -2.62492 | 0.000001 | -0.63867 | 1.89E-06 |
| 55723 | ASF1B | 1.668592 | 0.000001 | -1.42509 | 0.000001 |
| 41 | ASIC1 | -2.43497 | 0.000001 | -1.87453 | 0.000001 |
| 435 | ASL | -1.5974 | 0.000001 | 0.62219 | 2.73E-06 |
| 440 | ASNS | -2.14625 | 0.000001 | 2.599758 | 0.000001 |
| 443 | ASPA | -2.04144 | 0.000001 | -1.8853 | 0.000248 |
| 54829 | ASPN | 1.010023 | 0.00351 | 2.457895 | 0.000001 |
| 23245 | ASTN2 | -1.29804 | 0.000142 | 1.013596 | 0.000001 |
| 29028 | ATAD2 | 1.691491 | 0.000001 | -0.92461 | 0.000001 |
| 79915 | ATAD5 | 1.822229 | 0.000001 | -1.17222 | 1.87E-06 |
| 467 | ATF3 | 1.833793 | 0.000001 | 0.973161 | 0.00022 |
| 23130 | ATG2A | -1.58527 | 0.000001 | -0.61505 | 0.000001 |
| 481 | ATP1B1 | 1.573837 | 0.000001 | 2.529033 | 0.000001 |
| 91419 | ATP23 | -1.41767 | 9.31E-06 | -1.12616 | 1.57E-05 |
| 490 | ATP2B1 | 1.085909 | 0.000001 | -0.87681 | 0.000001 |
| 23545 | ATP6V0A2 | 1.422999 | 0.000001 | 1.219787 | 0.000001 |
| 155066 | ATP6V0E2 | -1.57462 | 0.000001 | -0.87432 | 0.000001 |
| 9550 | ATP6V1G1 | 1.586593 | 0.000001 | 0.758993 | 0.000001 |
| 148229 | ATP8B3 | -1.95283 | 0.000001 | -1.18168 | 0.001329 |
| 79895 | ATP8B4 | -3.00202 | 0.000001 | -1.96642 | 1.42E-06 |
| 6790 | AURKA | 1.257109 | 0.000001 | -1.2102 | 0.000001 |
| 8705 | B3GALT4 | -2.539 | 0.000001 | -1.45916 | 0.000001 |
| 338707 | B4GALNT4 | -1.70793 | 0.001844 | -2.1393 | 0.000001 |
| 8703 | B4GALT3 | 1.213178 | 0.000001 | 0.661737 | 0.000001 |
| 11041 | B4GAT1 | -1.72346 | 0.000001 | -0.99366 | 0.000001 |
| 79870 | BAALC | -1.79619 | 0.000001 | -1.47194 | 0.000121 |

| | | | | | |
|---|---|---|---|---|---|
| 60468 | BACH2 | -2.99768 | 6.5E-06 | 1.751167 | 0.000001 |
| 440465 | BAIAP2-DT | -3.1115 | 0.000001 | -1.7783 | 0.000001 |
| 80115 | BAIAP2L2 | 2.091388 | 0.000001 | -1.30293 | 0.001768 |
| 578 | BAK1 | 1.162 | 0.000001 | 0.601981 | 2.49E-06 |
| 116071 | BATF2 | -3.31994 | 0.000001 | -3.26616 | 0.000001 |
| 27113 | BBC3 | -1.22049 | 0.000001 | -0.63843 | 0.000001 |
| 8537 | BCAS1 | -2.43512 | 0.000131 | -2.07503 | 0.000001 |
| 1.01E+08 | BCL2L2-PABPN1 | 3.128138 | 0.000001 | -1.21729 | 5.77E-05 |
| 618 | BCYRN1 | 2.254783 | 0.000001 | 0.920454 | 0.048991 |
| 624 | BDKRB2 | -2.39402 | 0.000001 | -2.34022 | 0.000001 |
| 79656 | BEND5 | -2.31317 | 0.000483 | -1.07998 | 0.006604 |
| 27319 | BHLHE22 | -2.9795 | 0.000001 | -2.39296 | 1.22E-06 |
| 330 | BIRC3 | -1.221 | 0.000506 | -2.17577 | 0.000001 |
| 641 | BLM | 1.604891 | 0.000427 | -0.82008 | 0.00035 |
| 90427 | BMF | -3.77436 | 0.000001 | 0.991424 | 2.07E-05 |
| 650 | BMP2 | 2.436678 | 0.000001 | -3.75606 | 0.000001 |
| 652 | BMP4 | 2.5574 | 0.000181 | -2.75186 | 0.000001 |
| 658 | BMPR1B | 3.511105 | 0.000001 | 2.913335 | 0.000001 |
| 91653 | BOC | -1.82051 | 0.000001 | -0.78665 | 0.000001 |
| 666 | BOK | -1.39665 | 0.000001 | 1.24706 | 0.000001 |
| 23246 | BOP1 | 1.000017 | 0.000001 | 0.747209 | 0.000001 |
| 669 | BPGM | 1.322218 | 0.000001 | 1.238768 | 0.000001 |
| 266655 | BRD3OS | -1.38232 | 0.000001 | -0.61391 | 0.000001 |
| 57795 | BRINP2 | -2.79972 | 0.000001 | -2.68868 | 2.06E-06 |
| 55643 | BTBD2 | -1.4016 | 0.000001 | -0.99825 | 0.000001 |
| 11119 | BTN3A1 | -3.56311 | 0.000001 | -0.72212 | 0.000001 |
| 11118 | BTN3A2 | -1.87448 | 0.000001 | -1.36982 | 0.000001 |
| 10384 | BTN3A3 | -2.6577 | 0.000001 | -0.80686 | 1.18E-06 |
| 699 | BUB1 | 1.008854 | 0.000579 | -1.11868 | 0.000001 |

| | | | | | |
|---|---|---|---|---|---|
| 701 | BUB1B | 1.25949 | 2.44E-05 | -1.30473 | 0.000001 |
| 414152 | C10orf105 | -5.64042 | 0.000001 | -3.56161 | 0.000001 |
| 80007 | C10orf88 | 1.402196 | 0.000001 | 0.605378 | 4.54E-05 |
| 219833 | C11orf45 | -1.43358 | 3.34E-06 | -1.67212 | 2.64E-05 |
| 399947 | C11orf87 | -2.02056 | 0.000001 | -2.60908 | 0.000001 |
| 728568 | C12orf73 | 1.006105 | 8.28E-06 | 0.878983 | 6.27E-05 |
| 122525 | C14orf28 | -1.96986 | 0.000001 | 0.703357 | 2.41E-06 |
| 60686 | C14orf93 | -3.21506 | 0.000001 | -0.82016 | 0.000001 |
| 404550 | C16orf74 | -2.34764 | 7.85E-05 | -2.10125 | 0.000001 |
| 388284 | C16orf86 | -3.15379 | 0.000001 | -1.33236 | 1.26E-05 |
| 388272 | C16orf87 | 1.348474 | 0.000001 | 0.937394 | 0.000001 |
| 388327 | C17orf100 | -1.58283 | 0.000755 | 0.603435 | 0.046755 |
| 148423 | C1orf52 | 1.409345 | 0.000001 | 0.734748 | 0.000001 |
| 54964 | C1orf56 | -2.72662 | 0.000001 | -0.98709 | 2.31E-06 |
| 114897 | C1QTNF1 | -1.07398 | 0.000001 | -1.7823 | 0.000001 |
| 114898 | C1QTNF2 | 1.029767 | 0.000584 | -3.00308 | 0.000001 |
| 54058 | C21orf58 | -2.68396 | 0.000001 | -0.61935 | 0.019893 |
| 26005 | C2CD3 | 1.182557 | 1.29E-06 | 1.241506 | 0.000001 |
| 29798 | C2orf27A | -2.62711 | 0.000001 | 1.148276 | 0.000001 |
| 388963 | C2orf81 | -2.02263 | 0.000001 | -0.65064 | 0.008342 |
| 51161 | C3orf18 | -1.34258 | 0.000001 | -1.51186 | 0.000001 |
| 401097 | C3orf80 | 2.699543 | 0.000001 | 1.107981 | 0.001421 |
| 132989 | C4orf36 | -1.19016 | 0.009935 | -1.22727 | 0.000195 |
| 727 | C5 | -2.58533 | 0.000001 | -1.03622 | 0.000001 |
| 27202 | C5AR2 | 2.36085 | 0.002809 | 1.556163 | 0.009748 |
| 402573 | C7orf61 | 2.524554 | 0.000238 | -1.21146 | 0.041727 |
| 116328 | C8orf34 | -1.53566 | 0.000001 | 1.966012 | 0.000001 |
| 771 | CA12 | -1.38925 | 0.000001 | -1.16846 | 0.000001 |
| 91768 | CABLES1 | -1.52038 | 0.000001 | -2.11629 | 0.000001 |

| | | | | | |
|---|---|---|---|---|---|
| 8913 | CACNA1G | 1.078083 | 2.19E-05 | 1.271865 | 0.000001 |
| 93589 | CACNA2D4 | -2.12311 | 3.52E-06 | -0.89965 | 0.041337 |
| 8618 | CADPS | -1.15074 | 0.001073 | -1.72761 | 0.000001 |
| 794 | CALB2 | 1.086353 | 0.003179 | 1.34969 | 0.000001 |
| 57658 | CALCOCO1 | -2.641 | 0.000001 | 0.859278 | 0.000001 |
| 800 | CALD1 | -1.18921 | 0.000001 | 1.723383 | 0.000001 |
| 254228 | CALHM5 | -1.8835 | 0.000001 | 1.226004 | 0.000001 |
| 91860 | CALML4 | 2.964421 | 0.000001 | -1.5789 | 0.000286 |
| 23066 | CAND2 | -1.38008 | 0.003172 | -1.53636 | 0.000001 |
| 114769 | CARD16 | -1.79098 | 0.000001 | -2.77262 | 0.000001 |
| 23589 | CARHSP1 | -1.95428 | 0.000001 | -0.60881 | 0.000001 |
| 255082 | CASC2 | -3.0048 | 0.000001 | -0.8125 | 0.016669 |
| 834 | CASP1 | -2.92438 | 0.000001 | -3.0678 | 0.000001 |
| 843 | CASP10 | -2.49915 | 0.000001 | -1.28622 | 0.00285 |
| 1.01E+08 | CASP12 | 2.649487 | 0.000001 | -4.16954 | 0.000001 |
| 847 | CAT | -2.41782 | 0.000001 | -1.69507 | 0.000001 |
| 8436 | CAVIN2 | -1.85689 | 0.000001 | -1.91236 | 0.000001 |
| 112464 | CAVIN3 | -1.24038 | 0.000001 | -0.76271 | 5.81E-06 |
| 347273 | CAVIN4 | 1.288328 | 0.000001 | 2.122031 | 0.000001 |
| 867 | CBL | 1.652455 | 0.000001 | 0.697477 | 2.59E-06 |
| 868 | CBLB | 1.009281 | 0.001177 | 0.603883 | 0.003228 |
| 1.01E+08 | CBR3-AS1 | -1.79188 | 0.000001 | -0.75943 | 0.001593 |
| 875 | CBS | -1.52411 | 2.45E-05 | 1.338835 | 0.000001 |
| 23492 | CBX7 | -2.66746 | 0.000001 | -0.78254 | 0.000001 |
| 79839 | CCDC102B | -2.16141 | 0.000001 | -3.25289 | 0.000001 |
| 84317 | CCDC115 | -1.8366 | 0.000001 | -0.69883 | 0.000001 |
| 79635 | CCDC121 | -2.05346 | 0.000001 | -1.14302 | 9.42E-05 |
| 64753 | CCDC136 | -1.05752 | 0.000289 | -1.14164 | 6.52E-05 |
| 9720 | CCDC144A | -1.25013 | 0.00063 | 1.37958 | 2.27E-06 |

| | | | | | |
|---|---|---|---|---|---|
| 387856 | CCDC184 | -1.68309 | 0.002154 | -1.10445 | 0.017752 |
| 57577 | CCDC191 | -1.88407 | 0.000001 | -0.92389 | 0.00022 |
| 79140 | CCDC28B | -2.12233 | 0.000001 | -0.77948 | 0.000805 |
| 91057 | CCDC34 | 1.521913 | 0.000001 | -0.75538 | 0.000001 |
| 26112 | CCDC69 | -1.06687 | 0.000001 | -1.65206 | 0.000001 |
| 90557 | CCDC74A | -1.14809 | 0.000001 | -0.9016 | 0.000001 |
| 91409 | CCDC74B | -1.1035 | 1.31E-05 | -1.53543 | 0.000001 |
| 11007 | CCDC85B | -1.55355 | 0.000001 | -1.35182 | 0.000001 |
| 388115 | CCDC9B | -1.73668 | 0.000001 | -0.67092 | 3.52E-05 |
| 3491 | CCN1 | -1.0458 | 3.01E-06 | 1.49888 | 0.000001 |
| 4856 | CCN3 | 2.649979 | 0.000001 | -4.07521 | 0.000001 |
| 8840 | CCN4 | -1.81321 | 0.00196 | 1.437524 | 8.99E-06 |
| 8839 | CCN5 | -1.12125 | 0.000001 | -0.77418 | 3.14E-06 |
| 891 | CCNB1 | 1.33707 | 0.000001 | -0.78081 | 0.000001 |
| 57124 | CD248 | -1.924 | 0.000001 | -1.29084 | 0.000001 |
| 80381 | CD276 | 1.151527 | 0.000001 | -0.81046 | 0.000001 |
| 9936 | CD302 | -2.01332 | 0.000001 | -1.55875 | 0.000001 |
| 948 | CD36 | 2.183146 | 0.000001 | -3.75347 | 0.000001 |
| 958 | CD40 | -1.92227 | 0.000001 | -1.1476 | 0.000212 |
| 1604 | CD55 | 1.906417 | 0.000001 | 0.785704 | 2.28E-05 |
| 1.05E+08 | CD63-AS1 | -1.57475 | 0.000001 | -0.99732 | 0.000131 |
| 3732 | CD82 | -1.38189 | 0.000001 | -1.03397 | 0.000001 |
| 9308 | CD83 | 1.114796 | 0.000854 | -3.01995 | 0.000001 |
| 8555 | CDC14B | -1.27999 | 0.000001 | -1.28979 | 0.000001 |
| 991 | CDC20 | 1.823671 | 0.000001 | -1.12524 | 0.000001 |
| 993 | CDC25A | 2.815911 | 0.000001 | -0.9515 | 0.000001 |
| 148170 | CDC42EP5 | -1.45978 | 0.000001 | -0.71406 | 0.000001 |
| 990 | CDC6 | 2.938539 | 0.000001 | -0.83401 | 0.000001 |
| 8317 | CDC7 | 1.336371 | 1.35E-06 | -0.84488 | 2.55E-06 |

| | | | | | |
|---|---|---|---|---|---|
| 113130 | CDCA5 | 1.910988 | 0.000001 | -1.32867 | 0.000001 |
| 55143 | CDCA8 | 1.428758 | 0.000001 | -1.04172 | 0.000001 |
| 64866 | CDCP1 | 2.641974 | 0.000001 | -2.55239 | 0.000001 |
| 1022 | CDK7 | 1.148775 | 0.000001 | 0.683526 | 1.68E-06 |
| 8999 | CDKL2 | 2.062144 | 0.000001 | -2.21693 | 0.000001 |
| 1028 | CDKN1C | -4.20589 | 0.000001 | -1.07915 | 0.000001 |
| 1030 | CDKN2B | -1.31294 | 0.000001 | 1.736581 | 0.000001 |
| 1031 | CDKN2C | -2.1981 | 0.000001 | -1.77653 | 0.000001 |
| 1.17E+08 | CDRT15P4 | -3.58571 | 2.97E-06 | 1.161213 | 0.025077 |
| 81620 | CDT1 | 2.52959 | 0.000001 | -1.27877 | 0.000001 |
| 1.02E+08 | CEBPB-AS1 | -2.30692 | 0.001416 | 0.970169 | 0.022314 |
| 1054 | CEBPG | 1.657215 | 0.000001 | 1.129792 | 0.000001 |
| 57214 | CEMIP | -1.86363 | 6.8E-05 | -3.1411 | 0.000001 |
| 1058 | CENPA | 1.081889 | 0.00602 | -1.23192 | 0.000001 |
| 64105 | CENPK | 1.660537 | 0.000001 | -0.82863 | 1.48E-05 |
| 55839 | CENPN | 1.087996 | 0.000001 | 0.615198 | 4.35E-05 |
| 79682 | CENPU | 1.266978 | 1.51E-05 | -1.28467 | 0.000001 |
| 201254 | CENPX | 1.056913 | 0.000001 | -0.88448 | 0.000001 |
| 145508 | CEP128 | 1.151932 | 5.45E-05 | -1.05024 | 1.18E-06 |
| 55165 | CEP55 | 1.557513 | 0.000001 | -1.25105 | 0.000001 |
| 9702 | CEP57 | -1.1041 | 0.000001 | -0.71893 | 0.000001 |
| 80321 | CEP70 | -2.58779 | 0.000001 | -0.81572 | 7.93E-05 |
| 1E+08 | CERNA1 | -2.47156 | 6.75E-06 | -0.87165 | 0.04577 |
| 144535 | CFAP54 | 1.571576 | 0.000001 | 0.715282 | 0.045484 |
| 629 | CFB | -1.25612 | 0.000001 | -0.92988 | 0.000001 |
| 79094 | CHAC1 | -1.44257 | 0.001017 | 1.565355 | 0.000119 |
| 150356 | CHADL | -1.83628 | 4.57E-05 | -1.25166 | 0.000819 |
| 10036 | CHAF1A | 1.4788 | 0.000001 | -0.9782 | 0.000001 |
| 55636 | CHD7 | 2.04004 | 0.000001 | 1.42234 | 9.75E-06 |

| | | | | | |
|---|---|---|---|---|---|
| 55743 | CHFR | 1.36853 | 0.000001 | 0.677256 | 0.000001 |
| 1123 | CHN1 | 1.201863 | 0.000001 | 0.730314 | 0.000001 |
| 1129 | CHRM2 | -2.17488 | 0.000725 | -2.7528 | 0.000001 |
| 55584 | CHRNA9 | 2.698128 | 0.000001 | 5.330117 | 0.000001 |
| 113189 | CHST14 | -2.54137 | 0.000001 | -0.76321 | 0.000001 |
| 56548 | CHST7 | -2.19598 | 0.000001 | -0.81561 | 0.0001 |
| 337876 | CHSY3 | -2.13958 | 0.000001 | 0.948044 | 0.000001 |
| 27141 | CIDEB | -2.69627 | 0.000001 | -2.40317 | 0.000001 |
| 152302 | CIDECP1 | 1.047266 | 2.5E-06 | 0.617395 | 0.00116 |
| 8483 | CILP | -1.77779 | 9.54E-05 | 1.626081 | 0.000001 |
| 1164 | CKS2 | 2.062335 | 0.000001 | -0.86723 | 1.14E-06 |
| 122616 | CLBA1 | -1.77338 | 0.000001 | -0.62897 | 2.92E-05 |
| 9635 | CLCA2 | -1.50414 | 0.000001 | -1.48895 | 0.000001 |
| 23529 | CLCF1 | 1.195149 | 0.000001 | 1.29377 | 0.000001 |
| 5010 | CLDN11 | -1.7083 | 0.000001 | -4.71865 | 0.000001 |
| 137075 | CLDN23 | -1.9808 | 0.000001 | -0.9932 | 0.000001 |
| 9022 | CLIC3 | -2.78838 | 0.000001 | 2.799204 | 0.000001 |
| 63967 | CLSPN | 2.340919 | 0.000001 | -1.275 | 0.000001 |
| 64084 | CLSTN2 | 2.522952 | 0.000001 | 1.401363 | 0.000001 |
| 8218 | CLTCL1 | -1.32546 | 2.38E-06 | 1.946251 | 0.000001 |
| 134147 | CMBL | -1.59483 | 0.000001 | -1.91152 | 0.000001 |
| 56942 | CMC2 | 1.253245 | 0.000001 | 0.822296 | 0.000001 |
| 149111 | CNIH3 | 1.892974 | 5.39E-05 | -1.04203 | 0.000001 |
| 1.03E+08 | CNIH3-AS2 | 2.177128 | 0.000001 | -1.19745 | 0.023796 |
| 1265 | CNN2 | -1.53114 | 0.000001 | 1.179511 | 0.000001 |
| 26507 | CNNM1 | 2.074985 | 0.000001 | -1.35979 | 0.000001 |
| 163882 | CNST | 1.814253 | 0.000001 | 0.689039 | 0.000001 |
| 1300 | COL10A1 | 1.374885 | 0.000001 | 5.390615 | 0.000001 |
| 1301 | COL11A1 | -1.45127 | 0.000001 | 1.929037 | 1.14E-06 |

| | | | | | |
|---|---|---|---|---|---|
| 1277 | COL1A1 | -1.56287 | 0.000001 | 1.636121 | 0.000001 |
| 255631 | COL24A1 | -1.38513 | 0.001081 | 1.696087 | 0.000001 |
| 1287 | COL4A5 | -2.52097 | 0.000001 | -1.44677 | 0.000001 |
| 1289 | COL5A1 | -1.04363 | 0.000001 | 2.278971 | 0.000001 |
| 1290 | COL5A2 | -1.16792 | 0.000001 | 0.930962 | 0.000001 |
| 1293 | COL6A3 | -1.15527 | 0.000001 | 0.935272 | 0.000001 |
| 1295 | COL8A1 | -1.04684 | 0.000001 | 3.286536 | 0.000001 |
| 81035 | COLEC12 | -1.30022 | 0.000001 | -3.79139 | 0.000001 |
| 118881 | COMTD1 | -1.78648 | 0.000001 | -1.09373 | 0.000001 |
| 56997 | COQ8A | -3.80719 | 0.000001 | -0.62243 | 7.36E-05 |
| 79934 | COQ8B | -1.44582 | 0.000001 | -0.64023 | 0.000001 |
| 1.02E+08 | COSMOC | -1.76331 | 0.000001 | -1.3527 | 0.000466 |
| 64506 | CPEB1 | 1.406148 | 0.000001 | 0.823379 | 0.000001 |
| 1368 | CPM | -2.13203 | 0.000001 | 0.889484 | 8.31E-05 |
| 8895 | CPNE3 | -1.05969 | 0.000001 | -0.93995 | 0.000001 |
| 144402 | CPNE8 | 1.577809 | 0.000001 | -1.35722 | 0.000001 |
| 1374 | CPT1A | 1.281888 | 0.000001 | -0.84725 | 0.000001 |
| 56265 | CPXM1 | -1.16414 | 0.000145 | -1.35714 | 0.000001 |
| 57482 | CRACD | 1.327942 | 0.001862 | 0.643983 | 0.000001 |
| 90993 | CREB3L1 | -1.14191 | 0.000001 | 1.060154 | 0.000001 |
| 153222 | CREBRF | -2.12566 | 0.000001 | -0.67362 | 0.000001 |
| 1396 | CRIP1 | 5.254456 | 0.000001 | -1.5131 | 0.000001 |
| 9419 | CRIPT | -1.94785 | 0.000001 | 0.637595 | 0.000001 |
| 83716 | CRISPLD2 | -1.36998 | 0.000001 | 2.256117 | 0.000001 |
| 9696 | CROCC | -1.97066 | 0.000001 | 1.138103 | 0.000001 |
| 1407 | CRY1 | 1.718577 | 0.000001 | 1.110584 | 0.000001 |
| 51084 | CRYL1 | -2.10594 | 0.000001 | -1.82486 | 0.000001 |
| 730102 | CRYZL2P | -1.03921 | 0.000001 | -1.29051 | 0.000001 |
| 27254 | CSDC2 | -2.86222 | 0.000001 | 0.998928 | 0.000001 |

| | | | | | |
|---|---|---|---|---|---|
| 55790 | CSGALNACT1 | -1.42772 | 4.59E-06 | -0.97603 | 0.036714 |
| 55454 | CSGALNACT2 | 1.065074 | 0.000001 | 0.793651 | 0.000001 |
| 64651 | CSRNP1 | 1.753504 | 0.000001 | 0.616964 | 9.78E-06 |
| 115908 | CTHRC1 | -2.04793 | 0.000001 | 1.159977 | 0.000001 |
| 1503 | CTPS1 | 1.14931 | 0.000001 | 2.189432 | 0.000001 |
| 56474 | CTPS2 | -1.65901 | 0.000001 | -0.77084 | 0.000001 |
| 1519 | CTSO | -1.97213 | 0.000001 | -0.62197 | 0.000001 |
| 2919 | CXCL1 | -2.00749 | 0.000419 | -2.272 | 0.000001 |
| 6387 | CXCL12 | -1.98368 | 0.000001 | 0.649026 | 0.000308 |
| 58191 | CXCL16 | -2.30015 | 0.000001 | -1.01016 | 0.00811 |
| 6372 | CXCL6 | -3.02553 | 5.15E-06 | -1.17626 | 0.009745 |
| 7852 | CXCR4 | 3.304831 | 0.000001 | 1.609817 | 0.007593 |
| 51523 | CXXC5 | -1.73354 | 0.000001 | 0.86292 | 0.000001 |
| 54205 | CYCS | 1.326282 | 0.000001 | 0.657021 | 1.41E-05 |
| 1591 | CYP24A1 | -4.56252 | 0.000001 | -2.40969 | 2.62E-06 |
| 339761 | CYP27C1 | 2.909015 | 0.000001 | 0.907498 | 0.000001 |
| 113612 | CYP2U1 | -1.3138 | 0.000001 | -0.96245 | 0.000001 |
| 1551 | CYP3A7 | 1.684711 | 0.007686 | -2.17577 | 0.000132 |
| 9420 | CYP7B1 | -1.8716 | 0.000732 | -1.6818 | 0.000001 |
| 116159 | CYYR1 | -1.08939 | 0.000001 | -1.72579 | 0.000001 |
| 51339 | DACT1 | -1.55625 | 0.000725 | 3.272879 | 0.000001 |
| 147906 | DACT3 | -1.2408 | 1.05E-05 | 1.101888 | 0.000001 |
| 57291 | DANCR | 1.435961 | 3.76E-05 | -0.83908 | 3.23E-06 |
| 285761 | DCBLD1 | 1.345651 | 0.000001 | 1.28544 | 0.000001 |
| 9201 | DCLK1 | 1.553347 | 0.000001 | -1.89391 | 0.000001 |
| 166614 | DCLK2 | 1.167146 | 0.000001 | 2.113768 | 0.000001 |
| 1643 | DDB2 | -1.30058 | 0.000001 | -1.57773 | 0.000001 |
| 220042 | DDIAS | 2.249297 | 0.000001 | -0.91811 | 3.03E-05 |
| 9188 | DDX21 | 1.896545 | 0.000001 | 0.800434 | 0.000001 |

| | | | | | |
|---|---|---|---|---|---|
| 55601 | DDX60 | -3.44822 | 0.000001 | -0.94513 | 1.67E-06 |
| 50514 | DELEC1 | 4.064947 | 0.000001 | 1.580708 | 0.001579 |
| 79961 | DENND2D | -1.24492 | 0.000169 | -1.33204 | 2.12E-05 |
| 22898 | DENND3 | -2.12763 | 0.000001 | -1.33312 | 0.000001 |
| 55635 | DEPDC1 | 1.094361 | 0.001977 | -1.09709 | 0.000001 |
| 55789 | DEPDC1B | 1.835746 | 0.003961 | -1.26063 | 0.001472 |
| 91614 | DEPDC7 | -1.39433 | 0.000001 | -0.86012 | 0.001566 |
| 11067 | DEPP1 | -4.34864 | 0.000001 | -1.48355 | 0.001308 |
| 54487 | DGCR8 | 1.170475 | 0.000001 | 0.790755 | 0.000001 |
| 9162 | DGKI | 2.312452 | 0.000001 | 1.673719 | 0.000001 |
| 80017 | DGLUCY | -2.15765 | 0.000001 | -0.75973 | 0.000001 |
| 115817 | DHRS1 | -2.76356 | 0.000001 | -0.67831 | 0.000001 |
| 9704 | DHX34 | 1.463659 | 0.000001 | 0.64505 | 0.000001 |
| 79132 | DHX58 | -4.36448 | 0.000001 | -0.85248 | 0.000001 |
| 1734 | DIO2 | -1.33795 | 0.003386 | 2.832826 | 0.000001 |
| 388650 | DIPK1A | -1.93452 | 0.000001 | 0.860954 | 0.000001 |
| 115752 | DIS3L | -1.19606 | 0.000001 | -0.85573 | 0.000001 |
| 85458 | DIXDC1 | -1.94287 | 0.000001 | 1.210323 | 0.000001 |
| 22943 | DKK1 | 1.064385 | 0.000001 | -1.57169 | 0.000001 |
| 28514 | DLL1 | -1.07368 | 0.002418 | -1.4195 | 0.001056 |
| 1756 | DMD | 1.735603 | 1.4E-05 | 0.957722 | 0.000001 |
| 93099 | DMKN | -1.22659 | 0.000001 | -0.64804 | 0.000001 |
| 1763 | DNA2 | 1.211576 | 0.000001 | -0.93466 | 0.00012 |
| 23639 | DNAAF11 | -1.30195 | 7.78E-05 | -1.71796 | 0.000001 |
| 1767 | DNAH5 | -1.27265 | 0.000199 | 1.664908 | 0.000001 |
| 25822 | DNAJB5 | -1.47534 | 0.000001 | 1.474999 | 0.000001 |
| 23234 | DNAJC9 | 1.893842 | 0.000001 | -0.77028 | 0.000001 |
| 7802 | DNALI1 | 2.62568 | 0.000001 | -2.04031 | 0.000001 |
| 1.01E+08 | DNM3OS | -1.59334 | 0.000001 | 2.713599 | 0.000001 |

| | | | | | |
|---|---|---|---|---|---|
| 55619 | DOCK10 | 1.99612 | 0.000001 | 0.76944 | 0.000001 |
| 1.01E+08 | DOCK9-DT | -2.22974 | 0.003391 | 1.235382 | 0.026448 |
| 79930 | DOK3 | 1.422589 | 0.000001 | 0.82577 | 0.042793 |
| 55816 | DOK5 | 2.014019 | 0.000001 | -0.64718 | 0.008394 |
| 22845 | DOLK | -1.24637 | 0.000001 | -0.67791 | 0.000001 |
| 1802 | DPH2 | 1.355866 | 0.000001 | 0.80635 | 0.000001 |
| 349152 | DPY19L2P2 | 1.700785 | 0.000001 | -1.83696 | 0.000001 |
| 1806 | DPYD | -3.58614 | 0.000001 | -0.60477 | 0.001463 |
| 1809 | DPYSL3 | 2.021592 | 0.000001 | 0.867446 | 0.000001 |
| 1812 | DRD1 | 2.217316 | 0.00684 | 1.415278 | 0.020431 |
| 29940 | DSE | -1.72559 | 0.000001 | 1.27737 | 0.000001 |
| 1832 | DSP | -1.01707 | 0.000001 | 2.361081 | 0.000001 |
| 51514 | DTL | 3.322345 | 0.000001 | -1.23728 | 0.000001 |
| 1843 | DUSP1 | 1.679847 | 0.000001 | 0.899411 | 0.000001 |
| 1846 | DUSP4 | -1.04522 | 0.000443 | -2.07145 | 0.000001 |
| 1848 | DUSP6 | 1.120834 | 0.000001 | -1.3849 | 0.000001 |
| 1780 | DYNC1I1 | -1.6082 | 0.001613 | 1.06856 | 0.014229 |
| 1.07E+08 | DYNC1LI2-DT | -2.19479 | 0.001431 | 0.891918 | 0.033604 |
| 51626 | DYNC2LI1 | -1.61292 | 0.000001 | -0.73277 | 1.14E-05 |
| 199221 | DZIP1L | -1.21164 | 1.41E-06 | 1.116112 | 0.000001 |
| 1869 | E2F1 | 1.576582 | 0.000001 | -1.07155 | 0.000001 |
| 85403 | EAF1 | 1.761354 | 0.000001 | 0.74834 | 0.000001 |
| 253738 | EBF3 | -1.54109 | 0.000001 | -1.29648 | 0.000001 |
| 57593 | EBF4 | -1.5627 | 0.000649 | -0.67261 | 0.001352 |
| 10682 | EBP | -1.35807 | 0.000001 | -0.7155 | 0.000001 |
| 1891 | ECH1 | -1.03758 | 0.000001 | -0.9381 | 0.000001 |
| 1842 | ECM2 | -1.19982 | 0.000001 | 1.064006 | 0.000001 |
| 1894 | ECT2 | 1.357619 | 0.000001 | -0.90324 | 0.000001 |
| 80820 | EEPD1 | -1.37836 | 1.86E-06 | -2.31778 | 0.000001 |

| | | | | | |
|---|---|---|---|---|---|
| 79645 | EFCAB1 | 1.516875 | 0.000111 | -1.43082 | 0.010019 |
| 22979 | EFR3B | 1.22008 | 0.000001 | 1.435604 | 0.000001 |
| 1959 | EGR2 | 1.869672 | 1.91E-06 | 2.019796 | 0.000001 |
| 1962 | EHHADH | -2.21233 | 0.000001 | -0.69404 | 0.000141 |
| 8669 | EIF3J | 1.396949 | 0.000001 | 0.940767 | 0.000001 |
| 392617 | ELFN1 | -1.50667 | 0.000001 | -1.5312 | 0.000001 |
| 80237 | ELL3 | -2.06993 | 3.26E-05 | -1.45826 | 0.002677 |
| 90187 | EMILIN3 | -2.7788 | 0.0002 | -5.06711 | 0.000001 |
| 1.02E+08 | EMSLR | 3.844553 | 3.37E-06 | -1.76022 | 0.002801 |
| 2018 | EMX2 | -1.76382 | 0.000001 | -1.01075 | 0.000001 |
| 196047 | EMX2OS | -2.908 | 0.000001 | -1.54594 | 0.000001 |
| 59084 | ENPP5 | -1.32106 | 0.000001 | -0.98772 | 0.025808 |
| 957 | ENTPD5 | -1.50562 | 0.000001 | 0.622379 | 0.000001 |
| 2035 | EPB41 | -1.14111 | 0.000001 | -0.68133 | 0.000001 |
| 2037 | EPB41L2 | -1.7519 | 0.000001 | -1.10148 | 0.000001 |
| 2051 | EPHB6 | -2.63409 | 0.000001 | -2.0762 | 0.000001 |
| 2053 | EPHX2 | -2.79929 | 3.48E-06 | -1.90286 | 3.02E-06 |
| 2057 | EPOR | -1.05168 | 0.000664 | -1.40795 | 0.000001 |
| 83481 | EPPK1 | 2.583596 | 0.000109 | 3.320415 | 0.000001 |
| 64787 | EPS8L2 | -1.52575 | 0.000001 | -0.65496 | 0.000001 |
| 54821 | ERCC6L | 2.41666 | 0.000001 | -1.36696 | 0.000001 |
| 2069 | EREG | 2.833711 | 4.2E-05 | -0.97221 | 0.001222 |
| 2078 | ERG | 3.265892 | 4.35E-06 | 4.545728 | 0.000001 |
| 11161 | ERG28 | -2.20967 | 0.000001 | -0.62263 | 0.000001 |
| 2081 | ERN1 | 1.580035 | 0.000001 | 1.099093 | 0.000001 |
| 11082 | ESM1 | 1.365254 | 0.000155 | 0.949521 | 0.00807 |
| 9700 | ESPL1 | 1.139006 | 0.000001 | -1.61311 | 0.000001 |
| 2114 | ETS2 | 1.506527 | 0.000001 | -1.45079 | 0.000001 |
| 2120 | ETV6 | -2.0873 | 0.000001 | 1.633663 | 0.000001 |

| | | | | | |
|---|---|---|---|---|---|
| 51513 | ETV7 | -2.97134 | 0.000001 | -2.01755 | 0.00022 |
| 84141 | EVA1A | 1.71835 | 0.000001 | 1.162772 | 0.000001 |
| 51466 | EVL | -1.5606 | 0.000001 | 0.763592 | 2.62E-05 |
| 2134 | EXTL1 | -3.60938 | 0.000001 | 3.87425 | 0.000001 |
| 2138 | EYA1 | -1.6547 | 6E-05 | -2.19402 | 0.000001 |
| 7430 | EZR | 2.858113 | 0.000001 | 0.717763 | 3.96E-06 |
| 2150 | F2RL1 | 1.862945 | 0.006606 | 2.137481 | 2.25E-05 |
| 2152 | F3 | 1.686765 | 0.000001 | 1.240293 | 0.000001 |
| 2170 | FABP3 | -2.42876 | 0.000001 | -2.05595 | 0.000001 |
| 3992 | FADS1 | -2.22956 | 0.000001 | -0.85626 | 0.000001 |
| 9415 | FADS2 | -1.7501 | 0.000001 | -0.84543 | 0.000001 |
| 3995 | FADS3 | -1.8303 | 0.000001 | 0.796038 | 1.39E-06 |
| 2184 | FAH | -1.36469 | 0.000001 | -1.66016 | 0.000001 |
| 151313 | FAHD2B | -2.37079 | 0.000172 | -0.62367 | 0.004673 |
| 729234 | FAHD2CP | -1.36096 | 0.002813 | -0.85275 | 0.008 |
| 23017 | FAIM2 | -1.23203 | 0.001591 | -1.1653 | 9.54E-06 |
| 284611 | FAM102B | 1.617727 | 0.000001 | -0.75864 | 0.000001 |
| 11170 | FAM107A | 1.691147 | 0.000732 | -2.48915 | 1.15E-05 |
| 1.02E+08 | FAM111A-DT | -1.43701 | 0.000001 | -1.28421 | 2.9E-05 |
| 374393 | FAM111B | 1.332887 | 4.37E-05 | -1.9204 | 0.000001 |
| 81558 | FAM117A | -2.65893 | 0.000001 | -0.65317 | 0.002629 |
| 9715 | FAM131B | -1.94288 | 2.1E-05 | 0.740218 | 0.001014 |
| 25854 | FAM149A | -1.3084 | 0.000001 | -1.33177 | 0.000001 |
| 83648 | FAM167A | 1.149257 | 0.000001 | -0.66616 | 0.003757 |
| 165215 | FAM171B | -1.45842 | 0.000001 | -1.34948 | 0.000001 |
| 389558 | FAM180A | 2.481642 | 0.000001 | -0.94339 | 1.13E-06 |
| 220382 | FAM181B | -2.40114 | 0.000536 | -1.69246 | 0.001461 |
| 54757 | FAM20A | -3.75959 | 0.000001 | -1.91995 | 0.000001 |
| 116151 | FAM210B | -2.6563 | 0.000001 | -0.68425 | 0.000001 |

| | | | | | |
|---|---|---|---|---|---|
| 132720 | FAM241A | 1.73168 | 0.000001 | 1.104531 | 0.000001 |
| 131583 | FAM43A | -2.24347 | 0.000001 | -1.49538 | 0.000001 |
| 286336 | FAM78A | -2.50281 | 0.000001 | -1.5293 | 4.16E-06 |
| 81610 | FAM83D | 2.194841 | 0.000001 | -1.23192 | 0.000001 |
| 644815 | FAM83G | 2.160408 | 0.000001 | 1.307774 | 0.000001 |
| 645332 | FAM86C2P | -1.9496 | 0.000001 | -1.08314 | 0.000457 |
| 348926 | FAM86EP | -1.51091 | 1.7E-06 | -0.92303 | 0.004308 |
| 2175 | FANCA | 1.790982 | 0.000001 | -0.77627 | 0.000402 |
| 2176 | FANCC | 1.002998 | 4.53E-06 | -1.10577 | 0.000001 |
| 2177 | FANCD2 | 1.072797 | 0.000316 | -1.19392 | 0.000001 |
| 355 | FAS | 1.365791 | 0.000001 | -0.86228 | 0.000001 |
| 10826 | FAXDC2 | -3.57176 | 0.000001 | -2.05676 | 0.000001 |
| 2200 | FBN1 | -1.57137 | 0.000001 | 1.793292 | 0.000001 |
| 144699 | FBXL14 | 1.25038 | 0.000001 | -0.81471 | 1.45E-05 |
| 114907 | FBXO32 | -2.29628 | 0.000001 | 2.573438 | 0.000001 |
| 26271 | FBXO5 | 2.040782 | 0.000001 | -0.76733 | 0.000001 |
| 2237 | FEN1 | 1.887712 | 0.000001 | -0.76913 | 0.000001 |
| 654463 | FER1L6 | 1.56663 | 0.000001 | -1.7527 | 0.000625 |
| 2242 | FES | -2.17196 | 0.000001 | -1.38662 | 0.000001 |
| 9638 | FEZ1 | -1.59712 | 0.000001 | -0.71795 | 0.000001 |
| 121512 | FGD4 | -2.58879 | 0.000001 | -2.31258 | 0.000001 |
| 55785 | FGD6 | -2.01666 | 0.000001 | -0.68895 | 0.000682 |
| 2246 | FGF1 | 1.136952 | 0.000001 | 1.589044 | 0.000001 |
| 2255 | FGF10 | -1.86791 | 4.12E-05 | -2.57263 | 0.000001 |
| 2252 | FGF7 | -3.33511 | 0.000001 | -0.70191 | 9.07E-05 |
| 53834 | FGFRL1 | -1.34675 | 0.000001 | 0.871545 | 0.000001 |
| 2274 | FHL2 | 1.02192 | 0.000001 | 1.022014 | 0.000001 |
| 11153 | FICD | 1.76383 | 0.000001 | 0.947055 | 0.000001 |
| 63979 | FIGNL1 | 1.692529 | 0.000001 | -1.16132 | 0.000001 |

| | | | | | |
|---|---|---|---|---|---|
| 27145 | FILIP1 | -1.35542 | 0.006876 | 1.808023 | 0.000001 |
| 2323 | FLT3LG | -2.2154 | 0.000001 | -1.37006 | 0.000001 |
| 55640 | FLVCR2 | 2.351804 | 0.000001 | -0.95304 | 0.045246 |
| 2329 | FMO4 | -2.41314 | 0.000001 | -0.99596 | 2.62E-05 |
| 643988 | FNDC10 | -1.236 | 0.000001 | -1.21577 | 0.000001 |
| 79025 | FNDC11 | -2.0808 | 0.002111 | -1.50523 | 0.001635 |
| 64838 | FNDC4 | -1.08321 | 0.000001 | -1.1837 | 0.000001 |
| 2354 | FOSB | 3.351717 | 0.000001 | 1.505504 | 0.000425 |
| 8061 | FOSL1 | 2.722078 | 0.000001 | -0.93213 | 0.000001 |
| 2294 | FOXF1 | 1.709002 | 5.5E-05 | -1.31189 | 0.000001 |
| 10023 | FRAT1 | -4.52536 | 0.000001 | -1.47366 | 0.000001 |
| 84978 | FRMD5 | 1.905261 | 0.000001 | 1.694101 | 0.000001 |
| 145438 | FRMD6-AS1 | -1.85727 | 0.000001 | 1.977344 | 0.000001 |
| 25794 | FSCN2 | -1.7335 | 0.004997 | 1.350406 | 5.6E-06 |
| 10468 | FST | -2.25354 | 0.000001 | -1.41104 | 0.000001 |
| 2517 | FUCA1 | -1.60026 | 0.000001 | -0.94715 | 2.24E-06 |
| 645431 | FUT8-AS1 | -1.71722 | 0.000504 | 0.974616 | 0.001618 |
| 11211 | FZD10 | 2.107681 | 4.59E-05 | -1.5593 | 0.005493 |
| 7855 | FZD5 | 1.884034 | 0.000001 | -1.38607 | 2.59E-06 |
| 8325 | FZD8 | 1.80271 | 0.000001 | 3.287982 | 0.000001 |
| 23710 | GABARAPL1 | -2.62425 | 0.000001 | 0.690266 | 0.000001 |
| 51083 | GAL | 2.229047 | 8.34E-06 | 1.34771 | 0.01259 |
| 79690 | GAL3ST4 | -1.24473 | 0.000001 | -1.29236 | 0.000001 |
| 2584 | GALK1 | -2.27242 | 0.000001 | -0.82803 | 0.000001 |
| 130589 | GALM | -1.93481 | 0.000001 | -1.41935 | 0.000001 |
| 79695 | GALNT12 | 1.172011 | 0.000001 | -1.64098 | 0.000001 |
| 114805 | GALNT13 | -1.82048 | 0.000404 | 1.369388 | 0.02609 |
| 117248 | GALNT15 | -3.35549 | 0.000001 | -2.1868 | 0.000001 |
| 57452 | GALNT16 | -2.34586 | 0.000001 | 1.27334 | 0.000001 |

| | | | | | |
|---|---|---|---|---|---|
| 374378 | GALNT18 | 2.146625 | 1.11E-05 | 0.851931 | 0.001169 |
| 8693 | GALNT4 | 1.138679 | 0.000001 | -0.7424 | 0.000001 |
| 11227 | GALNT5 | -1.7619 | 0.000001 | -1.75956 | 0.000001 |
| 2592 | GALT | -1.58422 | 0.000001 | -1.13371 | 0.000001 |
| 2593 | GAMT | -1.02034 | 0.000001 | -1.51479 | 0.000001 |
| 64762 | GAREM1 | 1.74202 | 0.000001 | -1.27234 | 0.000157 |
| 84253 | GARNL3 | -1.81737 | 0.001413 | -1.33784 | 1.74E-05 |
| 2619 | GAS1 | -2.95667 | 0.000001 | -1.70823 | 0.000001 |
| 1.01E+08 | GAS1RR | -2.37709 | 0.000001 | -1.27621 | 0.002338 |
| 283431 | GAS2L3 | 1.788923 | 0.000001 | -1.27052 | 0.000001 |
| 8522 | GAS7 | 1.351221 | 0.000001 | 2.963689 | 0.000001 |
| 2627 | GATA6 | -1.76328 | 0.000001 | 1.117242 | 0.000001 |
| 2633 | GBP1 | -2.17691 | 4.83E-05 | 1.190173 | 0.000001 |
| 2634 | GBP2 | -3.30822 | 0.000001 | -1.87769 | 0.000001 |
| 115361 | GBP4 | -4.57661 | 0.000001 | -1.44858 | 0.018031 |
| 2644 | GCHFR | -1.07412 | 0.004314 | -1.3086 | 0.001507 |
| 2645 | GCK | -2.72513 | 0.000001 | -1.74551 | 0.000402 |
| 9245 | GCNT3 | 2.292356 | 0.000001 | -2.36349 | 3.78E-05 |
| 145781 | GCOM1 | 3.340618 | 0.000001 | -0.99552 | 0.012243 |
| 9615 | GDA | 5.19107 | 0.000001 | -2.62082 | 2.93E-06 |
| 2662 | GDF10 | -2.51378 | 9.23E-06 | 2.208141 | 0.000001 |
| 8200 | GDF5 | -3.18427 | 0.000001 | -2.73748 | 1.41E-06 |
| 2668 | GDNF | 1.558437 | 0.000001 | 1.649543 | 0.000001 |
| 81544 | GDPD5 | -2.4228 | 0.000001 | -0.95031 | 1.67E-06 |
| 54438 | GFOD1 | 1.984274 | 0.000001 | -1.19848 | 1.02E-06 |
| 84514 | GHDC | -1.08092 | 0.000001 | -0.70018 | 0.000001 |
| 2690 | GHR | -1.42006 | 0.000001 | -1.26975 | 0.000001 |
| 26157 | GIMAP2 | -4.10236 | 0.000001 | -2.0049 | 0.0006 |
| 9837 | GINS1 | 2.433048 | 0.000001 | -1.17062 | 0.000001 |

| | | | | | |
|---|---|---|---|---|---|
| 51659 | GINS2 | 1.75679 | 0.000001 | -1.30476 | 0.000001 |
| 64785 | GINS3 | 2.166794 | 0.000001 | -0.71379 | 0.000704 |
| 2736 | GLI2 | -2.17745 | 0.000001 | 1.324755 | 0.000001 |
| 148979 | GLIS1 | -2.8832 | 0.000001 | -0.76676 | 0.002376 |
| 2745 | GLRX | -1.0338 | 2.78E-06 | -1.00015 | 0.000001 |
| 2760 | GM2A | -1.11047 | 0.000001 | -0.89713 | 0.000001 |
| 51053 | GMNN | 1.28203 | 0.000001 | -0.64634 | 0.000105 |
| 2766 | GMPR | -1.84015 | 0.000001 | -1.21522 | 1.63E-05 |
| 2788 | GNG7 | -2.45879 | 0.000257 | -1.2702 | 0.001988 |
| 401647 | GOLGA7B | 1.949505 | 1.4E-05 | -1.22579 | 0.019699 |
| 2820 | GPD2 | -1.10961 | 0.000001 | -0.96016 | 0.000001 |
| 2824 | GPM6B | -1.26771 | 0.000001 | -2.65773 | 0.000001 |
| 27239 | GPR162 | -1.90521 | 4.84E-06 | -1.4469 | 6.45E-06 |
| 54328 | GPR173 | -1.50388 | 6.89E-05 | 0.849897 | 4.82E-06 |
| 54329 | GPR85 | -1.88523 | 0.000001 | 0.741797 | 0.000187 |
| 9737 | GPRASP1 | -2.03747 | 0.000001 | -1.14556 | 0.000001 |
| 9052 | GPRC5A | 3.276724 | 0.000001 | -0.70547 | 0.01826 |
| 51704 | GPRC5B | -3.7893 | 0.000001 | -1.31118 | 0.000001 |
| 2890 | GRIA1 | -1.63515 | 0.000001 | -3.0718 | 0.000001 |
| 2892 | GRIA3 | -1.12594 | 0.000001 | 0.991138 | 0.000001 |
| 116444 | GRIN3B | -1.16663 | 0.006881 | -1.58379 | 0.00012 |
| 2869 | GRK5 | -1.37986 | 0.000001 | -1.20264 | 0.000001 |
| 79792 | GSDMD | -2.99714 | 0.000001 | -0.7763 | 0.000001 |
| 2944 | GSTM1 | -1.03977 | 0.005168 | -0.87029 | 0.000001 |
| 2946 | GSTM2 | -1.55288 | 0.000001 | -1.39866 | 0.000001 |
| 2948 | GSTM4 | -1.41753 | 0.000001 | -1.31018 | 0.000001 |
| 2949 | GSTM5 | -1.30768 | 0.000001 | -1.40488 | 0.000001 |
| 119391 | GSTO2 | 1.888191 | 0.000001 | 0.812443 | 0.007449 |
| 23560 | GTPBP4 | 1.750245 | 0.000001 | 0.775194 | 0.000001 |

| | | | | | |
|---|---|---|---|---|---|
| 2982 | GUCY1A1 | 3.590239 | 0.000001 | -1.64333 | 0.000346 |
| 2983 | GUCY1B1 | 1.777991 | 0.000001 | -1.91508 | 0.000001 |
| 51454 | GULP1 | 1.647112 | 0.000001 | -0.644 | 0.00105 |
| 727936 | GXYLT2 | -1.62195 | 0.000001 | 2.016455 | 0.000001 |
| 8908 | GYG2 | -3.34217 | 0.000001 | -0.85382 | 0.00915 |
| 92815 | H2AW | 1.02547 | 0.004115 | -0.97746 | 2.28E-05 |
| 9563 | H6PD | -1.50588 | 0.000001 | -0.78003 | 0.000001 |
| 9464 | HAND2 | -1.68324 | 0.000001 | -1.17012 | 0.000001 |
| 79804 | HAND2-AS1 | -2.93671 | 0.000001 | -0.87649 | 0.004928 |
| 768096 | HAR1A | -1.99957 | 0.006105 | -1.44631 | 0.017658 |
| 3037 | HAS2 | 2.318289 | 0.000001 | 0.668576 | 0.020472 |
| 83903 | HASPIN | 2.242278 | 0.000001 | -0.89401 | 0.001033 |
| 54930 | HAUS4 | -2.14415 | 0.000001 | -0.65422 | 0.000001 |
| 1839 | HBEGF | 2.727731 | 0.000001 | 5.957083 | 0.000001 |
| 3055 | HCK | -2.73083 | 1.13E-06 | 2.27928 | 3.95E-05 |
| 79885 | HDAC11 | -2.03671 | 0.000001 | -0.63333 | 3E-05 |
| 10014 | HDAC5 | -1.40467 | 0.000001 | -0.66935 | 0.000001 |
| 9734 | HDAC9 | -1.0308 | 2.88E-05 | -0.91523 | 0.000001 |
| 23072 | HECW1 | -1.08132 | 0.000001 | 2.609844 | 0.000001 |
| 3070 | HELLS | 1.663808 | 0.000001 | -1.2207 | 0.000001 |
| 85441 | HELZ2 | -2.39254 | 0.000001 | -1.22484 | 0.000001 |
| 51191 | HERC5 | -2.86519 | 8.88E-05 | -1.6866 | 0.000001 |
| 55008 | HERC6 | -3.70976 | 0.000001 | -1.65026 | 0.000001 |
| 9709 | HERPUD1 | -1.40467 | 0.000001 | 1.235666 | 0.000001 |
| 23462 | HEY1 | 1.60313 | 0.001401 | 1.192891 | 0.00199 |
| 3077 | HFE | -1.35172 | 0.000001 | -1.23777 | 0.000001 |
| 55733 | HHAT | -1.71639 | 0.000001 | 0.727724 | 0.000001 |
| 11112 | HIBADH | -2.73976 | 0.000001 | -0.69437 | 0.000001 |
| 9026 | HIP1R | -1.13743 | 0.000001 | -0.90024 | 1.59E-05 |

| | | | | | |
|---|---|---|---|---|---|
| 3097 | HIVEP2 | 1.393308 | 0.000001 | 0.906955 | 0.000001 |
| 3108 | HLA-DMA | -1.206 | 0.000248 | -0.86808 | 0.021318 |
| 3115 | HLA-DPB1 | -1.32203 | 0.000001 | -0.85531 | 4.18E-06 |
| 3142 | HLX | 1.25517 | 0.000001 | 1.761994 | 0.000001 |
| 83872 | HMCN1 | 1.018062 | 0.000001 | 1.634028 | 1.58E-06 |
| 1E+08 | HMGA2-AS1 | -2.60847 | 0.000001 | -0.95231 | 0.000844 |
| 3149 | HMGB3 | -1.01974 | 0.000001 | 0.862362 | 0.000001 |
| 3157 | HMGCS1 | -1.20711 | 0.000001 | -0.60894 | 5.7E-06 |
| 9324 | HMGN3 | -1.80295 | 0.000001 | -0.82759 | 3.38E-05 |
| 22993 | HMGXB3 | 1.356458 | 0.000001 | 0.697067 | 0.000001 |
| 3161 | HMMR | -1.44458 | 6.23E-05 | -1.15754 | 0.000001 |
| 112817 | HOGA1 | -3.24781 | 0.000001 | -1.34492 | 0.028043 |
| 9456 | HOMER1 | 1.617071 | 0.000001 | 1.475758 | 0.000001 |
| 3209 | HOXA13 | -1.44557 | 7.44E-06 | -2.16231 | 0.000001 |
| 3205 | HOXA9 | -1.16146 | 0.000001 | -0.88736 | 3.28E-06 |
| 3212 | HOXB2 | 1.932295 | 0.000001 | 1.43702 | 0.000001 |
| 3225 | HOXC9 | -1.19338 | 0.000001 | 0.690914 | 0.000134 |
| 3236 | HOXD10 | -1.40041 | 0.000001 | 0.739342 | 0.015746 |
| 3234 | HOXD8 | -1.75244 | 0.000001 | 0.794546 | 0.000699 |
| 3235 | HOXD9 | -2.58682 | 0.000001 | 0.705217 | 0.031428 |
| 3241 | HPCAL1 | 1.07273 | 0.000001 | -1.01057 | 0.000001 |
| 3269 | HRH1 | 2.40062 | 0.000001 | -1.18329 | 0.000001 |
| 7923 | HSD17B8 | -2.64809 | 0.000001 | -0.95598 | 0.009758 |
| 3299 | HSF4 | -1.32815 | 6.18E-05 | -1.15943 | 0.010861 |
| 3313 | HSPA9 | 1.810346 | 0.000001 | 0.795955 | 0.000001 |
| 3316 | HSPB2 | -1.0227 | 0.000001 | -1.03243 | 0.000001 |
| 8988 | HSPB3 | -2.24749 | 0.000001 | -3.40687 | 0.000001 |
| 3339 | HSPG2 | -1.35315 | 0.000001 | 1.066811 | 0.000001 |
| 3351 | HTR1B | 2.163749 | 0.001978 | -0.77736 | 0.042311 |

| | | | | | |
|---|---|---|---|---|---|
| 94031 | HTRA3 | -1.00892 | 8.62E-06 | -2.1269 | 0.000001 |
| 84329 | HVCN1 | -1.87489 | 0.000202 | -0.88786 | 0.007866 |
| 3373 | HYAL1 | -2.24 | 0.000001 | -1.0087 | 0.047904 |
| 130026 | ICA1L | -1.37205 | 0.000001 | -0.61078 | 0.004373 |
| 3399 | ID3 | -1.87886 | 0.000001 | 3.067628 | 0.000001 |
| 3417 | IDH1 | -1.43755 | 0.000001 | -0.94562 | 0.000001 |
| 3418 | IDH2 | -2.67853 | 0.000001 | 0.627142 | 0.000001 |
| 55853 | IDI2-AS1 | -1.05684 | 0.000108 | 0.727294 | 0.008413 |
| 8870 | IER3 | 1.258743 | 0.000001 | 1.274302 | 0.001096 |
| 25900 | IFFO1 | -1.08567 | 0.000001 | -1.34974 | 0.000001 |
| 3428 | IFI16 | -1.16952 | 0.000001 | -0.7695 | 0.000001 |
| 10561 | IFI44 | -2.53074 | 1.73E-06 | -0.9782 | 0.000369 |
| 64135 | IFIH1 | -4.04522 | 0.000001 | -1.11474 | 0.00024 |
| 3434 | IFIT1 | -4.27002 | 0.000001 | -1.54469 | 0.00215 |
| 3433 | IFIT2 | -4.57637 | 0.000001 | -1.25572 | 0.000001 |
| 3437 | IFIT3 | -3.38251 | 0.000001 | -1.52131 | 0.000001 |
| 8519 | IFITM1 | -1.48984 | 0.000001 | -1.33267 | 0.000001 |
| 402778 | IFITM10 | 4.182947 | 0.000001 | 1.679223 | 0.000001 |
| 10581 | IFITM2 | -1.15636 | 0.000001 | -0.76933 | 2.39E-06 |
| 10410 | IFITM3 | -1.0138 | 1.3E-06 | -0.67915 | 5.58E-06 |
| 338376 | IFNE | 2.248851 | 1.8E-05 | 1.855263 | 0.000561 |
| 3475 | IFRD1 | 2.137024 | 0.000001 | 1.170437 | 0.000001 |
| 8100 | IFT88 | -1.00723 | 3.06E-06 | -0.64605 | 2.21E-06 |
| 57722 | IGDCC4 | -1.14345 | 0.000001 | -0.68112 | 0.000001 |
| 3486 | IGFBP3 | 1.474959 | 0.004401 | 3.739128 | 0.000001 |
| 3487 | IGFBP4 | -1.77396 | 0.000001 | -0.86575 | 0.000001 |
| 3488 | IGFBP5 | -1.11399 | 0.000121 | -0.67854 | 0.00035 |
| 79713 | IGFLR1 | -1.5716 | 0.003129 | -0.92847 | 0.038372 |
| 285313 | IGSF10 | -2.26391 | 0.000001 | -1.94283 | 0.000001 |

| | | | | | |
|---|---|---|---|---|---|
| 3589 | IL11 | 2.556305 | 0.000001 | 5.221622 | 0.000001 |
| 3592 | IL12A | 2.941435 | 0.000001 | 0.761309 | 0.012716 |
| 3601 | IL15RA | -1.70255 | 0.000001 | -1.55469 | 0.000001 |
| 8809 | IL18R1 | 2.78881 | 0.000001 | -1.36481 | 0.004595 |
| 9173 | IL1RL1 | 4.776909 | 0.000001 | -1.34814 | 0.026177 |
| 3557 | IL1RN | 3.290679 | 0.000001 | 1.252599 | 0.000253 |
| 53832 | IL20RA | -2.28026 | 1.64E-05 | -1.64077 | 4.95E-06 |
| 50615 | IL21R | -3.98527 | 0.000001 | 1.022074 | 4.72E-05 |
| 9235 | IL32 | -1.10495 | 0.000001 | 1.219666 | 5.41E-06 |
| 146433 | IL34 | -3.37921 | 0.000001 | -1.14785 | 0.01236 |
| 259307 | IL4I1 | -1.55953 | 0.005125 | -1.37155 | 0.006832 |
| 3569 | IL6 | -2.21392 | 4.34E-05 | 1.837103 | 0.000107 |
| 3570 | IL6R | 1.792853 | 0.000001 | -1.42193 | 0.000001 |
| 3575 | IL7R | 1.651852 | 0.004735 | 1.668263 | 0.000001 |
| 3625 | INHBB | -4.53795 | 0.000001 | -1.47415 | 0.002361 |
| 11185 | INMT | -3.06641 | 0.000001 | -0.9678 | 0.005224 |
| 642938 | INSYN2A | 3.149511 | 0.000001 | 3.396014 | 0.000001 |
| 1.01E+08 | IQCJ-SCHIP1 | 1.286607 | 0.000001 | 1.738687 | 0.000001 |
| 9922 | IQSEC1 | -1.2503 | 0.000001 | 0.644174 | 0.000001 |
| 134728 | IRAK1BP1 | -2.7448 | 0.000001 | -0.8992 | 0.000001 |
| 3660 | IRF2 | -2.2084 | 0.000001 | -1.05768 | 0.000001 |
| 3665 | IRF7 | -3.31861 | 0.000001 | -0.63444 | 0.005815 |
| 8660 | IRS2 | -1.23076 | 0.000001 | 0.733046 | 0.000001 |
| 79191 | IRX3 | -1.07293 | 0.000001 | 0.973162 | 0.000001 |
| 79190 | IRX6 | -1.02513 | 0.00229 | -2.30669 | 0.000001 |
| 3669 | ISG20 | -2.61691 | 0.000001 | -1.75842 | 0.000001 |
| 140862 | ISM1 | 1.309953 | 0.000001 | -1.49841 | 0.000372 |
| 79763 | ISOC2 | -1.28564 | 0.000001 | 0.671893 | 0.000001 |
| 3655 | ITGA6 | 1.604849 | 0.000001 | -1.28781 | 0.000001 |

| | | | | | |
|---|---|---|---|---|---|
| 3679 | ITGA7 | -1.45184 | 0.000001 | 1.218751 | 0.000001 |
| 1.02E+08 | ITGA9-AS1 | -1.90197 | 0.000001 | -1.44356 | 0.000001 |
| 1.02E+08 | ITGB8-AS1 | -1.3534 | 0.001999 | -1.6476 | 0.002391 |
| 80760 | ITIH5 | -1.22101 | 0.000001 | -1.20933 | 0.000001 |
| 3706 | ITPKA | 3.455612 | 0.000001 | -1.16275 | 0.005016 |
| 3707 | ITPKB | -1.47652 | 0.000001 | -2.04925 | 0.000001 |
| 85450 | ITPRIP | 2.291878 | 0.000001 | 1.568403 | 0.000001 |
| 10625 | IVNS1ABP | 1.230976 | 0.000001 | 2.471756 | 0.000001 |
| 23338 | JADE2 | -1.42517 | 0.000001 | -1.72935 | 0.000001 |
| 3718 | JAK3 | -2.05617 | 0.000227 | 1.33077 | 0.000472 |
| 23210 | JMJD6 | 1.84518 | 0.000001 | 0.947479 | 0.000001 |
| 8850 | KAT2B | -1.54052 | 0.000001 | -1.50417 | 0.000001 |
| 23522 | KAT6B | -2.48661 | 0.000001 | -0.65961 | 0.000001 |
| 143879 | KBTBD3 | -1.96115 | 0.000001 | -0.77209 | 0.00777 |
| 84078 | KBTBD7 | -1.7533 | 0.000001 | -0.87266 | 2.65E-06 |
| 3738 | KCNA3 | 1.949839 | 0.000001 | -1.21755 | 0.000869 |
| 8514 | KCNAB2 | -1.8122 | 0.000001 | -1.03216 | 0.000001 |
| 3745 | KCNB1 | -3.52888 | 0.000001 | -2.74964 | 0.000001 |
| 3750 | KCND1 | -1.53164 | 0.000101 | 0.951912 | 0.001137 |
| 3751 | KCND2 | -3.50272 | 0.000001 | -1.15688 | 0.014387 |
| 3752 | KCND3 | -1.79279 | 0.000001 | 1.054076 | 0.000001 |
| 3755 | KCNG1 | 2.148676 | 0.000001 | 2.127971 | 0.000001 |
| 3756 | KCNH1 | -2.19587 | 7.06E-06 | 4.262798 | 0.000001 |
| 3759 | KCNJ2 | -1.72779 | 0.000001 | -0.85263 | 0.000604 |
| 3763 | KCNJ6 | -1.74542 | 0.000001 | 1.225441 | 0.006527 |
| 60598 | KCNK15 | -1.19363 | 0.000001 | 1.094827 | 0.000001 |
| 3776 | KCNK2 | -1.98709 | 0.000001 | -1.56326 | 0.000001 |
| 51305 | KCNK9 | -2.90754 | 0.000266 | 3.680465 | 0.000001 |
| 3778 | KCNMA1 | 1.489881 | 0.000001 | 0.802512 | 0.000001 |

| | | | | | |
|---|---|---|---|---|---|
| 3783 | KCNN4 | 1.249542 | 0.004529 | 3.532209 | 0.000001 |
| 56479 | KCNQ5 | 2.629519 | 0.000001 | -0.79036 | 7.1E-05 |
| 3788 | KCNS2 | -2.04962 | 0.000001 | -1.96697 | 0.000001 |
| 11015 | KDELR3 | -1.92335 | 0.000001 | 0.622396 | 0.000001 |
| 57710 | KIAA1614 | -1.97841 | 0.000001 | -0.90682 | 0.000001 |
| 85449 | KIAA1755 | -3.33469 | 0.000001 | 2.559001 | 0.000001 |
| 23303 | KIF13B | -1.97458 | 0.000001 | -1.29633 | 0.000001 |
| 57576 | KIF17 | 1.635072 | 0.000001 | -0.61221 | 0.014228 |
| 81930 | KIF18A | 2.996753 | 0.000001 | -0.91836 | 8.14E-05 |
| 146909 | KIF18B | 1.734579 | 0.000001 | -1.37929 | 0.000001 |
| 55605 | KIF21A | 1.699695 | 0.000001 | 2.293599 | 0.000001 |
| 145694 | KIF23-AS1 | -1.14959 | 0.000742 | 0.661391 | 0.003037 |
| 347240 | KIF24 | 1.273885 | 0.000001 | -1.003 | 6.28E-06 |
| 26153 | KIF26A | 2.718362 | 0.000001 | -1.88831 | 0.000001 |
| 55083 | KIF26B | -2.48011 | 0.000001 | 2.275133 | 0.000001 |
| 24137 | KIF4A | 1.672381 | 0.000001 | -1.11672 | 0.000001 |
| 3833 | KIFC1 | 1.488153 | 0.000001 | -1.34947 | 0.000001 |
| 84623 | KIRREL3 | 1.513467 | 0.000001 | 1.635749 | 0.000001 |
| 3815 | KIT | -2.50245 | 0.000001 | -4.70682 | 0.000001 |
| 55857 | KIZ | -1.99689 | 0.000001 | -0.96062 | 0.000001 |
| 10365 | KLF2 | 1.08526 | 0.000001 | 0.756969 | 0.000001 |
| 8609 | KLF7 | -1.93737 | 0.000001 | 2.035456 | 0.000001 |
| 687 | KLF9 | -1.59617 | 0.000001 | -0.82903 | 0.000001 |
| 126823 | KLHDC9 | -2.68581 | 1.33E-06 | -1.32859 | 0.005154 |
| 11275 | KLHL2 | 1.154361 | 0.000001 | 0.680774 | 0.000001 |
| 377007 | KLHL30 | -2.27536 | 0.006519 | 1.138959 | 0.000235 |
| 3840 | KPNA4 | 1.171627 | 0.000001 | 0.629573 | 0.000001 |
| 83999 | KREMEN1 | -1.91938 | 0.000001 | -1.17625 | 0.000001 |
| 3861 | KRT14 | 5.060362 | 0.000001 | 2.311068 | 1.61E-05 |

| | | | | | |
|---|---|---|---|---|---|
| 3866 | KRT15 | 2.09349 | 0.007339 | -1.53617 | 0.011329 |
| 3868 | KRT16 | 6.081604 | 0.000001 | 1.665659 | 0.004588 |
| 3875 | KRT18 | 1.837781 | 0.000175 | 1.560005 | 0.000602 |
| 3855 | KRT7 | 1.068549 | 3.74E-05 | 4.52304 | 0.000001 |
| 3887 | KRT81 | 5.21462 | 0.000001 | 5.563944 | 0.000001 |
| 1E+08 | KTN1-AS1 | -1.35145 | 9.1E-05 | -0.73232 | 0.029892 |
| 56267 | KYAT3 | -1.20104 | 0.000001 | -0.93797 | 0.000001 |
| 54596 | L1TD1 | 2.338767 | 0.000001 | 1.673895 | 0.000001 |
| 3908 | LAMA2 | -1.20049 | 0.000001 | -0.67851 | 5.59E-05 |
| 10319 | LAMC3 | -1.77591 | 3.2E-06 | 1.770217 | 0.000001 |
| 150082 | LCA5L | -1.84884 | 8.23E-06 | 0.671668 | 0.047933 |
| 401562 | LCNL1 | -4.50939 | 0.000001 | -1.3516 | 0.000411 |
| 3936 | LCP1 | 3.918347 | 0.000001 | 1.171905 | 0.000001 |
| 9079 | LDB2 | -1.61526 | 0.000001 | -1.72334 | 0.000001 |
| 753 | LDLRAD4 | 2.156245 | 0.000001 | 3.619453 | 0.000001 |
| 51176 | LEF1 | 4.207423 | 0.000001 | 2.38188 | 0.000001 |
| 3953 | LEPR | -1.35337 | 0.000001 | -0.89134 | 0.000001 |
| 137994 | LETM2 | 1.158097 | 7.71E-05 | -1.07549 | 2.32E-05 |
| 3965 | LGALS9 | -1.96694 | 0.000001 | -0.91531 | 5.8E-06 |
| 29094 | LGALSL | 2.814951 | 0.000001 | -0.93512 | 0.00016 |
| 8549 | LGR5 | -1.93499 | 0.000001 | -1.28791 | 0.001124 |
| 10186 | LHFPL6 | -1.174 | 0.000001 | -0.81047 | 0.000001 |
| 644714 | LIMD1-AS1 | 1.169992 | 0.000001 | -1.23917 | 0.000001 |
| 80774 | LIMD2 | -1.75174 | 0.000001 | 0.799114 | 4.38E-06 |
| 64130 | LIN7B | -2.62189 | 0.000001 | 0.797323 | 0.004928 |
| 400619 | LINC00511 | 1.760269 | 0.000001 | 0.663355 | 0.046574 |
| 400121 | LINC00547 | 2.064241 | 0.002286 | 1.265507 | 0.043381 |
| 1.01E+08 | LINC00565 | -1.93363 | 0.000001 | 0.754592 | 0.00157 |
| 283624 | LINC00641 | 1.935437 | 0.000001 | 0.864745 | 0.000144 |

| | | | | | |
|---|---|---|---|---|---|
| 400660 | LINC00683 | -2.22718 | 4.29E-05 | -2.57326 | 0.000001 |
| 643529 | LINC00865 | -1.68251 | 0.001977 | 0.723337 | 0.000984 |
| 255031 | LINC00957 | -2.18184 | 0.000001 | -0.62703 | 0.035997 |
| 1.01E+08 | LINC01013 | 2.793987 | 1.05E-05 | 4.296355 | 0.000001 |
| 152742 | LINC01085 | 2.84914 | 0.000345 | -1.19141 | 0.041148 |
| 375295 | LINC01116 | -1.43502 | 0.000001 | 0.804689 | 0.020795 |
| 728431 | LINC01137 | -1.6748 | 0.000001 | 1.116005 | 0.000001 |
| 1.02E+08 | LINC01358 | 1.696522 | 1.23E-05 | -1.28756 | 0.014269 |
| 1E+08 | LINC01415 | -1.66501 | 0.001272 | 1.439678 | 0.00021 |
| 1.01E+08 | LINC01503 | -1.83175 | 0.000001 | 1.992592 | 0.000001 |
| 1.08E+08 | LINC02019 | 2.013035 | 0.000001 | -1.19185 | 0.041023 |
| 1.1E+08 | LINC02154 | 3.789346 | 0.000001 | -1.44742 | 0.01327 |
| 1.02E+08 | LINC02202 | -3.00881 | 0.000001 | -1.50887 | 0.000124 |
| 1E+08 | LINC02593 | -1.90676 | 0.000872 | 3.664144 | 0.000001 |
| 1.05E+08 | LINC02609 | -1.96373 | 3.72E-06 | -1.13502 | 0.009765 |
| 1.05E+08 | LINC02701 | -3.8631 | 0.000001 | -1.57053 | 0.000103 |
| 1.03E+08 | LINC02861 | -3.00358 | 0.000001 | -0.96655 | 0.008215 |
| 388407 | LINC02875 | -1.56888 | 0.003701 | -1.2112 | 0.003526 |
| 3988 | LIPA | -1.63856 | 0.000001 | -1.45407 | 0.000001 |
| 3990 | LIPC | -1.75691 | 0.000001 | -1.94284 | 1.16E-05 |
| 9388 | LIPG | 2.012183 | 0.000001 | 2.725597 | 0.000001 |
| 4005 | LMO2 | -1.59819 | 0.000132 | -3.89825 | 0.000001 |
| 25802 | LMOD1 | -2.98224 | 0.000001 | 1.282925 | 0.000001 |
| 1.05E+08 | LNCOG | 3.702988 | 0.000001 | 1.053265 | 0.006126 |
| 1.03E+08 | LNCTAM34A | -1.51123 | 4.26E-06 | -1.39367 | 3.4E-06 |
| 9361 | LONP1 | 1.437819 | 0.000001 | 0.970991 | 0.000001 |
| 91694 | LONRF1 | -1.1328 | 0.000001 | -0.77715 | 3.6E-06 |
| 79836 | LONRF3 | 3.744455 | 0.000001 | 1.024504 | 0.022634 |
| 4016 | LOXL1 | -2.06287 | 0.000001 | 0.657767 | 0.000001 |

| | | | | | |
|---|---|---|---|---|---|
| 84695 | LOXL3 | -1.98072 | 0.000001 | 1.059313 | 0.000001 |
| 84171 | LOXL4 | -3.10376 | 0.000001 | -1.03056 | 0.019478 |
| 23566 | LPAR3 | 3.432711 | 0.000001 | -1.76387 | 0.000001 |
| 2846 | LPAR4 | -1.91976 | 5.47E-05 | -1.671 | 3.5E-05 |
| 10161 | LPAR6 | -1.40174 | 2.37E-06 | -1.54712 | 0.000001 |
| 64900 | LPIN3 | -1.24125 | 0.000001 | -1.1488 | 0.000001 |
| 121227 | LRIG3 | -2.04666 | 0.000001 | -1.06241 | 0.000001 |
| 83938 | LRMDA | -1.49367 | 2.39E-06 | -0.74713 | 0.026545 |
| 29967 | LRP12 | 1.129184 | 0.000001 | 0.602287 | 0.000001 |
| 55805 | LRP2BP | 1.686577 | 2.53E-05 | 0.971868 | 0.029067 |
| 122769 | LRR1 | 1.255511 | 0.000001 | -0.69539 | 0.00014 |
| 10234 | LRRC17 | -1.90819 | 9.88E-05 | 1.82027 | 0.000001 |
| 55222 | LRRC20 | -2.3962 | 0.000001 | -1.65942 | 0.000001 |
| 81543 | LRRC3 | -2.07811 | 0.000001 | -0.92078 | 0.000001 |
| 55073 | LRRC37A4P | 1.681945 | 0.000001 | -0.69747 | 0.008079 |
| 201255 | LRRC45 | -1.66237 | 0.000001 | -1.62886 | 0.000001 |
| 54839 | LRRC49 | -1.03274 | 3.27E-06 | -0.72985 | 0.000274 |
| 55379 | LRRC59 | 1.225488 | 0.000001 | 0.7178 | 0.000001 |
| 221424 | LRRC73 | -1.50344 | 2.73E-05 | -1.11445 | 0.013067 |
| 84230 | LRRC8C | 1.480431 | 0.000001 | -0.70531 | 0.000376 |
| 120892 | LRRK2 | -2.19482 | 0.000001 | -1.3297 | 0.000001 |
| 221091 | LRRN4CL | -1.52462 | 0.000001 | -3.17997 | 0.000001 |
| 4047 | LSS | -2.06602 | 0.000001 | -1.98124 | 0.000001 |
| 4052 | LTBP1 | -1.37527 | 0.000001 | 1.653176 | 0.000001 |
| 8425 | LTBP4 | -1.57065 | 0.000001 | -0.62733 | 0.000001 |
| 541468 | LURAP1 | -1.27475 | 0.000001 | -1.46845 | 0.000001 |
| 206338 | LVRN | -1.32897 | 7.94E-06 | -1.38198 | 0.005285 |
| 1E+08 | LY6E-DT | -1.48949 | 8.21E-06 | -2.78475 | 0.000001 |
| 4065 | LY75 | -2.6454 | 0.000001 | -1.62166 | 0.001468 |

| | | | | | |
|---|---|---|---|---|---|
| 1.01E+08 | LY75-CD302 | -1.36264 | 0.001002 | -1.48432 | 0.014929 |
| 23643 | LY96 | -1.83836 | 0.000001 | -0.67843 | 0.000301 |
| 66004 | LYNX1 | -1.47028 | 0.000001 | -1.73801 | 0.000001 |
| 1.11E+08 | LYNX1-SLURP2 | -1.18199 | 8.74E-05 | -1.67159 | 5.8E-06 |
| 27076 | LYPD3 | 1.351569 | 0.002659 | -1.82898 | 0.000427 |
| 130574 | LYPD6 | -1.80332 | 0.000001 | -1.5709 | 0.000001 |
| 130576 | LYPD6B | -1.76186 | 0.0012 | -1.26831 | 0.040397 |
| 28992 | MACROD1 | -1.02401 | 0.000001 | -0.60377 | 0.000145 |
| 140733 | MACROD2 | -1.79656 | 0.000001 | -0.86575 | 0.003595 |
| 55506 | MACROH2A2 | -1.81938 | 0.000001 | -0.61455 | 2.92E-06 |
| 4085 | MAD2L1 | 1.150985 | 0.000001 | -1.10651 | 0.000001 |
| 9935 | MAFB | -2.7368 | 0.000001 | -1.5846 | 0.000001 |
| 4097 | MAFG | 2.055268 | 0.000001 | 0.803618 | 0.000001 |
| 9500 | MAGED1 | -1.07729 | 0.000001 | 0.752358 | 0.000001 |
| 728239 | MAGED4 | -1.73367 | 0.000001 | 0.652404 | 0.000001 |
| 81557 | MAGED4B | -1.38786 | 0.000001 | 0.689649 | 0.000001 |
| 57692 | MAGEE1 | -2.13548 | 0.000001 | -0.72312 | 0.008592 |
| 84549 | MAK16 | 1.859744 | 0.000001 | 0.737392 | 0.000001 |
| 7851 | MALL | 1.298542 | 0.00054 | -1.04829 | 2.16E-05 |
| 256691 | MAMDC2 | 3.254471 | 0.000001 | 0.818722 | 0.002829 |
| 84441 | MAML2 | -1.68456 | 0.000001 | 0.765186 | 0.000001 |
| 284358 | MAMSTR | -2.21356 | 0.000161 | -1.89208 | 2.6E-05 |
| 1E+08 | MAN1B1-DT | -1.05787 | 0.000499 | 0.60682 | 0.014982 |
| 57134 | MAN1C1 | -3.24688 | 0.000001 | -1.62636 | 0.000001 |
| 149175 | MANEAL | -1.15085 | 0.007171 | -1.30355 | 0.000266 |
| 54682 | MANSC1 | -1.82396 | 0.000001 | -2.03105 | 0.000001 |
| 4128 | MAOA | -2.02437 | 0.000001 | -1.88295 | 0.000001 |
| 5608 | MAP2K6 | -3.80763 | 0.000001 | -2.08331 | 0.000001 |
| 1.01E+08 | MAP3K2-DT | -1.50983 | 0.00014 | 1.069894 | 3.15E-06 |

| | | | | | |
|---|---|---|---|---|---|
| 1.18E+08 | MAP3K4-AS1 | -1.11017 | 0.000001 | 3.292109 | 0.000001 |
| 9064 | MAP3K6 | -1.57491 | 0.000001 | -0.60382 | 0.000001 |
| 56911 | MAP3K7CL | -3.33407 | 0.000001 | 2.273077 | 0.000001 |
| 5871 | MAP4K2 | -2.49499 | 0.000001 | -0.97475 | 0.000001 |
| 4135 | MAP6 | -1.5243 | 0.000001 | -0.6075 | 0.00084 |
| 4137 | MAPT | -2.20662 | 0.000001 | -2.55709 | 0.000001 |
| 57574 | MARCHF4 | 1.291864 | 0.000001 | 0.71282 | 1.8E-05 |
| 83742 | MARVELD1 | -1.78137 | 0.000001 | -0.60204 | 0.000001 |
| 5648 | MASP1 | -1.17924 | 0.000001 | -2.36503 | 0.000001 |
| 23139 | MAST2 | 1.083034 | 0.000001 | 1.04334 | 0.000001 |
| 23031 | MAST3 | -1.46072 | 0.000001 | 1.327309 | 0.000001 |
| 9782 | MATR3 | 1.080273 | 6.56E-06 | -0.95854 | 1.44E-06 |
| 55796 | MBNL3 | -1.20777 | 0.000001 | -2.03051 | 0.000001 |
| 4162 | MCAM | -1.38483 | 0.000119 | 1.712546 | 0.000001 |
| 23263 | MCF2L | 2.193155 | 6.06E-06 | -1.83079 | 0.000001 |
| 55388 | MCM10 | 3.060687 | 0.000001 | -0.94235 | 0.000001 |
| 4171 | MCM2 | 1.691637 | 0.000001 | -1.03255 | 0.000001 |
| 4172 | MCM3 | 1.476685 | 0.000001 | -0.91333 | 0.000001 |
| 4173 | MCM4 | 1.263143 | 0.000001 | -1.01788 | 0.000001 |
| 4175 | MCM6 | -1.29014 | 0.000001 | -0.6574 | 0.000001 |
| 84515 | MCM8 | 2.021027 | 0.000001 | -0.92411 | 0.000001 |
| 55283 | MCOLN3 | 1.038872 | 0.00044 | -0.64764 | 0.002872 |
| 79772 | MCTP1 | 2.316261 | 0.000001 | -1.78271 | 0.000001 |
| 55784 | MCTP2 | -1.84482 | 3.12E-06 | -0.87679 | 0.007007 |
| 2122 | MECOM | 2.904531 | 0.000001 | 1.993322 | 1.93E-06 |
| 4212 | MEIS2 | -2.06747 | 0.000001 | -0.60797 | 0.000001 |
| 56917 | MEIS3 | -1.51298 | 0.000001 | 0.980944 | 0.000001 |
| 9833 | MELK | 1.006767 | 2.95E-06 | -1.00958 | 0.000001 |
| 4222 | MEOX1 | -3.50583 | 0.000001 | 0.885284 | 0.003301 |

| | | | | | |
|---|---|---|---|---|---|
| 4223 | MEOX2 | -1.5177 | 0.00014 | -1.35238 | 0.000001 |
| 145873 | MESP2 | -1.97886 | 0.006185 | -1.10757 | 0.040594 |
| 4233 | MET | 2.240809 | 0.000001 | 0.90741 | 0.000001 |
| 728464 | METTL24 | -2.23936 | 0.000001 | -1.73651 | 0.000001 |
| 25840 | METTL7A | -2.51383 | 0.000001 | -2.49435 | 0.000001 |
| 84206 | MEX3B | -1.64483 | 0.000001 | 1.206327 | 0.000001 |
| 4237 | MFAP2 | -1.19117 | 0.000001 | 0.602057 | 0.000001 |
| 9848 | MFAP3L | 1.257549 | 0.000001 | 3.146492 | 0.000001 |
| 84709 | MGARP | -2.55689 | 0.000001 | -1.72358 | 0.000001 |
| 9645 | MICAL2 | 2.254099 | 0.000001 | 3.573604 | 0.000001 |
| 85377 | MICALL1 | 1.922269 | 0.000001 | 1.262675 | 0.000001 |
| 50488 | MINK1 | -1.26001 | 0.000001 | -0.66345 | 0.000001 |
| 399726 | MIR1915HG | -2.25128 | 0.000001 | -1.55744 | 0.000001 |
| 1E+08 | MIR193BHG | -1.21512 | 6.45E-06 | 1.429287 | 0.000001 |
| 84981 | MIR22HG | 1.023362 | 0.000001 | 1.794363 | 0.000001 |
| 284454 | MIR23AHG | 3.267426 | 0.000001 | 2.485744 | 0.000001 |
| 554202 | MIR31HG | 3.440541 | 0.000001 | 2.604241 | 0.000001 |
| 84848 | MIR503HG | -1.42757 | 0.00019 | 3.019142 | 0.000001 |
| 1.13E+08 | MIRLET7A1HG | 1.133355 | 0.000217 | 1.330357 | 5.17E-06 |
| 4286 | MITF | -2.40698 | 0.000001 | -1.77717 | 0.000001 |
| 10962 | MLLT11 | 1.290521 | 0.000001 | 2.407648 | 0.000001 |
| 4312 | MMP1 | 1.297377 | 0.000001 | -1.68978 | 0.000001 |
| 4319 | MMP10 | 1.280486 | 0.000001 | 2.924807 | 0.000001 |
| 4325 | MMP16 | 1.566486 | 0.000001 | 0.670246 | 1.32E-05 |
| 79817 | MOB3B | -2.11801 | 0.001372 | -2.20599 | 3.15E-05 |
| 283385 | MORN3 | -1.65211 | 0.00198 | -1.20351 | 0.009282 |
| 4343 | MOV10 | -2.39551 | 0.000001 | -1.07125 | 0.000001 |
| 54456 | MOV10L1 | -2.36462 | 4.15E-06 | 1.532798 | 0.000001 |
| 4350 | MPG | -1.59053 | 0.000001 | -0.94875 | 0.000001 |

| | | | | | |
|---|---|---|---|---|---|
| 10200 | MPHOSPH6 | 1.63032 | 0.000001 | -0.86782 | 0.000001 |
| 84954 | MPND | -1.98432 | 0.000001 | -0.63692 | 0.000243 |
| 55686 | MREG | 1.117986 | 0.000001 | -1.53847 | 0.000001 |
| 116534 | MRGPRE | 2.269031 | 0.000001 | -3.10659 | 0.000001 |
| 1E+08 | MSC-AS1 | -1.28677 | 1.72E-06 | 1.450597 | 0.000001 |
| 54996 | MTARC2 | -1.05954 | 8.12E-06 | -0.61504 | 0.000868 |
| 113115 | MTFR2 | 2.282801 | 0.000001 | -0.77807 | 0.021574 |
| 8898 | MTMR2 | 1.181883 | 0.000001 | 1.083167 | 0.000001 |
| 66036 | MTMR9 | 1.856066 | 0.000001 | 0.628793 | 0.000001 |
| 57509 | MTUS1 | 1.286411 | 0.000001 | -2.2702 | 0.000001 |
| 4597 | MVD | -2.18562 | 0.000001 | -0.66071 | 3.22E-06 |
| 4599 | MX1 | -2.21669 | 0.000924 | -1.50236 | 0.001035 |
| 4600 | MX2 | -3.81878 | 0.000001 | -1.43355 | 0.000001 |
| 26292 | MYCBP | -1.72084 | 0.000001 | 0.793923 | 1.09E-05 |
| 80177 | MYCT1 | 3.359748 | 9.59E-06 | 1.554423 | 0.010241 |
| 4629 | MYH11 | 1.468656 | 0.001249 | 0.946051 | 0.009122 |
| 10398 | MYL9 | -1.13726 | 0.000001 | 1.151846 | 0.000001 |
| 29116 | MYLIP | -2.03908 | 0.000001 | -2.7494 | 0.000001 |
| 8736 | MYOM1 | -2.21279 | 0.000323 | 0.790803 | 0.019123 |
| 58529 | MYOZ1 | -2.32823 | 0.009205 | 5.32999 | 0.000001 |
| 25924 | MYRIP | -1.20122 | 0.006594 | -3.07959 | 0.000001 |
| 1.01E+08 | MYZAP | 2.81164 | 0.000001 | -3.75482 | 0.000001 |
| 23148 | NACAD | -1.75257 | 0.000001 | -0.81046 | 0.000001 |
| 4668 | NAGA | -1.40267 | 0.000001 | -0.92039 | 0.000001 |
| 162417 | NAGS | 1.554865 | 1.26E-05 | -1.20871 | 0.000545 |
| 93100 | NAPRT | -1.15342 | 0.000001 | -0.87564 | 0.000001 |
| 89795 | NAV3 | 1.759484 | 0.000001 | 1.819529 | 0.000001 |
| 23218 | NBEAL2 | -2.12607 | 0.000001 | -0.90351 | 0.000001 |
| 84224 | NBPF3 | -1.8604 | 0.000001 | -1.11128 | 0.000001 |

| | | | | | |
|---|---|---|---|---|---|
| 83988 | NCALD | -1.66968 | 0.000001 | -2.15953 | 0.000001 |
| 64151 | NCAPG | 1.350544 | 0.000001 | -1.4595 | 0.000001 |
| 23397 | NCAPH | 2.127395 | 0.000001 | -1.28457 | 0.000001 |
| 4688 | NCF2 | 3.597751 | 0.000001 | 4.435987 | 0.000001 |
| 57447 | NDRG2 | -1.98651 | 0.000001 | -0.82819 | 0.001068 |
| 57446 | NDRG3 | -1.88096 | 0.000001 | -0.65509 | 0.000001 |
| 4739 | NEDD9 | -3.07213 | 0.000001 | 1.518854 | 0.000001 |
| 4747 | NEFL | 3.080849 | 0.000001 | -1.43071 | 0.000474 |
| 257194 | NEGR1 | -1.79268 | 0.000001 | 1.339174 | 0.000001 |
| 55247 | NEIL3 | 1.768385 | 4.66E-06 | -1.77035 | 0.000001 |
| 140609 | NEK7 | -1.26208 | 0.000001 | 1.6562 | 0.000001 |
| 284086 | NEK8 | -2.5478 | 0.000001 | -0.70831 | 0.000918 |
| 54492 | NEURL1B | -1.90804 | 7.94E-06 | -2.56466 | 0.000001 |
| 91624 | NEXN | -1.05361 | 0.000001 | 1.363804 | 0.000001 |
| 4772 | NFATC1 | 1.142776 | 5.48E-05 | 0.627592 | 0.002325 |
| 4774 | NFIA | -2.23534 | 0.000001 | -2.16225 | 0.000001 |
| 4781 | NFIB | -2.92156 | 0.000001 | -0.79285 | 0.005752 |
| 25791 | NGEF | 3.163696 | 0.000001 | 1.117461 | 0.032748 |
| 4803 | NGF | 1.758472 | 0.000001 | 2.010369 | 0.000001 |
| 283948 | NHLRC4 | -2.82103 | 0.000125 | -1.76907 | 0.000378 |
| 1E+08 | NINJ2-AS1 | -3.02051 | 0.000001 | -0.7293 | 0.003043 |
| 1.05E+08 | NKILA | -2.32789 | 0.001316 | 3.120752 | 0.000001 |
| 22871 | NLGN1 | -2.87177 | 0.000001 | -1.48111 | 0.000001 |
| 22829 | NLGN4Y | -1.15924 | 0.000813 | -0.7635 | 0.000001 |
| 22861 | NLRP1 | -1.38933 | 0.000001 | 1.059305 | 0.000001 |
| 4830 | NME1 | 1.657917 | 0.000001 | 0.75746 | 0.000001 |
| 4832 | NME3 | -1.87165 | 0.000001 | -1.10963 | 0.000001 |
| 344887 | NMRAL2P | 1.357671 | 0.000001 | 1.34997 | 0.000001 |
| 4837 | NNMT | -3.31286 | 0.000001 | 2.260614 | 0.000001 |

| | | | | | |
|---|---|---|---|---|---|
| 25819 | NOCT | 1.375447 | 0.000001 | 1.198015 | 0.000001 |
| 8996 | NOL3 | -1.22162 | 0.000001 | -0.64734 | 0.000598 |
| 140688 | NOL4L | -1.68086 | 0.000001 | 1.332762 | 0.000001 |
| 51491 | NOP16 | 2.16657 | 0.000001 | 0.637422 | 0.000399 |
| 29997 | NOP53 | -1.27557 | 0.000001 | -0.77244 | 0.000001 |
| 4853 | NOTCH2 | 1.100556 | 0.000001 | 0.716351 | 1.89E-05 |
| 10811 | NOXA1 | -1.91072 | 0.00046 | -1.3761 | 0.001994 |
| 4867 | NPHP1 | -1.38276 | 0.000001 | -0.96169 | 0.000001 |
| 1.01E+08 | NPTN-IT1 | 2.254614 | 0.000001 | 1.337086 | 6.15E-05 |
| 4884 | NPTX1 | -1.4307 | 0.000001 | -3.99169 | 0.000001 |
| 4885 | NPTX2 | 2.121254 | 1.38E-05 | -1.33526 | 0.000001 |
| 4886 | NPY1R | 3.456179 | 2.21E-05 | 1.826843 | 0.001419 |
| 190 | NR0B1 | 1.623819 | 0.000001 | -3.52664 | 0.000001 |
| 10062 | NR1H3 | -2.20252 | 0.000001 | -1.82509 | 0.000001 |
| 7025 | NR2F1 | -1.08509 | 0.000001 | 0.819765 | 0.000001 |
| 4929 | NR4A2 | -1.71696 | 0.000001 | -1.97873 | 0.000001 |
| 8013 | NR4A3 | -4.31977 | 0.000001 | -3.73504 | 0.000001 |
| 4893 | NRAS | 1.272267 | 0.000001 | 0.752469 | 0.000001 |
| 9315 | NREP | -1.75654 | 0.000001 | 1.996977 | 0.000001 |
| 3084 | NRG1 | 1.426572 | 0.000001 | 2.827327 | 0.000001 |
| 9542 | NRG2 | -2.27767 | 0.000288 | -2.17159 | 0.000153 |
| 51299 | NRN1 | -1.12423 | 0.000001 | -1.74692 | 0.000001 |
| 8829 | NRP1 | 1.478171 | 0.000001 | -0.61079 | 0.000485 |
| 221294 | NT5DC1 | -2.71028 | 0.000001 | -1.26123 | 0.000001 |
| 4907 | NT5E | -1.28954 | 0.000001 | -1.13228 | 0.000001 |
| 56953 | NT5M | -1.84347 | 0.000001 | -1.19372 | 0.000001 |
| 4913 | NTHL1 | -1.3434 | 0.000001 | -1.41373 | 0.000001 |
| 50863 | NTM | 1.034855 | 0.000001 | 2.192432 | 0.000001 |
| 9423 | NTN1 | -1.5092 | 0.000001 | -2.24899 | 0.000001 |

| | | | | | |
|---|---|---|---|---|---|
| 84628 | NTNG2 | -2.23469 | 0.000001 | -1.53269 | 0.000001 |
| 84284 | NTPCR | -1.66093 | 0.000001 | -0.87195 | 0.000001 |
| 4923 | NTSR1 | 2.129815 | 5.56E-05 | -1.70952 | 0.002987 |
| 81788 | NUAK2 | -2.33449 | 0.000001 | 1.570581 | 0.000001 |
| 283927 | NUDT7 | -3.04 | 0.000001 | -1.5197 | 0.000001 |
| 254552 | NUDT8 | -2.06299 | 0.000001 | -0.82385 | 6.03E-05 |
| 11248 | NXPH3 | -3.27419 | 0.000001 | 0.808694 | 0.001704 |
| 222950 | NYAP1 | 1.176409 | 0.000416 | -2.32243 | 0.000001 |
| 57523 | NYNRIN | -1.73142 | 0.000001 | -0.89533 | 0.000001 |
| 4938 | OAS1 | -3.77777 | 0.000001 | -1.80116 | 5.38E-05 |
| 4939 | OAS2 | -3.6308 | 0.000001 | -1.20803 | 0.001139 |
| 4940 | OAS3 | -2.6482 | 1.12E-05 | -1.89803 | 0.000001 |
| 84033 | OBSCN | -2.06023 | 0.000001 | -1.18089 | 0.000001 |
| 55130 | ODAD2 | -2.68493 | 0.000001 | -0.63048 | 0.003135 |
| 4953 | ODC1 | 4.452795 | 0.000001 | 0.621307 | 0.000001 |
| 283298 | OLFML1 | -1.6855 | 0.000001 | -2.23924 | 0.000001 |
| 169611 | OLFML2A | -3.03253 | 0.000001 | -1.24468 | 0.000001 |
| 4958 | OMD | -1.17122 | 0.002198 | -0.81201 | 0.012348 |
| 4987 | OPRL1 | -3.3578 | 0.000001 | -1.29666 | 1.18E-05 |
| 93129 | ORAI3 | -1.37162 | 0.000001 | -0.86147 | 0.000001 |
| 4998 | ORC1 | 2.484451 | 0.000001 | -1.33623 | 0.000001 |
| 23594 | ORC6 | 3.666652 | 0.000001 | -0.85432 | 0.000001 |
| 94103 | ORMDL3 | -1.42135 | 0.000001 | 0.672048 | 0.000001 |
| 23762 | OSBP2 | 2.439407 | 0.000001 | -1.18659 | 0.001447 |
| 114884 | OSBPL10 | 1.026965 | 0.000164 | 1.23702 | 0.000001 |
| 114876 | OSBPL1A | -1.99017 | 0.000001 | -0.85536 | 0.000001 |
| 114883 | OSBPL9 | -1.25946 | 0.000001 | 0.985247 | 0.000001 |
| 116039 | OSR2 | -2.22089 | 0.000001 | -2.67277 | 0.000001 |
| 192217 | OXCT2P1 | -1.41482 | 0.000001 | -0.94928 | 0.012017 |

| | | | | | |
|---|---|---|---|---|---|
| 9127 | P2RX6 | -2.27234 | 0.000001 | -1.54938 | 3.4E-05 |
| 5028 | P2RY1 | 2.683517 | 0.000001 | -2.84429 | 0.000001 |
| 29763 | PACSIN3 | -1.5961 | 0.000001 | -0.6541 | 0.000001 |
| 5058 | PAK1 | 1.098103 | 0.000001 | -0.65794 | 0.000001 |
| 5064 | PALM | -1.88925 | 0.000001 | 1.227253 | 0.000001 |
| 196743 | PAOX | -2.2456 | 0.000001 | -0.83112 | 2.34E-05 |
| 54852 | PAQR5 | 3.369764 | 0.000001 | -0.81906 | 0.001992 |
| 85315 | PAQR8 | -1.2311 | 0.000001 | -0.80959 | 1.19E-06 |
| 25849 | PARM1 | 2.282983 | 2.74E-06 | 0.968304 | 0.005435 |
| 84875 | PARP10 | -2.39077 | 0.000001 | -1.30464 | 0.000001 |
| 54625 | PARP14 | -2.89535 | 0.000001 | -0.75969 | 0.000001 |
| 83666 | PARP9 | -2.91558 | 0.000001 | -0.79104 | 0.000001 |
| 23178 | PASK | 1.001627 | 2.08E-05 | -0.82699 | 4.85E-05 |
| 1E+08 | PAXIP1-AS2 | -1.79078 | 0.000001 | -1.0355 | 0.000001 |
| 5095 | PCCA | -2.57791 | 0.000001 | -1.05005 | 0.000001 |
| 54510 | PCDH18 | -2.28843 | 0.000001 | -1.25004 | 0.000001 |
| 57526 | PCDH19 | 2.451127 | 0.000125 | 2.705191 | 0.000001 |
| 5099 | PCDH7 | -1.15712 | 0.000001 | 0.815133 | 0.000001 |
| 5106 | PCK2 | -1.78748 | 0.000001 | 1.930074 | 0.000001 |
| 54760 | PCSK4 | -2.70213 | 0.000001 | -0.8452 | 0.009122 |
| 255738 | PCSK9 | -2.52492 | 0.000001 | -1.31178 | 0.000141 |
| 51449 | PCYOX1 | -1.19173 | 0.000001 | -1.29152 | 0.000001 |
| 5833 | PCYT2 | -1.91191 | 0.000001 | -1.27392 | 0.000001 |
| 27250 | PDCD4 | -4.26428 | 0.000001 | -0.93192 | 0.000001 |
| 1.05E+08 | PDCD6IP-DT | 1.736155 | 0.000001 | 1.587963 | 0.000001 |
| 5137 | PDE1C | -1.02557 | 2.04E-06 | 1.196004 | 0.000001 |
| 5144 | PDE4D | 1.828392 | 0.000001 | 1.000988 | 0.000001 |
| 8654 | PDE5A | -2.86028 | 0.000001 | -1.32418 | 0.000001 |
| 27115 | PDE7B | -1.7576 | 1.16E-06 | -2.41769 | 0.000001 |

| | | | | | |
|---|---|---|---|---|---|
| 80310 | PDGFD | -1.67199 | 0.000001 | -2.58354 | 0.000001 |
| 5156 | PDGFRA | -1.25995 | 0.000001 | -0.88291 | 1.05E-05 |
| 5157 | PDGFRL | -2.46459 | 0.000001 | -2.32942 | 0.000001 |
| 5165 | PDK3 | 1.036025 | 2.59E-05 | -0.76111 | 0.000354 |
| 9124 | PDLIM1 | 1.946927 | 0.000001 | 0.722845 | 0.000001 |
| 27295 | PDLIM3 | 5.795793 | 0.000001 | 2.153668 | 0.000001 |
| 8572 | PDLIM4 | 1.791453 | 0.000001 | 2.283074 | 0.000001 |
| 10611 | PDLIM5 | 2.308551 | 0.000001 | 1.617038 | 0.000001 |
| 23024 | PDZRN3 | -1.55749 | 0.000001 | -0.8937 | 0.000001 |
| 8682 | PEA15 | 1.004376 | 0.000001 | 0.815738 | 0.000001 |
| 359809 | PEG13 | -3.32209 | 0.000001 | -1.51372 | 0.000001 |
| 1.01E+08 | PELATON | -1.61405 | 1.99E-06 | -1.33545 | 0.000001 |
| 246330 | PELI3 | -1.13138 | 0.000001 | -0.81548 | 0.000001 |
| 5187 | PER1 | 1.077085 | 0.000001 | -0.88491 | 0.000001 |
| 8864 | PER2 | 3.588615 | 0.000001 | 1.040004 | 4.38E-06 |
| 5202 | PFDN2 | 1.33927 | 0.000001 | 0.673359 | 0.000001 |
| 27315 | PGAP2 | -2.06381 | 0.000001 | -0.61842 | 0.000314 |
| 80162 | PGGHG | -3.60738 | 0.000001 | -1.35648 | 0.000001 |
| 283209 | PGM2L1 | 1.856964 | 0.000001 | 1.989575 | 0.000001 |
| 10424 | PGRMC2 | -1.08898 | 0.000001 | 0.609404 | 0.000001 |
| 5260 | PHKG1 | -2.5558 | 0.000001 | -3.30883 | 0.000001 |
| 22822 | PHLDA1 | 2.935417 | 0.000001 | -1.59043 | 0.000001 |
| 23239 | PHLPP1 | -1.05061 | 3.93E-05 | -0.73982 | 1.84E-05 |
| 5264 | PHYH | -1.24392 | 0.000001 | -0.65815 | 5.21E-06 |
| 254295 | PHYHD1 | -1.9035 | 0.000001 | -0.95716 | 0.000001 |
| 196500 | PIANP | -1.68655 | 0.000505 | -1.01083 | 1.93E-05 |
| 128344 | PIFO | 2.503484 | 0.000001 | -1.7841 | 8.48E-05 |
| 284098 | PIGW | 1.137083 | 0.000001 | 0.621109 | 5.49E-06 |
| 80235 | PIGZ | -1.97491 | 0.000001 | -0.9792 | 1.55E-05 |

| | | | | | |
|---|---|---|---|---|---|
| 113791 | PIK3IP1 | -3.94388 | 0.000001 | 0.753846 | 0.000001 |
| 8503 | PIK3R3 | -1.00579 | 8.55E-05 | -2.1998 | 0.000001 |
| 415116 | PIM3 | 1.149404 | 0.000001 | -1.08847 | 0.000001 |
| 114780 | PKD1L2 | -1.80429 | 0.000001 | -2.99966 | 0.000001 |
| 91461 | PKDCC | -1.07403 | 0.000001 | -0.83472 | 7.57E-05 |
| 5318 | PKP2 | 2.630172 | 0.002648 | 2.06501 | 3.07E-05 |
| 50487 | PLA2G3 | -2.82353 | 3.22E-05 | -1.61322 | 0.000628 |
| 22925 | PLA2R1 | -1.06319 | 0.000001 | -1.15586 | 0.000001 |
| 5920 | PLAAT4 | -2.02248 | 0.000001 | -1.72202 | 1.25E-05 |
| 219348 | PLAC9 | 1.046225 | 0.000001 | -0.70793 | 3.31E-06 |
| 5333 | PLCD1 | -1.53477 | 0.000001 | -1.01918 | 0.000001 |
| 84812 | PLCD4 | -1.61362 | 0.000317 | -2.19708 | 2.65E-05 |
| 5336 | PLCG2 | 1.863469 | 0.000001 | -0.87323 | 4.35E-05 |
| 257068 | PLCXD2 | 1.642414 | 0.000571 | 1.342354 | 8.6E-06 |
| 345557 | PLCXD3 | 5.933665 | 0.000001 | 0.741985 | 0.042753 |
| 5337 | PLD1 | -3.02948 | 0.000001 | -2.08047 | 0.000001 |
| 5338 | PLD2 | -2.68646 | 0.000001 | -0.88279 | 0.000001 |
| 26499 | PLEK2 | 2.006457 | 0.000001 | 2.924048 | 0.000001 |
| 57480 | PLEKHG1 | -2.09318 | 0.000001 | -2.90746 | 0.000001 |
| 130271 | PLEKHH2 | -2.06864 | 0.000001 | -0.76926 | 0.010737 |
| 51177 | PLEKHO1 | 1.531596 | 0.000001 | 0.638319 | 2.27E-06 |
| 80301 | PLEKHO2 | -1.04279 | 0.000001 | -0.85627 | 0.000001 |
| 1263 | PLK3 | 1.437461 | 0.000001 | 1.026942 | 0.000001 |
| 10733 | PLK4 | 1.859915 | 0.000001 | -1.38173 | 0.000001 |
| 5352 | PLOD2 | 1.646038 | 0.000001 | 2.636623 | 0.000001 |
| 8612 | PLPP2 | 1.3014 | 3.42E-05 | -1.74292 | 0.000001 |
| 8613 | PLPP3 | -2.93708 | 0.000001 | -1.92018 | 0.000001 |
| 84814 | PLPP7 | -1.42425 | 0.000001 | -0.74404 | 0.002537 |
| 84898 | PLXDC2 | -1.73 | 0.000001 | 1.365797 | 0.000001 |

| | | | | | |
|---|---|---|---|---|---|
| 5362 | PLXNA2 | 3.091236 | 0.000001 | 1.235382 | 0.000001 |
| 10154 | PLXNC1 | -2.73901 | 0.000001 | -0.63697 | 0.000257 |
| 5373 | PMM2 | 1.260185 | 0.000001 | 0.625008 | 0.000001 |
| 119548 | PNLIPRP3 | 3.307316 | 0.000001 | -3.07753 | 0.000001 |
| 57469 | PNMA8B | -2.51153 | 0.000001 | -1.95431 | 0.000001 |
| 56902 | PNO1 | 2.066983 | 0.000001 | 0.69178 | 0.000001 |
| 375775 | PNPLA7 | -2.71552 | 0.000001 | -1.22633 | 0.000001 |
| 79883 | PODNL1 | -1.25754 | 0.000001 | 1.740982 | 0.000001 |
| 5427 | POLE2 | 1.803002 | 0.000001 | -1.10004 | 1.02E-05 |
| 10721 | POLQ | 2.127335 | 0.000001 | -1.09202 | 8.42E-06 |
| 10849 | POLR1G | 1.622601 | 0.000001 | 2.51534 | 0.000001 |
| 8497 | PPFIA4 | -2.11824 | 0.000001 | -1.43613 | 0.001748 |
| 285755 | PPIL6 | 1.664146 | 0.000001 | -0.63751 | 0.016217 |
| 333926 | PPM1J | -2.77335 | 2.7E-06 | -1.16612 | 0.005925 |
| 152926 | PPM1K | -2.03429 | 0.000001 | -0.8802 | 0.000001 |
| 132160 | PPM1M | -1.38419 | 0.000001 | -0.61939 | 2.74E-06 |
| 81706 | PPP1R14C | 2.278228 | 0.00044 | 0.929526 | 0.000344 |
| 79660 | PPP1R3B | -1.14094 | 0.000001 | 0.857741 | 0.000001 |
| 5507 | PPP1R3C | -1.28824 | 0.000001 | -0.6184 | 0.000152 |
| 90673 | PPP1R3E | -1.56486 | 0.000001 | -0.65288 | 7.57E-05 |
| 57718 | PPP4R4 | -2.42887 | 0.000001 | -1.82629 | 0.000001 |
| 5538 | PPT1 | 1.881519 | 0.000001 | -0.63121 | 0.000001 |
| 157285 | PRAG1 | 2.420638 | 0.000001 | 2.072696 | 0.000001 |
| 1.09E+08 | PRAL | -3.21864 | 0.000001 | 1.168499 | 0.032079 |
| 10650 | PRELID3A | 1.081728 | 3.15E-05 | 1.003802 | 0.000001 |
| 5549 | PRELP | -1.86808 | 0.000001 | -1.50002 | 0.000001 |
| 57580 | PREX1 | -2.2082 | 0.000001 | -2.02654 | 0.000001 |
| 144165 | PRICKLE1 | -1.92945 | 0.000001 | 1.578884 | 0.000001 |
| 166336 | PRICKLE2 | -1.4472 | 0.000001 | 0.834858 | 0.000001 |

| | | | | | |
|---|---|---|---|---|---|
| 145270 | PRIMA1 | -2.58815 | 0.000166 | -2.6384 | 0.000001 |
| 222171 | PRR15 | -1.98 | 0.000622 | 3.275803 | 0.000001 |
| 51334 | PRR16 | -1.12211 | 0.000001 | -1.01812 | 0.000001 |
| 80863 | PRRT1 | -2.35819 | 0.000001 | -1.62755 | 1.28E-05 |
| 112476 | PRRT2 | -2.17415 | 0.000001 | -1.74078 | 0.000001 |
| 285368 | PRRT3 | -1.54516 | 0.000001 | -0.63353 | 0.00031 |
| 11098 | PRSS23 | -1.17859 | 0.000001 | 1.02939 | 0.000001 |
| 127281 | PRXL2B | -1.10042 | 0.000001 | -1.00969 | 0.000001 |
| 29968 | PSAT1 | -1.02996 | 0.000768 | 2.3379 | 0.000001 |
| 5662 | PSD | -1.13678 | 0.002223 | -0.92597 | 0.005752 |
| 5699 | PSMB10 | -1.66417 | 0.000001 | -0.77713 | 0.000001 |
| 5696 | PSMB8 | -1.3548 | 0.000001 | -1.13274 | 0.000001 |
| 5698 | PSMB9 | -3.16459 | 0.000001 | -1.8995 | 0.000001 |
| 29893 | PSMC3IP | 3.244665 | 0.000001 | -0.87926 | 0.000001 |
| 5720 | PSME1 | -1.39687 | 0.000001 | -0.85378 | 0.000001 |
| 5721 | PSME2 | -1.18255 | 0.000001 | -0.84911 | 0.000001 |
| 5727 | PTCH1 | -2.145 | 0.000001 | -0.72293 | 0.000255 |
| 9536 | PTGES | -1.93074 | 0.000001 | -1.9971 | 0.000001 |
| 5737 | PTGFR | -1.16836 | 0.000001 | -2.90339 | 0.000001 |
| 5740 | PTGIS | -1.07144 | 0.000001 | -1.47578 | 0.00043 |
| 5743 | PTGS2 | 2.067741 | 7.71E-05 | -1.4281 | 0.000001 |
| 5745 | PTH1R | -1.96707 | 6.53E-06 | -0.80784 | 0.006527 |
| 5754 | PTK7 | -1.54944 | 0.000001 | 1.546418 | 0.000001 |
| 11156 | PTP4A3 | -2.11532 | 0.000001 | -1.113 | 7.9E-06 |
| 138639 | PTPDC1 | 1.992029 | 0.000001 | 1.05597 | 0.000001 |
| 5787 | PTPRB | 2.170009 | 8.93E-05 | 0.820762 | 8.8E-05 |
| 5791 | PTPRE | 1.190916 | 0.000001 | 2.063851 | 0.000001 |
| 5798 | PTPRN | 1.675254 | 0.000001 | 1.810599 | 0.000001 |
| 10076 | PTPRU | -1.99821 | 0.000001 | -0.80044 | 6.2E-06 |

| | | | | | |
|---|---|---|---|---|---|
| 9232 | PTTG1 | 1.294071 | 5.91E-06 | -1.23721 | 0.000001 |
| 9933 | PUM3 | 1.006896 | 0.000001 | 0.618046 | 2.09E-06 |
| 5817 | PVR | 1.883143 | 0.000001 | 1.268063 | 0.000001 |
| 54899 | PXK | -2.42743 | 0.000001 | -1.07983 | 0.000001 |
| 5827 | PXMP2 | -1.03574 | 1.02E-06 | -0.77309 | 0.000001 |
| 29108 | PYCARD | -2.10556 | 0.000001 | -2.11805 | 0.000001 |
| 5831 | PYCR1 | -1.11235 | 0.000001 | 1.614276 | 0.000001 |
| 5836 | PYGL | -2.25616 | 0.000001 | -0.89967 | 0.000001 |
| 79912 | PYROXD1 | 1.036476 | 0.000001 | 0.608743 | 0.000001 |
| 84795 | PYROXD2 | -2.27456 | 0.000001 | -2.00416 | 0.000001 |
| 23475 | QPRT | -2.0439 | 0.000001 | -1.34209 | 0.000001 |
| 1.01E+08 | RAB11B-AS1 | -3.87181 | 0.000001 | -0.69684 | 0.006901 |
| 9727 | RAB11FIP3 | -1.17316 | 0.000001 | -0.72748 | 0.000001 |
| 84440 | RAB11FIP4 | -1.81579 | 1.71E-06 | -1.67308 | 0.000259 |
| 5874 | RAB27B | 1.578214 | 0.008229 | -1.6773 | 0.000001 |
| 27314 | RAB30 | -1.02209 | 1.83E-06 | 0.714141 | 6.95E-05 |
| 23682 | RAB38 | 1.302107 | 0.000391 | -2.64538 | 0.000001 |
| 5864 | RAB3A | -1.28626 | 0.003145 | -1.5226 | 0.001471 |
| 9545 | RAB3D | -1.12834 | 1.7E-05 | -2.97731 | 0.000001 |
| 115273 | RAB42 | -2.56514 | 0.000001 | -1.5709 | 0.000001 |
| 338382 | RAB7B | -1.15577 | 0.000001 | -2.58912 | 0.000001 |
| 5876 | RABGGTB | 1.795139 | 0.000001 | 0.681489 | 0.000001 |
| 5880 | RAC2 | -1.20367 | 0.000001 | -1.87624 | 0.000001 |
| 1.01E+08 | RAD51-AS1 | -2.09448 | 0.000001 | 0.892672 | 0.007392 |
| 10635 | RAD51AP1 | 1.577618 | 1.46E-06 | -1.19426 | 0.000001 |
| 5890 | RAD51B | -1.84456 | 1.35E-06 | -0.80115 | 0.001796 |
| 8438 | RAD54L | 1.355169 | 3.84E-05 | -1.46106 | 0.000001 |
| 5883 | RAD9A | 1.020256 | 3.54E-06 | -0.78385 | 0.000915 |
| 5898 | RALA | 2.213046 | 0.000001 | 1.284954 | 0.000001 |

| | | | | | |
|---|---|---|---|---|---|
| 10267 | RAMP1 | -1.11531 | 0.000772 | 2.404744 | 0.000001 |
| 5909 | RAP1GAP | 1.251492 | 0.000933 | 1.185928 | 0.001175 |
| 10411 | RAPGEF3 | -2.21816 | 0.000001 | -1.07319 | 0.007724 |
| 51735 | RAPGEF6 | -1.04558 | 2.44E-06 | -0.97337 | 0.000001 |
| 65059 | RAPH1 | 1.785565 | 0.000001 | 0.685246 | 5.31E-05 |
| 5914 | RARA | 1.709691 | 0.000001 | 0.975349 | 0.000001 |
| 1.02E+08 | RARA-AS1 | -2.16582 | 0.000226 | 1.491895 | 0.000001 |
| 158158 | RASEF | 1.842081 | 0.000001 | 1.001562 | 0.000001 |
| 91608 | RASL10B | -1.34391 | 0.005527 | -0.82518 | 0.000001 |
| 9770 | RASSF2 | 3.242009 | 0.000001 | 1.284447 | 0.000001 |
| 83937 | RASSF4 | -2.9925 | 0.000001 | -0.60935 | 0.049827 |
| 83593 | RASSF5 | 1.527332 | 0.000001 | -2.17165 | 0.000001 |
| 55225 | RAVER2 | -1.71624 | 0.000001 | -1.79582 | 0.000001 |
| 64080 | RBKS | -1.33237 | 0.000754 | -0.99316 | 0.024229 |
| 5933 | RBL1 | 1.584937 | 0.000001 | -1.08842 | 0.000001 |
| 27303 | RBMS3 | -3.47079 | 0.000001 | 0.804731 | 0.000001 |
| 5947 | RBP1 | -1.7801 | 0.001353 | 1.178849 | 0.000001 |
| 1104 | RCC1 | 1.276242 | 0.000001 | -0.68499 | 0.000001 |
| 283248 | RCOR2 | -2.3042 | 7.67E-06 | -1.12772 | 5.08E-06 |
| 157506 | RDH10 | -1.47787 | 0.000001 | -0.68547 | 0.003241 |
| 5959 | RDH5 | -1.91607 | 0.000001 | -2.017 | 0.000001 |
| 51308 | REEP2 | -1.02043 | 0.000921 | -0.78494 | 2.46E-05 |
| 84957 | RELT | 2.540312 | 0.000001 | 0.726427 | 0.004686 |
| 54463 | RETREG1 | 1.722866 | 0.000001 | -1.85483 | 0.000001 |
| 5980 | REV3L | -1.54913 | 0.000001 | -0.98525 | 0.000001 |
| 5982 | RFC2 | 1.222625 | 0.000001 | -0.6215 | 0.000001 |
| 5983 | RFC3 | 1.934124 | 0.000001 | -0.71907 | 1.22E-06 |
| 5993 | RFX5 | -1.48872 | 0.000001 | -0.61927 | 1.19E-05 |
| 731220 | RFX8 | 1.24841 | 0.000001 | -1.36922 | 0.000001 |

| | | | | | |
|---|---|---|---|---|---|
| 28984 | RGCC | 2.185936 | 0.000469 | -2.06289 | 0.000001 |
| 9104 | RGN | -2.73324 | 0.000001 | -1.38586 | 2.24E-05 |
| 6001 | RGS10 | -1.4486 | 0.000001 | -1.01534 | 0.000001 |
| 8786 | RGS11 | -1.96442 | 0.000001 | -0.75797 | 0.003439 |
| 10636 | RGS14 | -1.71856 | 0.000001 | -1.27148 | 7.63E-05 |
| 6004 | RGS16 | 2.284826 | 0.000001 | 1.461466 | 0.000001 |
| 26575 | RGS17 | 1.212557 | 7.21E-06 | -1.38979 | 0.000001 |
| 5997 | RGS2 | 2.949688 | 0.000001 | -2.98685 | 0.000001 |
| 5999 | RGS4 | 3.065401 | 0.000001 | 0.834162 | 0.034672 |
| 9028 | RHBDL1 | -1.91294 | 4.32E-05 | -1.17806 | 0.009719 |
| 23221 | RHOBTB2 | -1.25374 | 0.000001 | -0.87279 | 0.000001 |
| 54509 | RHOF | -1.26436 | 0.000161 | -1.25297 | 0.000163 |
| 85415 | RHPN2 | 1.766299 | 7.03E-06 | 0.761517 | 0.003985 |
| 26150 | RIBC2 | 3.155499 | 0.000001 | -1.56322 | 9.55E-05 |
| 83547 | RILP | -1.03921 | 0.000001 | -1.1715 | 0.000001 |
| 85376 | RIMBP3 | -1.48394 | 0.000434 | -1.39617 | 0.000736 |
| 9610 | RIN1 | 2.203701 | 0.000001 | -2.3705 | 0.000001 |
| 11035 | RIPK3 | -2.3366 | 0.000001 | -1.04641 | 1.57E-05 |
| 9750 | RIPOR2 | -3.09568 | 0.000001 | -3.44757 | 0.000001 |
| 140876 | RIPOR3 | -3.18244 | 0.000001 | -1.62037 | 0.000001 |
| 6038 | RNASE4 | -1.48904 | 0.000001 | -0.76473 | 0.000001 |
| 6041 | RNASEL | -2.02756 | 0.000001 | -0.70187 | 2.34E-06 |
| 8153 | RND2 | -2.01403 | 0.000001 | -1.7651 | 0.000001 |
| 7732 | RNF112 | -2.25893 | 0.000001 | 1.802554 | 0.000001 |
| 114804 | RNF157 | -1.66785 | 0.000001 | -2.51711 | 0.000001 |
| 388591 | RNF207 | -2.00266 | 1.79E-05 | -0.83066 | 0.049931 |
| 22838 | RNF44 | -2.32688 | 0.000001 | 0.644426 | 1.46E-05 |
| 6092 | ROBO2 | -1.39567 | 0.000208 | -3.49088 | 0.000001 |
| 79641 | ROGDI | -2.03295 | 0.000001 | 0.791702 | 2.84E-05 |

| | | | | | |
|---|---|---|---|---|---|
| 6094 | ROM1 | -1.42597 | 0.000001 | -0.69913 | 0.01243 |
| 4919 | ROR1 | -1.04577 | 0.000001 | -1.9452 | 0.000001 |
| 6096 | RORB | 1.792899 | 0.005707 | -1.19825 | 0.026435 |
| 729215 | RPL32P29 | -2.84729 | 0.000001 | -1.03028 | 0.000966 |
| 54913 | RPP25 | 1.596527 | 0.000001 | -0.88566 | 0.000001 |
| 6196 | RPS6KA2 | -1.59415 | 0.000001 | -0.62318 | 0.000001 |
| 58528 | RRAGD | 1.406673 | 0.000001 | -0.99528 | 0.000001 |
| 9136 | RRP9 | 1.297049 | 0.000001 | 0.641107 | 3.13E-05 |
| 284654 | RSPO1 | -1.8127 | 0.003433 | -1.40077 | 0.02252 |
| 1.01E+08 | RTCA-AS1 | -2.60614 | 0.000001 | -0.87812 | 0.00736 |
| 219790 | RTKN2 | 1.69873 | 3.89E-06 | 1.595685 | 0.000001 |
| 146760 | RTN4RL1 | -1.48735 | 0.000929 | -2.71431 | 0.000001 |
| 64108 | RTP4 | -3.72826 | 0.000001 | -1.39074 | 0.006625 |
| 860 | RUNX2 | -1.131 | 0.000001 | 0.777813 | 0.003283 |
| 112611 | RWDD2A | -1.434 | 2.4E-05 | 0.606143 | 2.93E-05 |
| 59350 | RXFP1 | -2.55518 | 0.000001 | -1.77426 | 0.002362 |
| 6263 | RYR3 | 1.719554 | 0.001138 | 1.378867 | 0.006105 |
| 140576 | S100A16 | 1.947374 | 0.000001 | 0.697449 | 3.02E-06 |
| 6274 | S100A3 | 1.051486 | 0.000471 | -1.0955 | 0.000413 |
| 6275 | S100A4 | 1.005313 | 0.000001 | -1.9317 | 0.000001 |
| 6297 | SALL2 | -2.27676 | 0.000001 | -1.34885 | 0.000001 |
| 1.05E+08 | SAP30-DT | -2.21721 | 0.000001 | -0.99437 | 0.009245 |
| 54938 | SARS2 | -1.22946 | 0.000001 | -0.60696 | 2.38E-05 |
| 60485 | SAV1 | -1.20813 | 0.000001 | -0.68505 | 0.000001 |
| 949 | SCARB1 | -1.09576 | 0.000001 | -0.83401 | 0.000001 |
| 85477 | SCIN | -1.29165 | 0.000001 | -1.94956 | 0.000001 |
| 6323 | SCN1A | -1.20659 | 0.002825 | -1.4125 | 0.017244 |
| 6324 | SCN1B | -1.56276 | 0.000001 | -1.34653 | 0.000001 |
| 6326 | SCN2A | -2.28336 | 0.000001 | -1.78764 | 0.000001 |

| | | | | | |
|---|---|---|---|---|---|
| 6331 | SCN5A | 2.647925 | 0.000723 | -1.44093 | 3.56E-05 |
| 90507 | SCRN2 | -2.62818 | 0.000001 | -0.94476 | 0.000001 |
| 222663 | SCUBE3 | -1.19851 | 0.000536 | 1.458062 | 0.000001 |
| 6382 | SDC1 | -1.26677 | 0.000001 | 2.026432 | 0.000001 |
| 163859 | SDE2 | 1.094588 | 0.000001 | 0.646901 | 0.000001 |
| 221935 | SDK1 | -1.43305 | 7.03E-05 | -1.25852 | 0.000001 |
| 9717 | SEC14L5 | -2.03131 | 5.26E-06 | -2.7093 | 0.000001 |
| 6398 | SECTM1 | -2.04826 | 0.000001 | -3.39559 | 0.000001 |
| 23231 | SEL1L3 | 1.007577 | 3.85E-05 | 2.931909 | 0.000001 |
| 8991 | SELENBP1 | -2.01493 | 0.000001 | -1.889 | 0.000001 |
| 85465 | SELENOI | 1.105891 | 0.000001 | 0.625848 | 1.07E-05 |
| 6414 | SELENOP | -1.39488 | 0.000001 | -1.29803 | 0.000001 |
| 10371 | SEMA3A | -1.31545 | 5.03E-05 | -1.64371 | 0.000113 |
| 7869 | SEMA3B | -1.31461 | 0.000001 | -1.60871 | 0.000001 |
| 64218 | SEMA4A | -1.71892 | 0.003447 | -1.19911 | 0.019699 |
| 8482 | SEMA7A | 1.186245 | 0.000001 | 3.344416 | 0.000001 |
| 55964 | SEPTIN3 | -1.21085 | 0.000168 | -2.96636 | 0.000001 |
| 5414 | SEPTIN4 | -2.06533 | 0.000634 | -2.00366 | 0.000001 |
| 23157 | SEPTIN6 | -1.80852 | 0.000001 | 0.738267 | 2.22E-06 |
| 253190 | SERHL2 | -1.47981 | 0.001079 | -1.24595 | 0.000332 |
| 256987 | SERINC5 | 1.342249 | 0.000001 | 1.137344 | 0.000001 |
| 5055 | SERPINB2 | 3.231672 | 0.000001 | -1.96916 | 9.67E-06 |
| 5269 | SERPINB6 | -1.0333 | 0.000001 | 0.662794 | 0.000001 |
| 221756 | SERPINB9P1 | -3.49871 | 0.000001 | -2.31532 | 0.000001 |
| 5054 | SERPINE1 | 1.680513 | 0.000001 | 3.553446 | 0.000001 |
| 143686 | SESN3 | -2.75158 | 0.000001 | -1.10937 | 3.6E-05 |
| 6422 | SFRP1 | -1.28053 | 0.000001 | -2.24079 | 0.000001 |
| 6423 | SFRP2 | -1.37458 | 0.000001 | -2.02793 | 0.000001 |
| 6446 | SGK1 | 1.456745 | 0.000001 | 1.875614 | 0.000001 |

| | | | | | |
|---|---|---|---|---|---|
| 151648 | SGO1 | 1.55214 | 0.000248 | -0.90155 | 0.001745 |
| 10603 | SH2B2 | -1.78944 | 0.000001 | -0.94204 | 0.000209 |
| 400745 | SH2D5 | 1.968485 | 0.000001 | -0.8946 | 0.000001 |
| 6450 | SH3BGR | -1.59638 | 2.19E-06 | 1.251089 | 2.55E-06 |
| 80851 | SH3BP5L | 1.293309 | 0.000001 | 1.221943 | 0.000001 |
| 9644 | SH3PXD2A | -1.09837 | 0.000001 | 1.751934 | 0.000001 |
| 79628 | SH3TC2 | 2.963024 | 0.000001 | -1.2222 | 0.01063 |
| 53358 | SHC3 | 1.264322 | 0.000001 | -1.71187 | 0.000001 |
| 79801 | SHCBP1 | 1.404193 | 0.000001 | -1.06525 | 0.000001 |
| 55337 | SHFL | -2.80937 | 0.000001 | -0.75251 | 0.000001 |
| 85352 | SHISAL1 | 1.240988 | 0.000001 | -1.16412 | 0.000001 |
| 6474 | SHOX2 | -1.36202 | 0.000001 | -1.35226 | 0.000001 |
| 57619 | SHROOM3 | -1.20384 | 0.000269 | -1.29395 | 0.000001 |
| 59307 | SIGIRR | -2.11403 | 0.000001 | -1.42694 | 0.000001 |
| 6495 | SIX1 | -1.00421 | 3.3E-05 | -1.23475 | 0.000001 |
| 220134 | SKA1 | 1.356286 | 6.97E-06 | -1.24708 | 0.000001 |
| 221150 | SKA3 | 1.419171 | 1.44E-06 | -1.41503 | 0.000001 |
| 387640 | SKIDA1 | -2.96254 | 0.000001 | -1.75723 | 0.000001 |
| 56996 | SLC12A9 | -1.31982 | 0.000001 | -0.94119 | 0.000001 |
| 51296 | SLC15A3 | -1.85313 | 0.000001 | -1.38995 | 0.000001 |
| 6566 | SLC16A1 | 1.429805 | 0.000001 | 0.743141 | 0.000001 |
| 10560 | SLC19A2 | 1.188908 | 0.000001 | 1.840479 | 0.000001 |
| 6506 | SLC1A2 | 1.224614 | 0.000001 | -0.96617 | 2.23E-05 |
| 6507 | SLC1A3 | -1.20144 | 0.000001 | -1.68471 | 0.000001 |
| 6510 | SLC1A5 | -1.0593 | 0.000001 | 1.602165 | 0.000001 |
| 6512 | SLC1A7 | -1.05612 | 0.000392 | -0.69489 | 0.006159 |
| 6574 | SLC20A1 | 1.641006 | 0.000001 | 0.664899 | 3.99E-05 |
| 5002 | SLC22A18 | -1.58377 | 0.000001 | -1.08316 | 0.000001 |
| 6581 | SLC22A3 | 1.798652 | 1.1E-05 | 2.06492 | 0.000001 |

| | | | | | |
|---|---|---|---|---|---|
| 1468 | SLC25A10 | -1.73314 | 0.000001 | -1.0621 | 0.000001 |
| 8604 | SLC25A12 | -1.2759 | 0.000001 | -0.6642 | 0.000001 |
| 10165 | SLC25A13 | 1.560356 | 0.000001 | 1.049166 | 0.000001 |
| 11000 | SLC27A3 | -2.12269 | 0.000001 | -2.2748 | 0.000001 |
| 440584 | SLC2A1-AS1 | -1.8845 | 0.004053 | -1.16819 | 0.008896 |
| 154091 | SLC2A12 | -3.28149 | 0.000001 | -1.93182 | 0.000001 |
| 56731 | SLC2A4RG | -1.61011 | 0.000001 | -0.83105 | 0.000001 |
| 9906 | SLC35E2A | -1.81664 | 0.000001 | -1.15499 | 0.000001 |
| 728661 | SLC35E2B | -1.95299 | 0.000001 | -0.6685 | 0.000001 |
| 339665 | SLC35E4 | 1.854171 | 0.000001 | -0.83148 | 8.67E-06 |
| 222553 | SLC35F1 | 2.421909 | 0.005112 | 1.778645 | 0.00045 |
| 54733 | SLC35F2 | 1.727125 | 0.000001 | 0.677972 | 0.00077 |
| 206358 | SLC36A1 | 2.196914 | 0.000001 | 0.766757 | 1.38E-06 |
| 54020 | SLC37A1 | -1.67106 | 0.000105 | -1.17205 | 1.46E-05 |
| 81539 | SLC38A1 | 1.664529 | 0.000001 | 1.034226 | 0.000001 |
| 92745 | SLC38A5 | 1.238061 | 0.000001 | 1.983376 | 0.000001 |
| 64116 | SLC39A8 | -2.04215 | 0.000001 | -1.11325 | 0.000001 |
| 6520 | SLC3A2 | 2.356559 | 0.000001 | 0.786517 | 0.000001 |
| 30061 | SLC40A1 | -2.53975 | 0.000001 | -1.86585 | 0.000001 |
| 124935 | SLC43A2 | -2.56818 | 0.000001 | 1.233352 | 0.000001 |
| 50651 | SLC45A1 | -2.90087 | 0.000001 | -1.19422 | 0.000294 |
| 85414 | SLC45A3 | 1.016529 | 0.001393 | 0.748974 | 0.014681 |
| 283537 | SLC46A3 | -2.04083 | 0.000001 | 1.722535 | 0.000001 |
| 55244 | SLC47A1 | -1.09915 | 0.000001 | -2.36842 | 0.000001 |
| 84179 | SLC49A3 | -1.41809 | 0.000001 | 1.23128 | 0.000001 |
| 9497 | SLC4A7 | 1.552209 | 0.000001 | 0.81802 | 0.000001 |
| 55065 | SLC52A1 | -2.60269 | 8.09E-06 | -1.34879 | 0.027418 |
| 152078 | SLC66A1L | -1.43796 | 0.000001 | -1.30139 | 0.000001 |
| 130814 | SLC66A3 | -1.42685 | 0.000001 | -0.67055 | 0.000001 |

| | | | | | |
|---|---|---|---|---|---|
| 55117 | SLC6A15 | 2.028493 | 0.000001 | -1.42466 | 0.000001 |
| 23657 | SLC7A11 | 1.39901 | 0.000001 | 0.824532 | 0.000001 |
| 57709 | SLC7A14 | -1.34567 | 8.88E-05 | -4.07073 | 0.000001 |
| 9057 | SLC7A6 | 1.422172 | 0.000001 | 0.942474 | 0.000001 |
| 121456 | SLC9A7P1 | -1.72412 | 0.004581 | 1.049106 | 0.012408 |
| 285195 | SLC9A9 | -1.57313 | 0.000001 | -2.86408 | 0.000001 |
| 6578 | SLCO2A1 | 1.753775 | 0.000359 | 3.751209 | 0.000001 |
| 28231 | SLCO4A1 | 6.203951 | 0.000001 | -3.09426 | 0.000001 |
| 162394 | SLFN5 | -1.17656 | 0.000001 | 0.612195 | 0.000001 |
| 4088 | SMAD3 | -2.03908 | 0.000001 | -1.25633 | 0.000001 |
| 4091 | SMAD6 | -3.96442 | 0.000001 | -0.61955 | 3.02E-05 |
| 6595 | SMARCA2 | -1.93828 | 0.000001 | -0.89607 | 0.000001 |
| 6603 | SMARCD2 | -1.03274 | 0.000001 | -0.68991 | 0.000001 |
| 644596 | SMIM10L2B | -2.50647 | 0.000001 | -0.61092 | 0.003546 |
| 132332 | SMIM43 | -1.37771 | 9.64E-06 | 2.336909 | 0.000001 |
| 10924 | SMPDL3A | -1.11973 | 0.000001 | -0.89923 | 0.000001 |
| 64754 | SMYD3 | 1.185545 | 0.000001 | 0.654285 | 2.77E-06 |
| 6617 | SNAPC1 | 1.296018 | 0.000001 | 0.734886 | 0.000001 |
| 8420 | SNHG3 | 2.388394 | 0.000001 | 0.683095 | 0.000001 |
| 8303 | SNN | 1.683682 | 0.000001 | -1.03501 | 0.000001 |
| 92017 | SNX29 | -1.65573 | 0.000001 | -0.88544 | 0.000001 |
| 144481 | SOCS2-AS1 | -1.98555 | 0.000001 | -2.54257 | 0.000001 |
| 6648 | SOD2 | -1.93315 | 6.38E-06 | -1.45914 | 0.000001 |
| 10174 | SORBS3 | -1.1519 | 0.000001 | -1.0211 | 0.000001 |
| 6272 | SORT1 | -1.08571 | 0.000001 | 1.16299 | 0.000001 |
| 6664 | SOX11 | 2.412202 | 0.000001 | 0.95064 | 0.000001 |
| 6666 | SOX12 | -1.15871 | 0.000001 | -0.6713 | 0.000001 |
| 6659 | SOX4 | -2.18968 | 0.000001 | 1.225447 | 0.003516 |
| 6662 | SOX9 | -1.16023 | 0.000001 | 2.284951 | 0.000001 |

| | | | | | |
|---|---|---|---|---|---|
| 54558 | SPATA6 | -3.17186 | 0.000001 | -1.63084 | 0.000001 |
| 55812 | SPATA7 | -1.07133 | 0.000001 | -1.02651 | 0.000001 |
| 8877 | SPHK1 | 1.593792 | 0.000001 | 1.592241 | 0.000001 |
| 474343 | SPIN2B | -1.32659 | 1.67E-06 | -0.98711 | 0.000001 |
| 84501 | SPIRE2 | -1.78223 | 0.000001 | -1.14148 | 0.000001 |
| 6696 | SPP1 | 3.374196 | 0.000001 | 2.842818 | 0.000001 |
| 10252 | SPRY1 | -2.33147 | 0.000001 | -2.27071 | 0.000001 |
| 80176 | SPSB1 | 1.092972 | 0.000001 | 0.77227 | 0.000001 |
| 6711 | SPTBN1 | -1.1306 | 0.000001 | -0.91183 | 0.000001 |
| 58472 | SQOR | -1.25667 | 0.000001 | -0.66641 | 0.000001 |
| 10011 | SRA1 | 1.018448 | 0.000001 | 0.97477 | 0.000001 |
| 6720 | SREBF1 | -1.28006 | 0.000001 | -0.80573 | 0.000001 |
| 6728 | SRP19 | 1.12213 | 0.000001 | 0.690304 | 0.000001 |
| 140809 | SRXN1 | 1.555127 | 0.000001 | 1.1905 | 0.000001 |
| 136853 | SSC4D | 1.33132 | 0.000593 | 1.391114 | 0.00014 |
| 54961 | SSH3 | -1.84264 | 0.000001 | -0.84775 | 0.000001 |
| 8082 | SSPN | -2.12254 | 0.000001 | 0.692151 | 0.000001 |
| 1.01E+08 | ST3GAL6-AS1 | 1.346908 | 0.00234 | 1.054418 | 0.028963 |
| 6480 | ST6GAL1 | -1.79207 | 0.000763 | -0.99463 | 6.23E-05 |
| 6489 | ST8SIA1 | -1.57275 | 0.000001 | -3.79106 | 0.000001 |
| 55620 | STAP2 | -2.2065 | 3.12E-06 | -1.05449 | 0.000855 |
| 90627 | STARD13 | -1.7349 | 0.000001 | 1.262722 | 0.000001 |
| 6776 | STAT5A | -1.59442 | 0.000001 | -0.83517 | 0.000001 |
| 55240 | STEAP3 | -1.0077 | 1.16E-05 | -1.29684 | 0.000001 |
| 79689 | STEAP4 | -5.25634 | 0.000001 | -3.46235 | 0.000001 |
| 50861 | STMN3 | -1.23486 | 0.000001 | -0.80693 | 0.000155 |
| 2040 | STOM | -1.08809 | 0.000001 | -0.91316 | 0.000001 |
| 11037 | STON1 | -2.83996 | 0.000001 | -0.72916 | 0.002313 |
| 85439 | STON2 | 3.420808 | 0.000001 | -0.87383 | 0.033728 |

| | | | | | |
|---|---|---|---|---|---|
| 57464 | STRIP2 | 2.246654 | 0.000001 | -0.95365 | 0.002447 |
| 375057 | STUM | -3.23383 | 8.71E-05 | -2.39733 | 2.29E-05 |
| 8677 | STX10 | -1.0686 | 0.000001 | -0.83263 | 0.000001 |
| 112755 | STX1B | -1.969 | 5.28E-06 | 0.982732 | 0.001886 |
| 29091 | STXBP6 | 1.763871 | 0.000001 | -2.11173 | 0.000001 |
| 8801 | SUCLG2 | -1.28673 | 0.000001 | -0.74312 | 0.000001 |
| 1.02E+08 | SUGCT-AS1 | 2.641123 | 0.000001 | 1.059364 | 0.003185 |
| 57653 | SUGT1P4-STRA6LP-CCDC180 | -1.67135 | 0.000339 | -0.81413 | 0.014152 |
| 23213 | SULF1 | -1.06627 | 0.000432 | 2.043995 | 0.000001 |
| 25830 | SULT4A1 | -1.75515 | 0.000207 | -1.9482 | 0.00088 |
| 6839 | SUV39H1 | 1.050262 | 0.000001 | -0.74586 | 0.000107 |
| 79987 | SVEP1 | -2.38983 | 0.000001 | -0.95809 | 2.23E-06 |
| 6840 | SVIL | -1.54628 | 0.000001 | -1.37321 | 0.000001 |
| 55638 | SYBU | 1.085035 | 4.98E-06 | 1.324055 | 0.000001 |
| 79953 | SYNDIG1 | -1.89085 | 0.000883 | 4.036853 | 0.000001 |
| 9145 | SYNGR1 | -1.10774 | 0.000001 | -1.78068 | 0.000001 |
| 284612 | SYPL2 | 1.678368 | 0.000001 | -1.73288 | 0.000001 |
| 255928 | SYT14 | 1.157259 | 0.000001 | 1.242306 | 0.000001 |
| 6863 | TAC1 | 3.50261 | 0.000001 | 1.91307 | 0.000205 |
| 6884 | TAF13 | 1.403314 | 0.000001 | 1.00035 | 0.000001 |
| 55080 | TAPBPL | -1.15976 | 0.000001 | -0.77224 | 0.000001 |
| 1.08E+08 | TARS1-DT | -1.9692 | 0.002356 | 0.915339 | 0.004528 |
| 80835 | TAS1R1 | -2.78771 | 5.68E-06 | -3.4827 | 0.000001 |
| 30851 | TAX1BP3 | -1.4089 | 0.000001 | 1.091923 | 0.000001 |
| 9882 | TBC1D4 | -1.61609 | 0.000001 | -0.72739 | 0.000001 |
| 11138 | TBC1D8 | -2.38213 | 0.000001 | -2.87869 | 0.000001 |
| 9096 | TBX18 | -1.40809 | 0.000001 | -0.92651 | 0.000001 |

| | | | | | |
|---|---|---|---|---|---|
| 6915 | TBXA2R | -1.40899 | 4.49E-05 | -1.19592 | 0.000474 |
| 123036 | TC2N | 2.76292 | 0.000001 | 1.058073 | 0.009979 |
| 285966 | TCAF2 | -1.169 | 0.003752 | -1.54153 | 0.000107 |
| 6920 | TCEA3 | -2.92841 | 0.000001 | -2.4937 | 0.000001 |
| 10312 | TCIRG1 | -1.43236 | 0.000001 | -0.6592 | 0.000001 |
| 6996 | TDG | 1.539638 | 0.000001 | 1.088881 | 0.000001 |
| 283643 | TEDC1 | 2.561467 | 0.000001 | -0.63187 | 0.00048 |
| 55714 | TENM3 | -1.01824 | 0.000001 | 2.22769 | 0.000001 |
| 26011 | TENM4 | -2.33854 | 0.000001 | 0.839878 | 3.07E-05 |
| 26136 | TES | 2.933434 | 0.000001 | 1.54705 | 0.000001 |
| 51368 | TEX264 | -1.27185 | 0.000001 | -0.66808 | 0.000001 |
| 7023 | TFAP4 | -1.53098 | 0.000001 | -1.47961 | 0.000001 |
| 7029 | TFDP2 | -1.53645 | 0.000001 | -0.67213 | 0.000001 |
| 7035 | TFPI | -1.81528 | 0.000001 | -1.51303 | 0.000001 |
| 7043 | TGFB3 | -1.18587 | 0.000001 | -1.30401 | 3.02E-06 |
| 7049 | TGFBR3 | -1.00767 | 1.36E-06 | -1.8506 | 0.000001 |
| 1.01E+08 | TGFBR3L | -1.80104 | 0.000001 | -1.1812 | 0.000699 |
| 79725 | THAP9 | -1.65191 | 1.23E-05 | -0.69498 | 0.00728 |
| 7056 | THBD | 1.303218 | 0.000001 | -1.32448 | 0.004895 |
| 7057 | THBS1 | -3.82794 | 0.000001 | 1.206134 | 0.000001 |
| 1.19E+08 | THBS1-AS1 | -3.97664 | 0.000001 | 0.737938 | 0.008064 |
| 1.19E+08 | THBS1-IT1 | -1.16056 | 5.45E-05 | 1.404731 | 0.000001 |
| 51337 | THEM6 | -1.53123 | 0.000001 | -1.99022 | 0.000001 |
| 7067 | THRA | -1.13742 | 0.000001 | -0.74595 | 0.000001 |
| 7068 | THRB | -1.10744 | 0.000001 | -1.2475 | 0.000001 |
| 10440 | TIMM17A | 1.405779 | 0.000001 | 0.687335 | 7.11E-06 |
| 1.13E+08 | TIMM23B-AGAP6 | 1.504688 | 0.000001 | -0.8707 | 0.007275 |
| 7079 | TIMP4 | 1.919848 | 0.000001 | -2.09201 | 0.000001 |
| 7088 | TLE1 | 1.520044 | 0.000001 | -0.62901 | 4.22E-05 |

| | | | | | |
|---|---|---|---|---|---|
| 7097 | TLR2 | -1.68497 | 0.008784 | -3.25974 | 0.000001 |
| 7098 | TLR3 | -3.10855 | 0.000001 | -1.83241 | 5.47E-06 |
| 7099 | TLR4 | 1.319366 | 0.000001 | 0.866331 | 1.71E-05 |
| 7108 | TM7SF2 | -2.54945 | 0.000001 | -0.76069 | 0.000001 |
| 113277 | TMEM106A | -1.83645 | 1.53E-05 | 0.914095 | 0.000001 |
| 84314 | TMEM107 | -1.15481 | 1.29E-05 | -1.09112 | 1.13E-06 |
| 338773 | TMEM119 | 1.605939 | 0.000001 | -1.36931 | 0.000001 |
| 441027 | TMEM150C | -2.35944 | 0.000001 | -1.97546 | 0.000001 |
| 25907 | TMEM158 | 2.371772 | 0.000001 | -3.05138 | 0.000001 |
| 728229 | TMEM191B | -2.0487 | 2.15E-05 | -1.56679 | 0.002022 |
| 645426 | TMEM191C | -1.54125 | 0.001909 | -1.04475 | 0.027306 |
| 440104 | TMEM198B | -4.04303 | 0.000001 | -0.60277 | 2.61E-05 |
| 388335 | TMEM220 | -1.28034 | 0.000001 | -0.87043 | 0.000001 |
| 161145 | TMEM229B | -3.55167 | 0.000001 | -2.7938 | 0.000001 |
| 339453 | TMEM240 | -1.34054 | 0.000001 | -0.9688 | 2.03E-06 |
| 84866 | TMEM25 | -1.45458 | 0.000001 | -0.64649 | 0.001795 |
| 219623 | TMEM26 | -1.39651 | 0.002333 | -2.82243 | 0.000001 |
| 59353 | TMEM35A | 1.098834 | 3E-06 | -2.71923 | 0.000001 |
| 757 | TMEM50B | -1.33565 | 0.000001 | -0.85259 | 0.000001 |
| 55092 | TMEM51 | 2.18481 | 0.000001 | -1.06057 | 4.12E-05 |
| 79639 | TMEM53 | -1.41851 | 0.000001 | -0.96328 | 1.26E-05 |
| 9725 | TMEM63A | -1.37859 | 0.000001 | -0.95694 | 0.000001 |
| 137835 | TMEM71 | 1.427214 | 0.000978 | -1.86538 | 0.000722 |
| 51754 | TMEM8B | -2.04232 | 0.000001 | -0.74302 | 0.000001 |
| 641649 | TMEM91 | -1.10945 | 0.007989 | 1.645058 | 1.54E-05 |
| 27346 | TMEM97 | -1.76644 | 0.000001 | -1.05618 | 0.000001 |
| 7111 | TMOD1 | 2.131125 | 1.28E-06 | 1.649822 | 0.000001 |
| 3371 | TNC | 1.982746 | 0.000001 | 1.28924 | 0.000001 |
| 25816 | TNFAIP8 | -2.20591 | 0.000001 | -0.88468 | 0.000001 |

| | | | | | |
|---|---|---|---|---|---|
| 126282 | TNFAIP8L1 | -2.36868 | 0.000001 | -0.75816 | 1.02E-05 |
| 8797 | TNFRSF10A | 2.387063 | 0.000001 | -0.86086 | 0.000421 |
| 389641 | TNFRSF10A-DT | -1.72294 | 6.39E-05 | -1.27525 | 0.012833 |
| 8794 | TNFRSF10C | 1.009185 | 0.00015 | -1.59669 | 1.95E-06 |
| 4982 | TNFRSF11B | -2.60994 | 0.000001 | -2.61666 | 0.000001 |
| 51330 | TNFRSF12A | 1.974146 | 0.000001 | 1.477277 | 0.000001 |
| 7133 | TNFRSF1B | -1.6619 | 0.000001 | -2.02245 | 0.000001 |
| 8743 | TNFSF10 | -4.4583 | 0.000001 | -1.48872 | 0.010685 |
| 8742 | TNFSF12 | -1.71808 | 0.000001 | -1.27178 | 0.000001 |
| 7292 | TNFSF4 | -1.34859 | 0.0015 | 1.405383 | 0.000001 |
| 7145 | TNS1 | -1.08587 | 0.000001 | 1.7895 | 0.000001 |
| 7148 | TNXB | 2.078874 | 9.88E-05 | -2.39026 | 0.000001 |
| 1E+08 | TOMM40P4 | -2.34382 | 0.000001 | 0.921429 | 0.00021 |
| 4796 | TONSL | 1.068133 | 0.000001 | -1.07861 | 0.000001 |
| 64222 | TOR3A | -1.46428 | 0.000001 | -0.69008 | 0.000001 |
| 54863 | TOR4A | 3.147884 | 0.000001 | -1.01851 | 0.000001 |
| 9537 | TP53I11 | -1.75534 | 0.000001 | 0.647216 | 0.000001 |
| 9540 | TP53I3 | -1.02214 | 0.000001 | 1.404543 | 0.000001 |
| 7168 | TPM1 | -1.00692 | 0.000001 | 3.388652 | 0.000001 |
| 7169 | TPM2 | -1.31864 | 0.000001 | 1.31123 | 0.000001 |
| 51673 | TPPP3 | 8.361435 | 0.000001 | -1.34264 | 0.00541 |
| 8460 | TPST1 | -1.71467 | 0.000001 | 0.9217 | 0.000001 |
| 28755 | TRAC | 2.360467 | 0.004892 | 2.435893 | 0.000001 |
| 10293 | TRAIP | 1.172608 | 0.000204 | -0.83167 | 0.001134 |
| 22906 | TRAK1 | 1.040374 | 0.000001 | 1.012316 | 0.000001 |
| 9881 | TRANK1 | -2.69566 | 0.000001 | -0.7443 | 1.83E-05 |
| 83696 | TRAPPC9 | -1.82188 | 0.000001 | -0.68861 | 9.15E-06 |
| 11277 | TREX1 | -2.94422 | 0.000001 | -0.96628 | 2.1E-06 |
| 10221 | TRIB1 | 1.62215 | 0.000001 | 0.998407 | 0.000001 |

| | | | | | |
|---|---|---|---|---|---|
| 9865 | TRIL | -3.28832 | 0.000001 | -1.90837 | 0.000001 |
| 9830 | TRIM14 | -2.59756 | 0.000001 | -1.19037 | 0.000001 |
| 23321 | TRIM2 | -2.43029 | 0.000001 | -0.96549 | 0.000001 |
| 10346 | TRIM22 | -1.40375 | 0.000001 | -0.95521 | 0.000001 |
| 53840 | TRIM34 | -3.25014 | 0.000001 | -1.04039 | 5.84E-05 |
| 140691 | TRIM69 | -2.19182 | 0.000001 | -0.64863 | 1.15E-05 |
| 81786 | TRIM7 | 1.936863 | 2.06E-06 | -1.62997 | 1.22E-06 |
| 9319 | TRIP13 | 1.751416 | 0.000001 | -0.92684 | 0.000001 |
| 10024 | TROAP | 1.249145 | 0.001552 | -0.90632 | 0.000535 |
| 8989 | TRPA1 | -3.41449 | 0.000001 | -2.07351 | 0.000001 |
| 7222 | TRPC3 | -1.64563 | 0.009292 | -1.45624 | 0.000001 |
| 7225 | TRPC6 | -2.24653 | 0.000001 | -2.43076 | 0.000001 |
| 10194 | TSHZ1 | -2.36876 | 0.000001 | -0.90497 | 0.000001 |
| 128553 | TSHZ2 | -1.50546 | 0.000001 | -1.77353 | 0.000001 |
| 85480 | TSLP | 1.810503 | 3.47E-06 | 1.033688 | 0.01742 |
| 83882 | TSPAN10 | -1.32964 | 0.000001 | -1.54399 | 0.000001 |
| 27075 | TSPAN13 | 2.891247 | 0.000001 | 2.83902 | 0.000001 |
| 10077 | TSPAN32 | -2.67099 | 0.001779 | -1.01518 | 0.049578 |
| 10867 | TSPAN9 | -1.94828 | 0.000001 | -0.91102 | 0.000001 |
| 9256 | TSPOAP1 | -3.22338 | 0.000001 | -1.65204 | 0.000001 |
| 150737 | TTC30B | -1.47002 | 5.13E-06 | -1.01646 | 5.16E-06 |
| 123016 | TTC8 | -1.61111 | 0.000001 | 0.674887 | 0.000001 |
| 23508 | TTC9 | 1.870873 | 0.000235 | 1.71352 | 0.000001 |
| 7272 | TTK | 1.183761 | 0.001337 | -1.31349 | 0.000001 |
| 7280 | TUBB2A | 1.571532 | 0.000001 | 1.047042 | 0.000001 |
| 347733 | TUBB2B | 1.624511 | 0.000001 | 1.435997 | 0.000001 |
| 84617 | TUBB6 | 1.763023 | 0.000001 | 0.823534 | 0.000001 |
| 56995 | TULP4 | 1.219943 | 0.000001 | 0.762978 | 0.000001 |
| 7291 | TWIST1 | -1.16113 | 0.000001 | -1.48926 | 0.000001 |

| | | | | | |
|---|---|---|---|---|---|
| 117581 | TWIST2 | -1.3842 | 0.000001 | -1.56781 | 0.000001 |
| 494514 | TYMSOS | -2.47139 | 1.08E-05 | -1.79463 | 0.000433 |
| 219743 | TYSND1 | -1.37214 | 4.43E-05 | -1.44457 | 0.000001 |
| 1.03E+08 | U2AF1L5 | 1.712125 | 0.000001 | 0.871158 | 6.84E-05 |
| 55075 | UACA | -2.05517 | 0.000001 | 0.601175 | 0.000001 |
| 6675 | UAP1 | 1.653932 | 0.000001 | 1.192659 | 0.000001 |
| 7318 | UBA7 | -4.1835 | 0.000001 | -1.71646 | 0.000001 |
| 124402 | UBALD1 | -2.25331 | 0.000001 | 0.895047 | 0.000001 |
| 11065 | UBE2C | 1.996343 | 0.000001 | -0.91902 | 6.76E-06 |
| 9246 | UBE2L6 | -2.3249 | 0.000001 | -0.73823 | 0.000001 |
| 84993 | UBL7 | -1.2434 | 0.000001 | 0.675975 | 0.000001 |
| 7351 | UCP2 | -1.68566 | 4.54E-05 | 1.198942 | 3.64E-06 |
| 7368 | UGT8 | 2.473299 | 0.005837 | -1.24544 | 0.026183 |
| 29128 | UHRF1 | 1.568088 | 0.000001 | -0.81553 | 0.000001 |
| 54887 | UHRF1BP1 | -1.0321 | 0.001335 | 1.25903 | 0.000001 |
| 80329 | ULBP1 | 1.466998 | 0.000107 | 1.881101 | 0.000001 |
| 80328 | ULBP2 | 1.277931 | 6.41E-06 | 1.734511 | 0.000001 |
| 79465 | ULBP3 | 1.321417 | 0.000001 | -0.73464 | 0.00014 |
| 8408 | ULK1 | -3.12858 | 0.000001 | 0.935546 | 0.000001 |
| 8633 | UNC5C | 3.204062 | 6.27E-05 | 1.902204 | 0.000245 |
| 81622 | UNC93B1 | -1.97038 | 0.000001 | -2.05067 | 0.000001 |
| 7374 | UNG | -1.67509 | 0.000001 | -0.68566 | 0.000001 |
| 7378 | UPP1 | 3.422285 | 0.000001 | 1.021466 | 0.000001 |
| 219333 | USP12 | 1.006964 | 0.000001 | 0.619034 | 0.000001 |
| 158880 | USP51 | -1.4752 | 0.000001 | -0.62049 | 0.000595 |
| 10090 | UST | -1.20707 | 0.000001 | -1.11683 | 0.000001 |
| 8674 | VAMP4 | -1.76852 | 0.000001 | -0.84898 | 0.000001 |
| 1462 | VCAN | -1.23137 | 0.000001 | 2.27339 | 0.000001 |
| 7421 | VDR | -2.88447 | 0.000001 | 1.582585 | 0.000001 |

| | | | | | |
|---|---|---|---|---|---|
| 7422 | VEGFA | 1.225285 | 0.000001 | 2.30772 | 0.000001 |
| 5212 | VIT | -1.50022 | 0.000001 | -1.36036 | 0.000001 |
| 7436 | VLDLR | 1.532961 | 0.000001 | 1.123978 | 0.000001 |
| 155382 | VPS37D | -2.10562 | 0.000001 | -1.14708 | 0.000001 |
| 64856 | VWA1 | -1.77374 | 0.000001 | -1.86056 | 0.000001 |
| 4013 | VWA5A | -3.59459 | 0.000001 | -0.66254 | 0.011332 |
| 375690 | WASH5P | 1.542944 | 0.000001 | 0.739822 | 0.000118 |
| 115825 | WDFY2 | 1.123965 | 0.000001 | 1.102534 | 0.000001 |
| 404201 | WDFY3-AS2 | -1.15016 | 1.96E-06 | -0.63266 | 0.000679 |
| 11169 | WDHD1 | 1.594446 | 0.000001 | -0.84881 | 0.000001 |
| 9948 | WDR1 | 1.025482 | 0.000001 | 0.606716 | 0.000001 |
| 114987 | WDR31 | -3.95299 | 0.000001 | -0.63359 | 0.000631 |
| 84058 | WDR54 | -2.00887 | 0.000001 | -0.74489 | 0.000001 |
| 284403 | WDR62 | 2.751366 | 0.000001 | -1.12155 | 0.000001 |
| 79968 | WDR76 | 1.665473 | 0.000001 | -1.07827 | 0.000001 |
| 7473 | WNT3 | -1.34295 | 1.76E-06 | -0.69914 | 0.01757 |
| 7483 | WNT9A | 1.607848 | 1.79E-06 | -0.74163 | 0.000207 |
| 55884 | WSB2 | 1.800018 | 0.000001 | 0.67526 | 0.000001 |
| 54739 | XAF1 | -3.2791 | 0.000001 | 1.278014 | 0.000001 |
| 7508 | XPC | 1.48891 | 0.000001 | -0.6439 | 0.000001 |
| 7516 | XRCC2 | 2.542198 | 0.000001 | -0.95554 | 6.39E-06 |
| 7517 | XRCC3 | 1.521563 | 0.000001 | -0.69749 | 0.001378 |
| 10897 | YIF1A | -1.17025 | 0.000001 | 0.887392 | 0.000001 |
| 219539 | YPEL4 | -4.11655 | 0.000001 | -1.33718 | 0.000843 |
| 79693 | YRDC | 2.348041 | 0.000001 | 1.20074 | 0.000001 |
| 729013 | ZBED5-AS1 | -2.95035 | 0.000001 | -0.9685 | 0.000113 |
| 221527 | ZBTB12 | -1.2413 | 2.81E-06 | -1.09426 | 0.000001 |
| 219654 | ZCCHC24 | -1.27819 | 0.000001 | -0.63534 | 0.000001 |
| 254887 | ZDHHC23 | -1.84878 | 2.95E-06 | -1.14712 | 0.000399 |

| | | | | | |
|---|---|---|---|---|---|
| 90637 | ZFAND2A | 2.552527 | 0.000001 | 0.684536 | 4.94E-05 |
| 1E+08 | ZFHX4-AS1 | -2.32689 | 3.23E-05 | -0.96989 | 0.016091 |
| 57677 | ZFP14 | -1.30102 | 2.6E-06 | -0.71768 | 0.000413 |
| 139735 | ZFP92 | 3.788793 | 0.000001 | 0.716626 | 0.020521 |
| 1E+08 | ZGLP1 | 1.801183 | 0.000001 | 1.01591 | 0.006277 |
| 55345 | ZGRF1 | 1.350426 | 0.000001 | -1.10385 | 1.25E-06 |
| 57178 | ZMIZ1 | -1.53867 | 0.000001 | 0.666522 | 0.000001 |
| 7754 | ZNF204P | -2.10017 | 0.001156 | -1.32254 | 0.005476 |
| 7761 | ZNF214 | -1.08153 | 8.29E-05 | -1.05854 | 0.000144 |
| 7764 | ZNF217 | -1.60944 | 0.000001 | -0.63794 | 2.58E-06 |
| 113835 | ZNF257 | 2.412165 | 1.3E-05 | -0.71364 | 0.041948 |
| 11179 | ZNF277 | -1.01363 | 0.000001 | -0.91991 | 0.000001 |
| 150142 | ZNF295-AS1 | 3.136631 | 0.000001 | 1.67666 | 0.000001 |
| 134466 | ZNF300P1 | 1.743825 | 0.000001 | 0.759061 | 0.026512 |
| 22891 | ZNF365 | 2.31183 | 0.000001 | 2.364843 | 0.000001 |
| 195828 | ZNF367 | 2.413405 | 0.000001 | -1.88249 | 0.000001 |
| 55893 | ZNF395 | -2.89613 | 0.000001 | -1.57883 | 0.000001 |
| 79797 | ZNF408 | 1.654446 | 0.000001 | 0.791848 | 0.000001 |
| 90333 | ZNF468 | 2.12291 | 0.000001 | 1.041862 | 0.008554 |
| 57615 | ZNF492 | 1.6494 | 0.005134 | -0.91139 | 0.025625 |
| 147807 | ZNF524 | -1.59018 | 0.000001 | -0.63256 | 2.25E-06 |
| 147837 | ZNF563 | -1.08164 | 0.007734 | -0.80295 | 0.025746 |
| 51385 | ZNF589 | 1.055766 | 0.000001 | -1.60917 | 0.000001 |
| 57507 | ZNF608 | -1.23723 | 4.5E-05 | -1.45305 | 0.000001 |
| 90874 | ZNF697 | 1.939519 | 0.000001 | 0.725013 | 0.000001 |
| 619279 | ZNF704 | -3.12978 | 0.000001 | -0.79152 | 0.008243 |
| 730087 | ZNF726 | 1.28301 | 0.001823 | -0.96792 | 0.024927 |
| 284371 | ZNF841 | 1.598759 | 0.000001 | 0.797988 | 0.000001 |
| 169834 | ZNF883 | -1.03278 | 0.004201 | -0.72291 | 0.033155 |

| | | | | | |
|---|---|---|---|---|---|
| 7643 | ZNF90 | 2.884359 | 0.000001 | -0.82853 | 0.043155 |
| 11130 | ZWINT | 1.601078 | 0.000001 | -1.20729 | 0.000001 |

Table S4 Common pathways enriched in the meta-analysis of both datasets.

Common pathways in both datasets and p-value < 0.01 were shown. The number in the table indicates the differentially expressed genes belonging to individual pathway and the total number of genes were shown in bracket from same pathway. LiCl lane: Differentially regulated genes by LiCl treated SSc fibroblast; TGF-β1 lane: TGF-β1 treated foreskin fibroblast (GSE232435)

| entity | name | DE genes (Total) LiCl | p-value (LiCl) | DE genes (Total) TGF-β1 | p-value (TGF-β1) |
|---|---|---|---|---|---|
| 10474 | Basal cell carcinoma | 28 (54) | 0.0006 | 26 (55) | 0.0013 |
| 10276 | Calcium signaling pathway | 69 (185) | 0.0179 | 76 (189) | 0.0000 |
| 10465 | Chemical carcinogenesis - receptor activation | 56 (152) | 0.0039 | 56 (146) | 0.0009 |

| | | | | | |
|---|---|---|---|---|---|
| 10279 | Cytokine-cytokine receptor interaction | 86 (186) | 0.0000 | 79 (164) | 0.0000 |
| 10193 | Glutathione metabolism | 24 (46) | 0.0020 | 22 (47) | 0.0038 |
| 10204 | Glycosaminoglycan biosynthesis - chondroitin sulfate / dermatan sulfate | 11 (20) | 0.0306 | 10 (19) | 0.0274 |
| 10242 | Metabolic pathways | 419 (1272) | 0.0001 | 360 (1245) | 0.0018 |
| 10288 | Neuroactive ligand-receptor interaction | 59 (166) | 0.0452 | 73 (156) | 0.0000 |
| 10459 | Pathways in cancer | 153 (454) | 0.0042 | 175 (457) | 0.0000 |
| 10265 | PPAR signaling pathway | 23 (55) | 0.0332 | 26 (55) | 0.0013 |
| 10491 | Rheumatoid arthritis | 35 (69) | 0.0003 | 23 (64) | 0.0060 |
| 10460 | Transcriptional misregulation in cancer | 55 (152) | 0.0180 | 48 (146) | 0.0041 |
| 10280 | Viral protein interaction with cytokine and cytokine receptor | 35 (59) | 0.0000 | 21 (46) | 0.0120 |

Figure S1

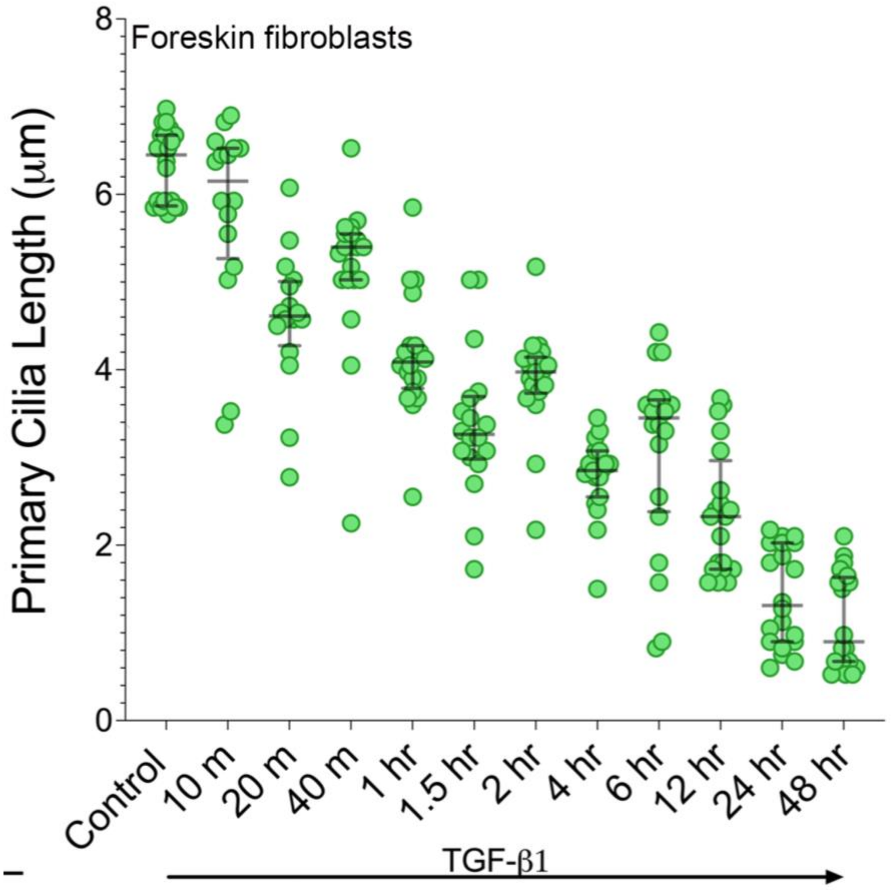

**Supplementary Figure S1. TGF-β1 induced time-dependent PC length shortening.** TGF-β1 induced time-dependent PC length (mean ± SD from 20 determinations) shortening measured over time by treating Foreskin fibroblasts with TGF-β1(10ng/ml) for different time periods (up to 48 hours) after serum starvation (0.1%BSA, ON), followed by IF with antibodies against Arl13b. The lengths of the PC were measured from the 3D reconstruction of the Z-volumes using ImageJ.

Figure S2

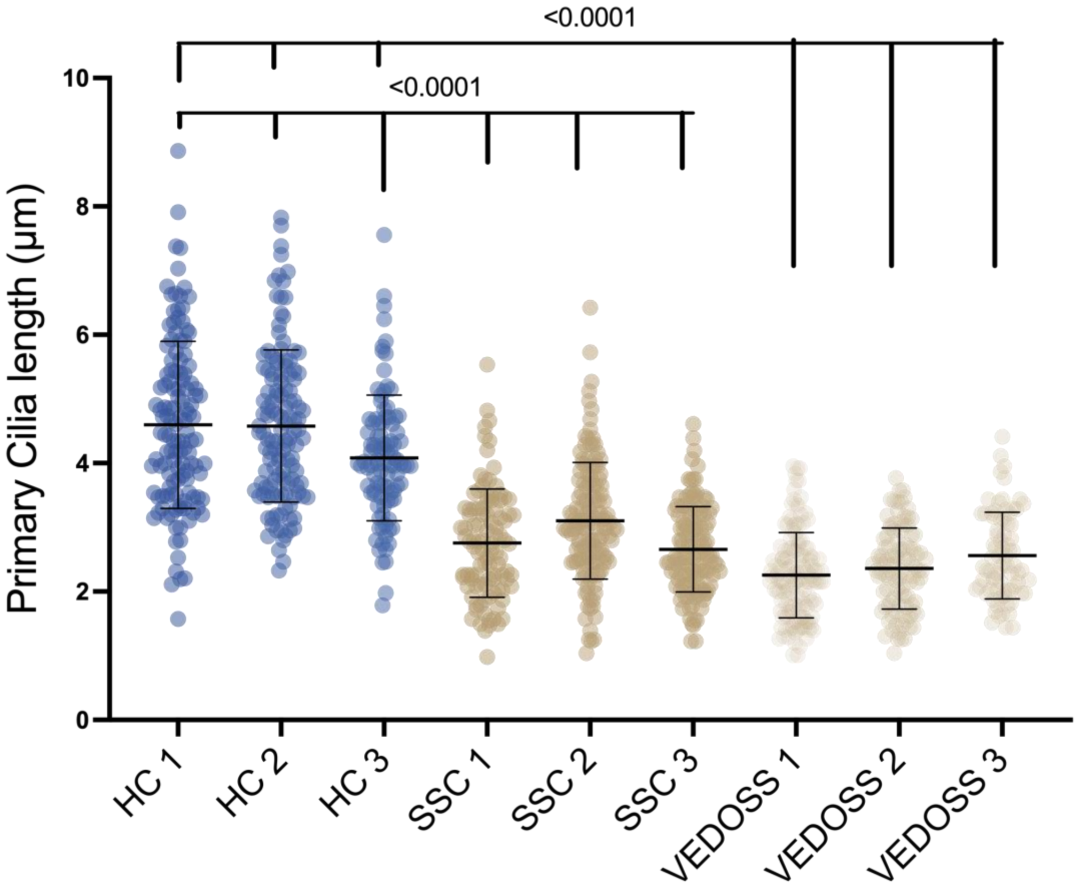

**Supplementary Figure S2. Primary cilia length reduced in skin fibroblasts with very early diagnosis of SSc (VEDOSS).** Primary dermal fibroblasts at ultra-confluence were starved in DMEM media containing 0.5% FBS for 48 h, fixed and subjected to immunofluorescence for Acetylated Alpha-Tubulin antibody. Length of cilia was determined in 100 cells per cell line. Mean cilia length (μm) for each biological donor (HC, SSc, VEDOSS; n=3,3,3). One-way ANOVA followed by Sidak's multiple comparison test.

Figure S3

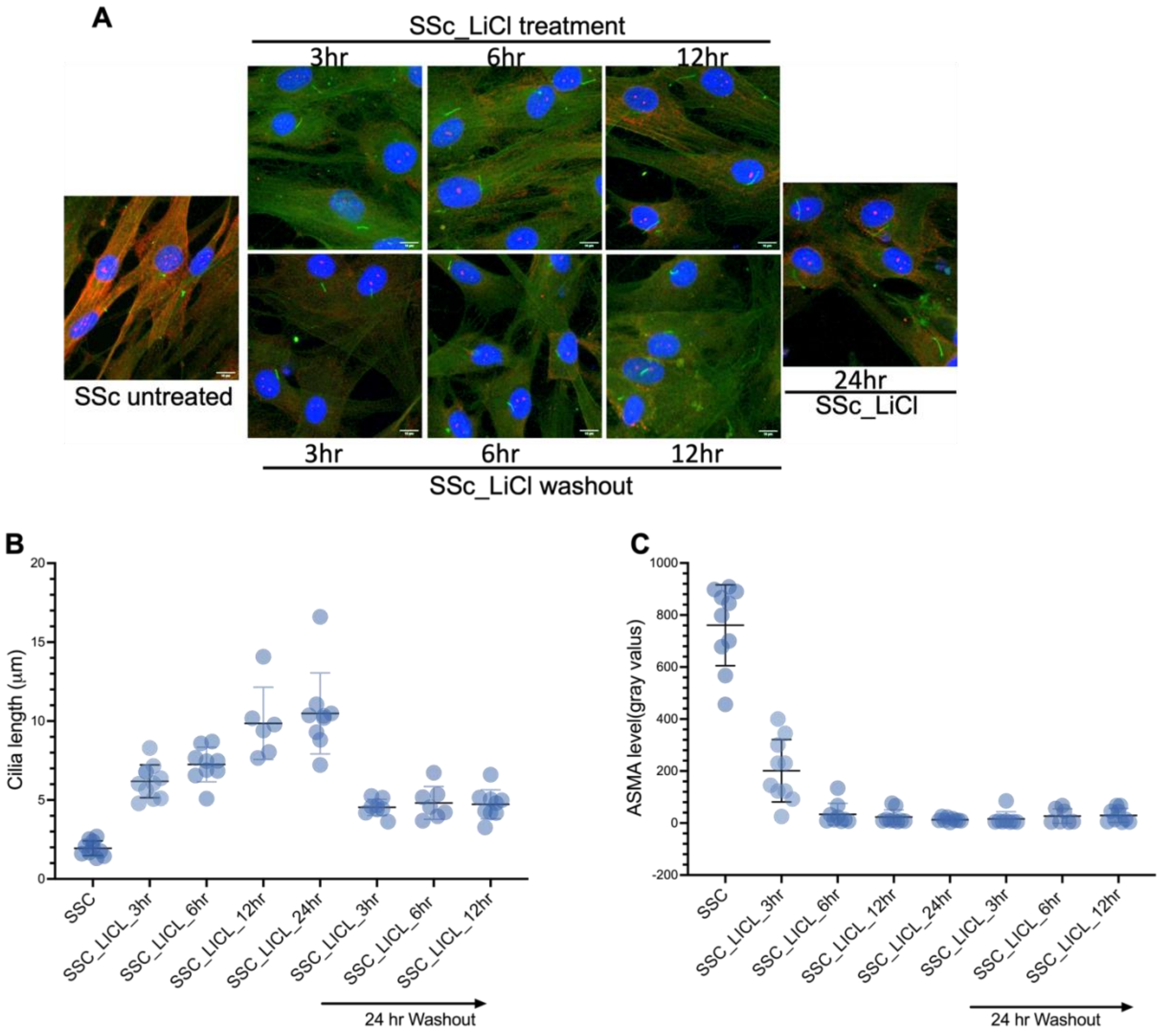

**Supplementary Figure S3. LiCl-mediated ciliary length restoration is stable and irreversible.** Confluent SSc skin fibroblasts from were incubated in media with 0.1% BSA for 12 hr and then incubated in media with LiCl (50mM). LiCl was washed out by media change in one of the batches and cultures were subjected to an additional 24 hr incubation. All cells were fixed and subjected to IF against antibodies to Arl13(green) and ASMA (red). (A) the leftmost image represents untreated cells, while the rightmost image represents 24 hr LiCl-treated cells. The middle lower panel represents cells subjected to LiCl washout followed by additional 24 hr incubation, while the upper panel represents cells without LiCl washout. Cells were imaged with Leica SP8 confocal microscope with z-volume. The scale bar represents 10 μm. (B)The lengths of the PC were measured from the 3D reconstruction of the Z-volumes using Imaris. The graphs represent the quantification of PC length. Each dot represents the quantification of PC length from a different cell. C) The graphs represent the quantification of the ASMA level, which was determined using ImageJ.

**Figure S4**

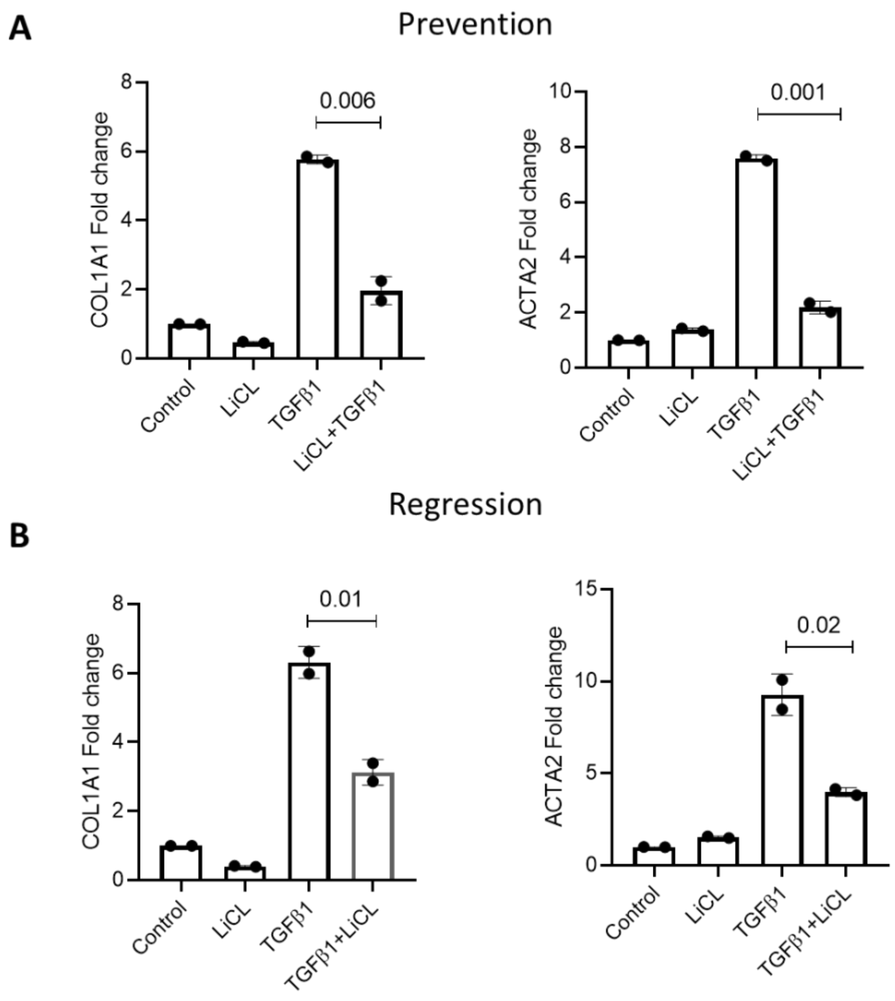

**Supplementary Figure S4: LiCl treatment both prevents and reverses TGF-β1-induced changes.** (A) Foreskin fibroblasts were 0.1% BSA starved for 24 hr, followed by LiCl pretreatment for 60 min, and treatment with TGF-β1(10 ng/ml) for 24 hr (preventive study). Quantification of ACTA2, COL1A1 via qPCR. Unpaired t test (n=2 independent experiment) (B). LiCl was added to the cultures 24 hr after TGF-β1 stimulation (regression experiment), and RNA levels were determined by qPCR. Unpaired t test (n=2 independent experiment).

**Figure S5**

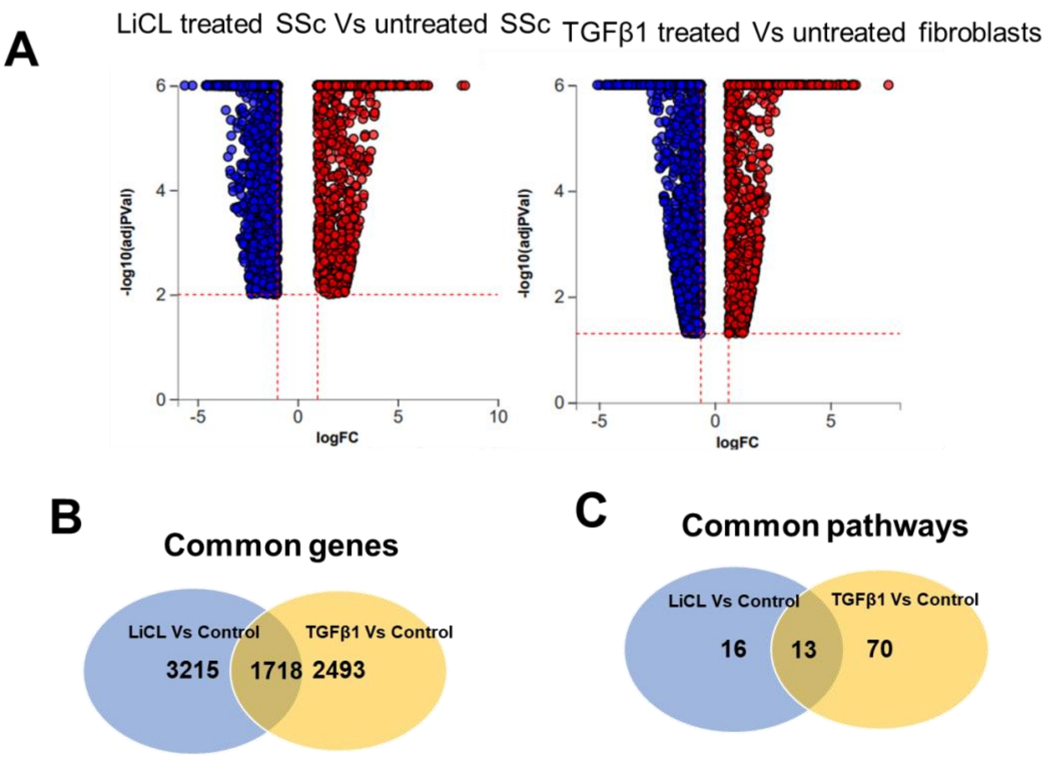

**Supplementary Figure S5. Gene expression changes induced by LiCl and TGF-β1 treatment of healthy and SSc fibroblasts.** (A) Volcano plots representing $\log_2$ fold change (*x* axis) and adjusted *P* value (*y* axis) of total RNA transcripts in LiCl treated SSc fibroblast (left) and TGF-β1 treated healthy skin fibroblasts (right) compared with matched controls. For LiCl treated vs. untreated SSc fibroblast (Total of 4933 genes were differentially regulated, out of which 2831 genes were downregulated, and 2102 genes were upregulated). For TGF-β1 treated vs untreated healthy skin fibroblasts (4211 genes were differentially regulated, out of which 2419 genes were downregulated; 1792 genes were upregulated). The Venn diagrams show common genes (B) and common pathways (C) that were differentially regulated between the two analyses.

### Figure S6

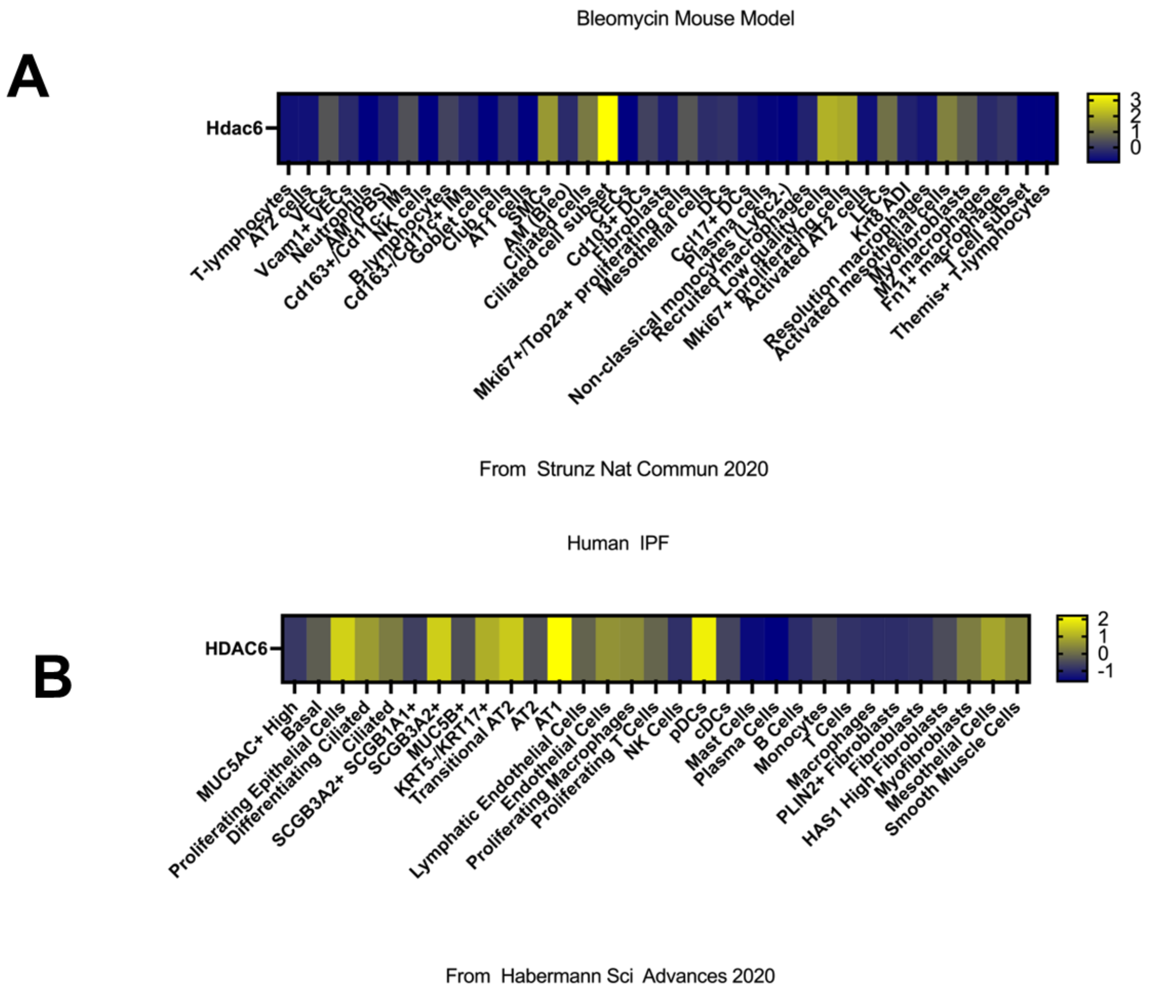

**Supplementary Figure S6: HDAC6 expression in lung fibrosis.**

HDAC6 levels were queried from publicly available single-cell RNAseq database for (A) bleomycin mouse model (Strunz 2020) and (B) human IPF(Habermann 2020).

**Figure S7**

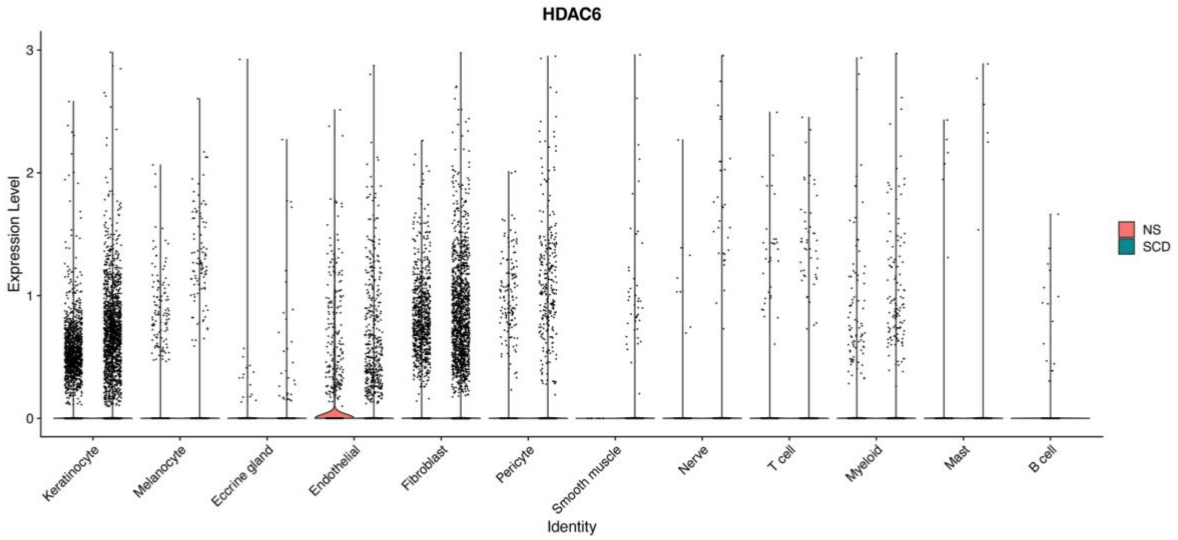

**Supplementary Figure S7: HDAC6 expression SSc skin biopsies.**

Violin plot representing the expression of HDAC6 within the single cell RNAseq data of control (NS, red) vs SSc skin biopsies (SCD, blue). Each dot represents single cell within the data. The position of the dot represents the expression of HDAC6.

**Figure S8**

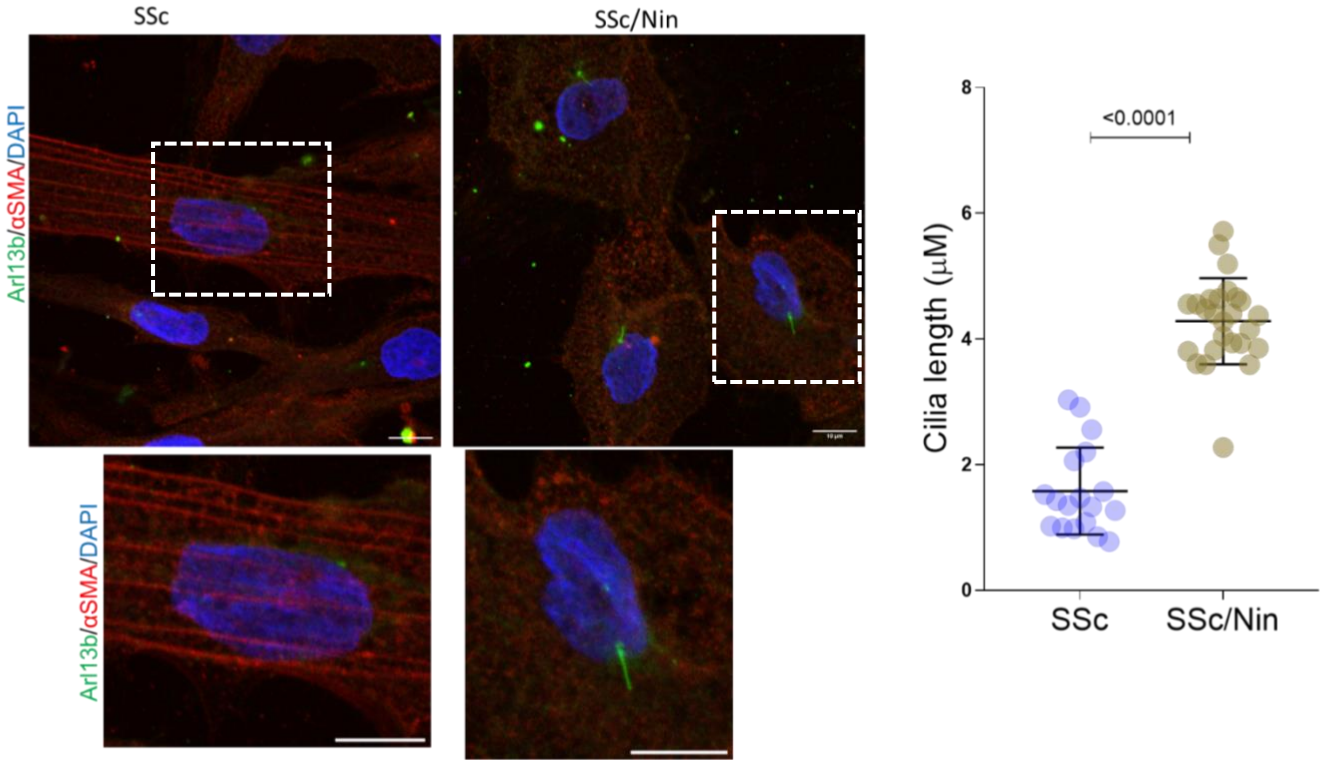

**Supplementary Figure S8: Nintedanib (Nin) treatment increases PC length in SSc skin fibroblast.** Confluent SSc skin fibroblasts (n=2) were serum-starved for 12 hr (1% FBS), followed by treatment of Nintedanib (2 µM; selleckchem) 24 hr. (A) Immunolabelling with Arl13b (green), αSMA (Red). Representative images; bar=10 µm. Magnified insets shown. The graph represents the quantification of PC length (mean ± SD from 9-10 determinations/SSc cell line). Unpaired t test.

### Figure S9

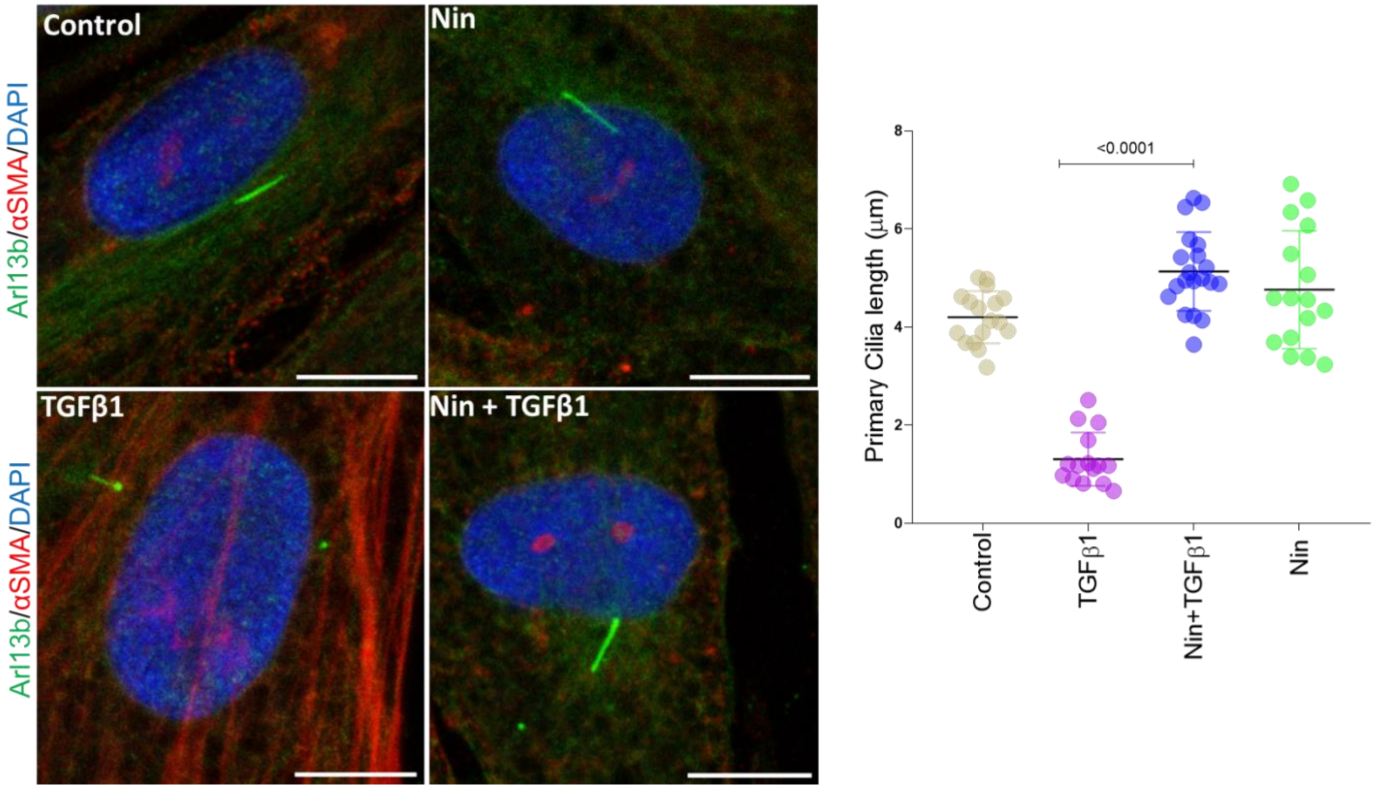

**Supplementary Figure S9: Nintedanib treatment prevents TGF-β1 induced PC shortening in lung fibroblasts (CCL210).** CCL210 lung fibroblasts were 0.1% BSA starved for 24 hr, followed by Nintedanib (Nin) (2μM; selleckchem) pretreatment for 60 min, and treatment with TGF-β1(10 ng/ml) for 24 hr. The graph represents the quantification of PC length (mean ± SD from 15-20 determinations) in the presence and absence TGF-β1. Unpaired t test. The lengths of the PC were measured from the 3D reconstruction of the Z-volumes using ImageJ.

**Figure S10**

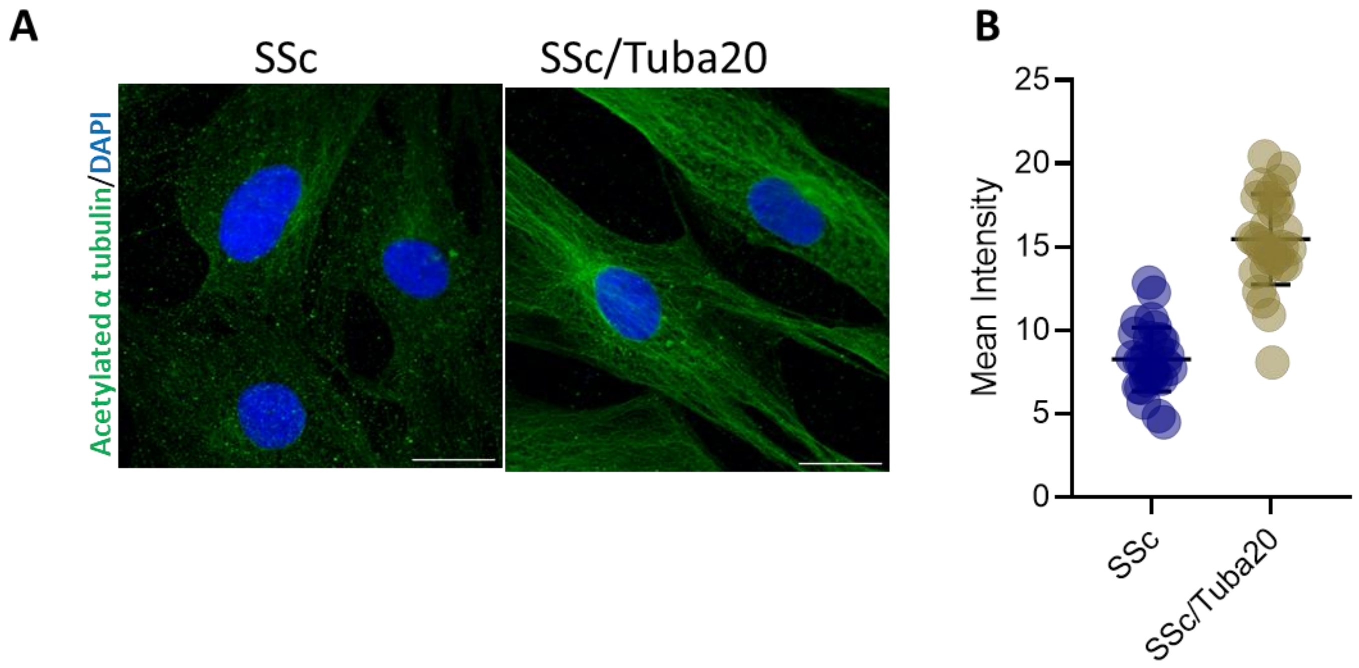

**Supplementary Figure S10: Tubacin treatment increases acetylated α tubulin expression in SSc skin fibroblast.** Confluent SSc skin fibroblasts (n=1) were serum-starved for 12 hr (1% FBS), followed by treatment of tubacin treatment (20µM) for 24 hr. (A) Immunolabelling with acetylated α tubulin (green) antibody. (B) Quantification of acetylated α tubulin mean intensity were measured by ImageJ.
